# Using systems medicine to identify a therapeutic agent with potential for repurposing in Inflammatory Bowel Disease

**DOI:** 10.1101/513838

**Authors:** Katie Lloyd, Stamatia Papoutsopoulou, Emily Smith, Philip Stegmaier, Francois Bergey, Lorna Morris, Madeleine Kittner, Hazel England, Dave Spiller, Mike HR White, Carrie A Duckworth, Barry J Campbell, Vladimir Poroikov, Vitor AP Martins dos Santos, Alexander Kel, Werner Muller, D Mark Pritchard, Chris Probert, Michael D Burkitt, the SysmedIBD consortium

## Abstract

**Objective:** Inflammatory bowel diseases cause significant morbidity and mortality. Aberrant NF-κB signalling is strongly associated with these conditions, and several established drugs influence the NF-κB signalling network to exert their effect. This study aimed to identify drugs which alter NF-κB signalling and may be repositioned for use in inflammatory bowel disease.

**Design:** The SysmedIBD consortium established a novel drug-repurposing pipeline based on a combination of in-silico drug discovery and biological assays targeted at demonstrating an impact on NF-kappaB signalling, and a murine model of IBD.

**Results:** The drug discovery algorithm identified several drugs already established in IBD, including corticosteroids. The highest-ranked drug was the macrolide antibiotic Clarithromycin, which has previously been reported to have anti-inflammatory effects in aseptic conditions.

Clarithromycin’s effects were validated in several experiments: it influenced NF-κB mediated transcription in murine peritoneal macrophages and intestinal enteroids; it suppressed NF-κB protein shuttling in murine reporter enteroids; it suppressed NF-κB (p65) DNA binding in the small intestine of mice exposed to LPS, and it reduced the severity of dextran sulphate sodium-induced colitis in C57BL/6 mice. Clarithromycin also suppressed NF-κB (p65) nuclear translocation in human intestinal enteroids.

**Conclusions:** These findings demonstrate that *in-silico* drug repositioning algorithms can viably be allied to laboratory validation assays in the context of inflammatory bowel disease; and that further clinical assessment of clarithromycin in the management of inflammatory bowel disease is required.

## Introduction

Inflammatory bowel diseases (IBD) affect 0.5-1.0% of people in the Western world and cause substantial morbidity and cost to society^1 2^. The aetiology of IBD is complex. There are aberrant inflammatory responses, leading to mucosal damage and the disease phenotype.

Diverse therapeutic approaches are employed: Many patients with mild ulcerative colitis are treated with mesalazine preparations, which predominantly act topically on the colonic mucosa to suppress inflammation^3^, others require systemic therapy with highly-specific biologic agents targeting inflammatory mediators^4 5^. While these drugs have very different mechanisms of action, they influence host inflammatory responses and either directly or indirectly alter NF-κB signalling.

The NF-κB signalling network is a tightly-controlled, dynamically-regulated signal transduction pathway with several well-described transcription regulatory feedback loops^6^: this orchestrates innate immune responses by regulating transcription through dimers of the five NF-κB proteins (RelA(p65), RelB, NF-κB1(p50), NF-κB2(p52) and c-Rel). Signalling through this network is characterised by the oscillation of NF-κB proteins between the cytoplasm and nucleus; most clearly demonstrated for the RelA(p65) subunit^7^. In addition to changes in the inflammatory milieu, this network also affects several other processes which are dysregulated during chronic gastrointestinal inflammation including cell turnover, DNA damage responses and cell senescence^8^. Several studies have associated aberrant NF-κB signalling with IBD (reviewed in refs^9-11^).

Because of the complexity of NF-κB signalling, targeting specific components of the network has not been a successful drug development strategy to date, mainly because of the ubiquitous nature of NF-κB signalling. The impact of gross inhibition of critical members of the NF-κB signalling cascade has undesirable off-target effects, prompting calls for highly specific inhibitors which target components of the network in precisely-selected cell types^12^. This approach is useful in certain circumstances such as multiple myeloma, where targeting a specific cancer cell lineage is desirable and may be achievable. In contrast, for complex benign inflammatory diseases, including IBD, this strategy is less likely to be successful due to challenges in identifying a specific target cell population.

We propose an alternative strategy, to ‘nudge’ an individual’s NF-κB signalling network towards an idealised healthy phenotype. This approach may offer a more pragmatic way of targeting aberrant NF-κB signalling, without the limitations of selective tissue targeting or gross pathway inhibition.

To investigate this in the context of inflammatory bowel disease, we used a combination of novel bioinformatic analyses and laboratory studies to identify agents likely to impact NF-κB signalling and inflammatory bowel disease. We then validated the efficacy of the most highly ranked agent, both in terms of inhibiting NF-κB activity, and modulating an *in-vivo* model of colitis.

## Methods

### Pathway analysis for master regulator search

Molecules that regulate the expression of differentially expressed genes through control of the activity of NF-κB sub-units were defined as master regulators and were identified by applying a key-node analysis algorithm^36^, using the TRANSPATH^®^ database of gene regulatory and signal transduction pathways^37^.

Key-nodes were prioritised based on the weighted ratio between the number of molecules from the input set that could be reached from the key-node in ≤10 steps and the total number of reachable nodes. The higher the score, the greater the chance that the key-node plays a master regulatory role.

### In-silico drug discovery

Relationships between the chemical structure of compounds and their biological activities were analysed using large-scale prediction of activity spectra^38 39^ (http://genexplain.com/pass/) to discover potential new drugs for IBD treatment.

The PASS algorithm is based on Bayesian estimates of probabilities of molecules belonging to the classes of active and inactive compounds^40^. The predicted activity spectrum is presented in PASS by the list of activities with probabilities “to be active” Pa and “to be inactive” Pi calculated for each activity.

For each compound of the tested library, a cumulative score was computed across the 46 PASS activities selected in the previous analysis.

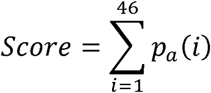

here *p*_*a*_*(i)* is the probability for the given compound to be active for each activity *i*.

For each compound tested, the cumulative score was required to be higher than 3.0, and the Pa for the PASS activity “Transcription factor NF-κB inhibitor” was positive.

### Patient recruitment and ethics

Samples used for the generation of human enteroids were donated by patients without evidence of IBD attending for colonoscopy at the Royal Liverpool and Broadgreen University Hospitals NHS Trust. Samples were collected following written informed consent and favourable ethical opinion from North West-Liverpool East Research Ethics Committee (15/NW/0045).

### Animal maintenance and welfare

All animal breeding, maintenance and procedures were performed under a UK Home Office licence, with local Animal Welfare and Ethics Review Board approval. Transgenic animals were maintained at the University of Manchester. Wild-type mice were purchased from Charles River (Margate, UK) and maintained either at the University of Manchester or University of Liverpool in SPF facilities with access to standard chow and drinking water *ad-libitum* unless otherwise specified. Standard 12-hour light/dark cycles were used; standard temperature and humidity levels were maintained.

### Transgenic mouse strains

#### Human TNFα luciferase mice (hTNF.LucBAC)

This line has been engineered to have a bacterial artificial chromosome (BAC) expressing luciferase under the regulation of the entire human TNF promoter. Primary cultures established from this mouse enable direct measurement of TNF promoter activity in a non-transformed *ex-vivo* system^22^.

#### Human p65-DsRedxp/IκBalpha-eGFP mouse (p65-DsRedxp/IκBalpha-eGFP)

This line expresses fusion proteins of NF-κB(p65) and the *Discosoma* red fluorescent protein – Express (DsRedxp) under the regulation of the native human p65 promoter, and IκBalpha and enhanced green fluorescent protein (eGFP), regulated by the human IκBalpha promoter^41^. Primary cultures established from this mouse enable direct visualisation of these fusion proteins.

#### Peritoneal macrophage isolation, in vitro culture and luciferase assay

Resident peritoneal macrophages were obtained from mice by standard methods. Cells were stimulated with either 10ng/mL LPS (*Salmonella enterica* serovar Minnesota R595; Calbiochem), TNF (R&D Systems) at 40 ng/mL, muramyl dipeptide (MDP, Invivogen) at 10μg/mL or Flagellin (Novus Bio) at 500ng/ml. Luciferase activity was measured over time in a CO_2_ luminometer (Lumistar Omega, BMG Labtech).

#### Flow Cytometry

Peritoneal macrophages were stimulated with 1μg/ml LPS for 20min. Fixed cells were stained for phosphor-p65 (PE Mouse anti-NF-κB p65 (pS529), BD Bioscience, clone K10-895.12.50) and analysed by flow cytometry using an LSRII Cytometer (BD). Data analyses were performed with FlowJo887 software (Tree Star).

#### Primary Epithelial cell culture

Enteroids were generated from human and murine tissue using modifications of established protocols. Tissue samples were disaggregated by calcium chelation in EDTA followed by mechanical disaggregation in a sucrose/sorbitol solution. Crypt pellets were resuspended in Matrigel (Corning, UK) and were maintained in media containing a combination of recombinant growth factors and growth factor containing conditioned media (supplemental methods). Enteroids were passaged at least once prior to experiments.

#### Enteroid immunohistochemistry

Enteroids were pre-treated with 10μM clarithromycin or 1% v/v DMSO (vehicle) for 30mins. Following pre-treatment, 100ng/mL recombinant human TNF (Peprotech) was applied for a further 30mins. Fixed enteroids were transferred into Richard-Allan™ Histogel™ (Thermofisher) and processed for histology. Immunohistochemical analysis was performed to visualise p65 (Cell Signaling Technologies, #8242).

#### Murine intestinal organoid confocal microscopy

Proximal enteroids from p65-DsRedxp/IκBalpha-eGFP dual-reporter mice were passaged into glass-bottom dishes (Thermo Scientific™ Nunc™ Glass Bottom Dishes) in phenol red-free complete media. Images were taken for 6-8 enteroids per dish using a Zeiss Laser Scanning Microscope (LSM880) with a C-Apochromat 40x/1.2 W Korr FCS M27 objective. Enteroids were imaged for 30mins before treatment with 10μM clarithromycin or 1% v/v DMSO vehicle. Thirty minutes later, cultures were stimulated with 100ng/ml TNF and imaged for a further 3hrs. Images and videos were processed using Zen 2011 software and CellTracker (Warwick Systems Biology Centre).

#### LPS-induced NF-κB activation in mice

Groups of adult (8-10-week-old) male C57BL/6 mice received 50mg/kg clarithromycin i.p, or vehicle daily for three days. Following a 3-day washout period, either 50mg/kg clarithromycin i.p., or 0.9% w/v saline vehicle was administered, the next day either 50mg/kg clarithromycin i.p. or vehicle was followed by either 0.125mg/kg ultrapure LPS from *E. coli* K12 (Invivogen) i.p. or 0.9% w/v saline vehicle. Animals were culled 90 minutes after LPS/vehicle administration; small intestinal mucosal scrapes were prepared in RIPA buffer with protease inhibitors (Sigma, Gillingham, UK). NF-κB p65 DNA binding was quantified using a TransAM DNA-binding ELISA (ActiveMotif, La Hulpe, Belgium).

#### Dextran sulphate sodium-induced colitis

Groups of adult (8-10-week-old) male C57BL/6 mice were administered either 10 mg/kg clarithromycin, or vehicle by orogastric gavage daily for four days. Following a washout period, animals received 2.5% DSS in drinking water, for five days. Animals recovered for a further three days. From the start of DSS treatment to termination of the experiment either 10 mg/kg clarithromycin or normal saline vehicle was administered by daily orogastric gavage. Tissue samples were harvested and prepared for histological analysis, including quantitative histology, as previously described^42^.

#### Statistical analysis of laboratory work

Statistical analyses were performed using GraphPad Prism v.7.0 software. Specific statistical tests are annotated in the text and figure legends. Reported *p-*values are two-tailed. Normality testing was performed with the D’Agostino & Pearson test.

## Results

To predict drugs that may influence disrupted NF-κB signalling in IBD, a drug-discovery strategy combining data from multiple sources (archived ChIP-seq analyses; natural language text-mining of published abstracts; data from IBD GWAS analyses; and known IBD biomarkers and drug targets curated in the HumanPSD database) was developed (Figure 1).

**Figure 1:**
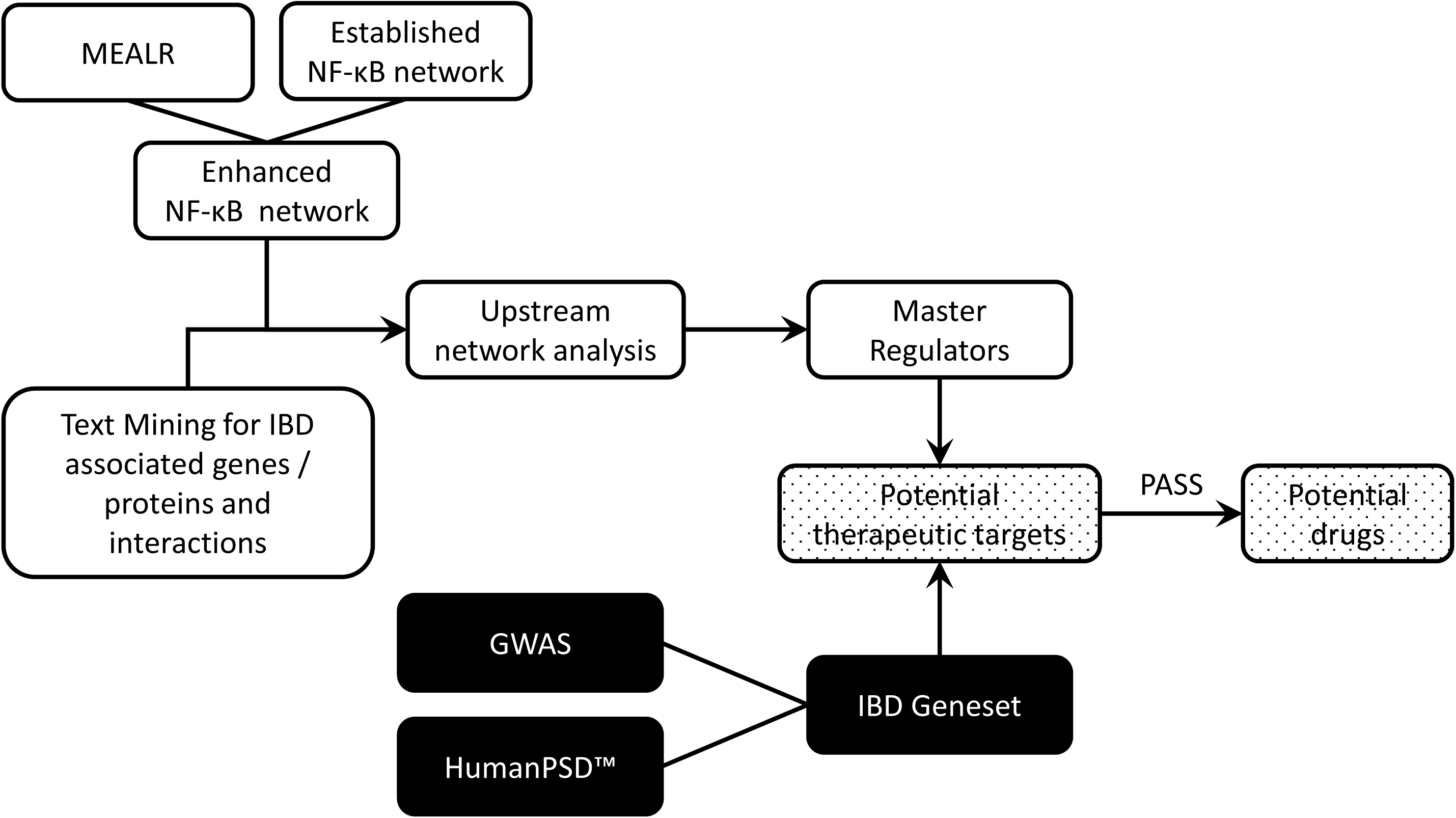
Schematic representation of the bioinformatic approach to identifying drugs with the potential to modulae IBD and NF-κB signalling.

### Developing an enhanced NF-κB signalling network

Signalling molecules comprising (transcription) regulatory feedback loops present promising drug targets because they can influence the dynamics of a signalling pathway of interest, including the TNF-alpha/NF-κB pathway. To augment existing knowledge about NF-κB signalling, we developed an analysis workflow to identify genes encoding potential components of transcription regulatory feedback loops combining known NF-κB-involving signalling pathways, ChIP-seq assay based NF-κB/RelA-bound genomic regions from the ENCODE project (GEO series GSE31477), and a newly-developed method to find combinations of enriched DNA-sequence motifs (Motif Enrichment Analysis by Logistic Regression (MEALR)) (see supplementary materials for details). Combinations of prioritised motifs tend to coincide with transcription factors that are known to cooperate. Our analysis found 24 transcription factors that appear to cooperate in NF-κB signalling within the genomic regions reported by ENCODE (Table 1) as well as 90 potential NF-κB/RelA-target genes that encode components of known pathways. (Table 2). The results were used to annotate the TRANSPATH^®^ database of mammalian signal transduction and metabolic pathways.

**Table 1:**
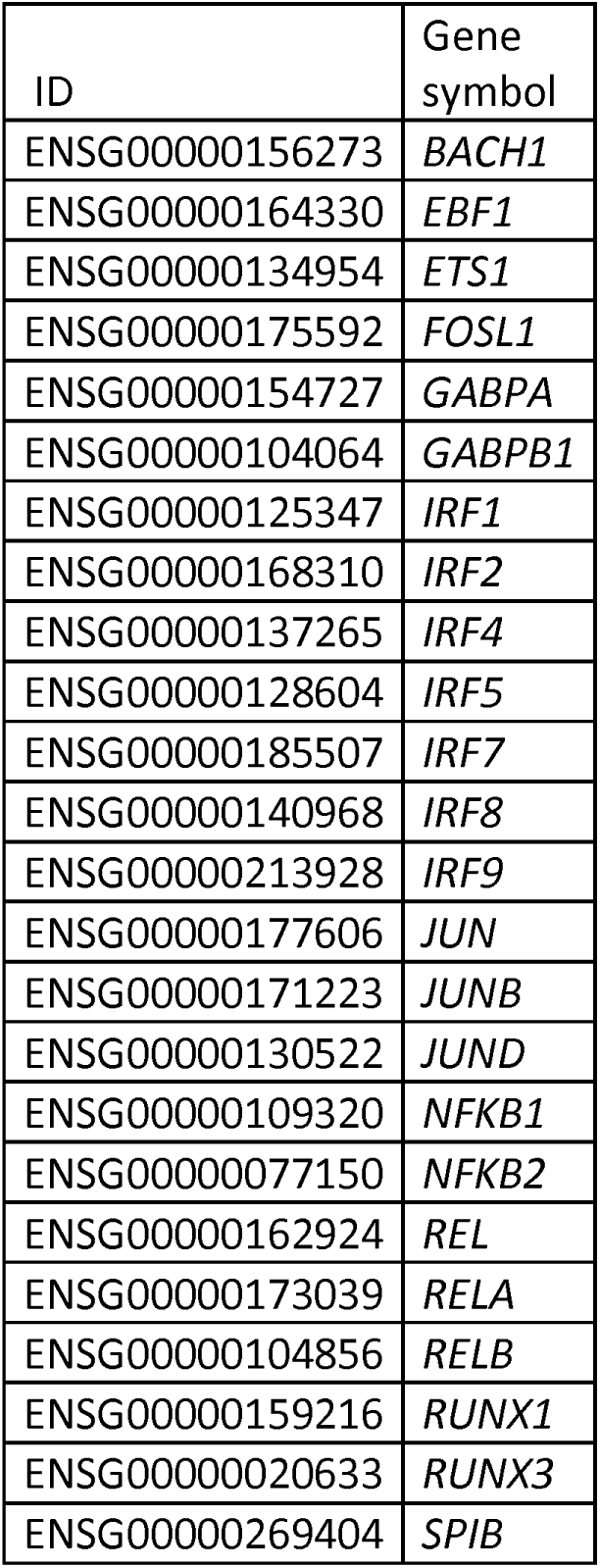
Transcription factors predicted to contribute to NF-κB regulatory cascades by MEALR analysis

**Table 2:**
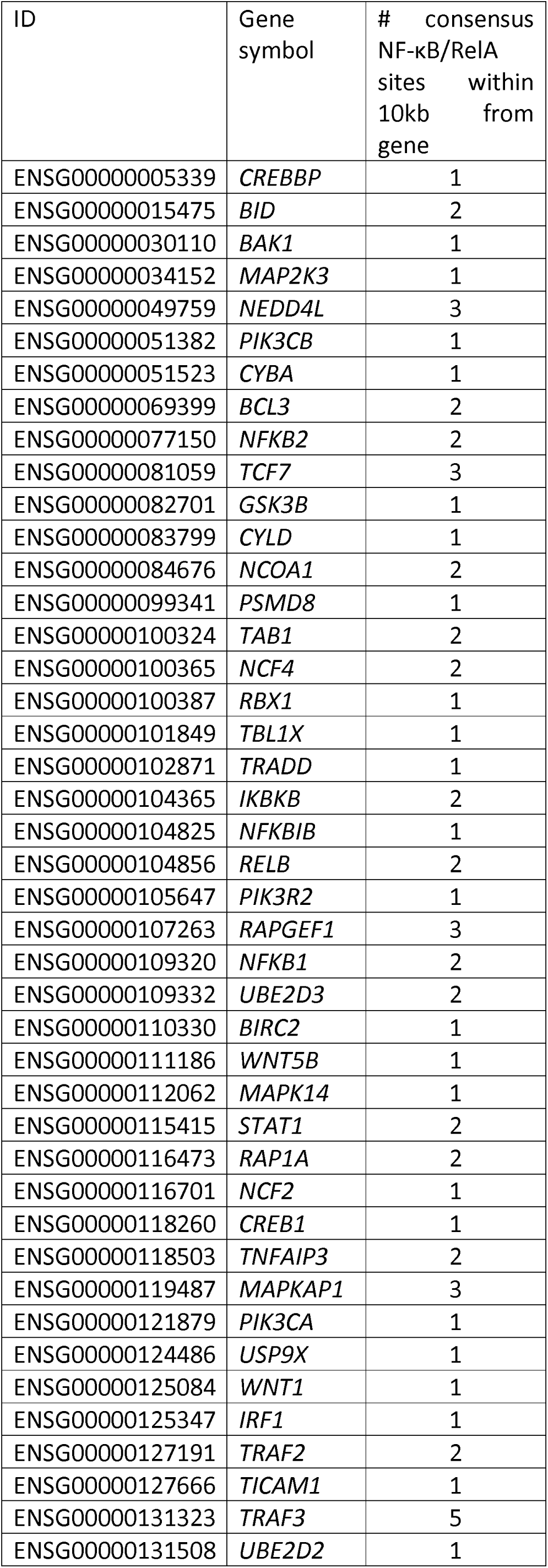
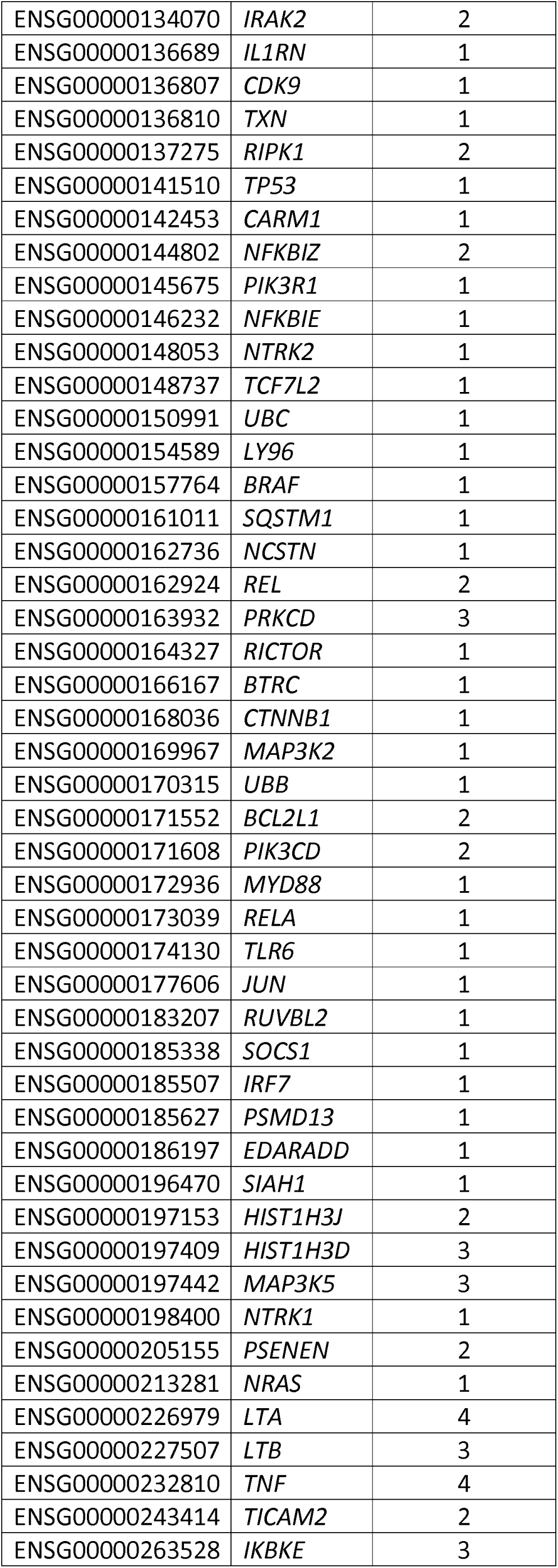
NF-κB target genes identified by MEALR to be involved in regulatory cascades

| ID | Gene symbol | # consensus NF-κB/RelA sites within 10kb from gene |
| --- | --- | --- |
| ENSG00000005339 | *CREBBP* | 1 |
| ENSG00000015475 | *BID* | 2 |
| ENSG00000030110 | *BAK1* | 1 |
| ENSG00000034152 | *MAP2K3* | 1 |
| ENSG00000049759 | *NEDD4L* | 3 |
| ENSG00000051382 | *PIK3CB* | 1 |
| ENSG00000051523 | *CYBA* | 1 |
| ENSG00000069399 | *BCL3* | 2 |
| ENSG00000077150 | *NFKB2* | 2 |
| ENSG00000081059 | *TCF7* | 3 |
| ENSG00000082701 | *GSK3B* | 1 |
| ENSG00000083799 | *CYLD* | 1 |
| ENSG00000084676 | *NCOA1* | 2 |
| ENSG00000099341 | *PSMD8* | 1 |
| ENSG00000100324 | *TAB1* | 2 |
| ENSG00000100365 | *NCF4* | 2 |
| ENSG00000100387 | *RBX1* | 1 |
| ENSG00000101849 | *TBL1X* | 1 |
| ENSG00000102871 | *TRADD* | 1 |
| ENSG00000104365 | *IKBKB* | 2 |
| ENSG00000104825 | *NFKBIB* | 1 |
| ENSG00000104856 | *RELB* | 2 |
| ENSG00000105647 | *PIK3R2* | 1 |
| ENSG00000107263 | *RAPGEF1* | 3 |
| ENSG00000109320 | *NFKB1* | 2 |
| ENSG00000109332 | *UBE2D3* | 2 |
| ENSG00000110330 | *BIRC2* | 1 |
| ENSG00000111186 | *WNT5B* | 1 |
| ENSG00000112062 | *MAPK14* | 1 |
| ENSG00000115415 | *STAT1* | 2 |
| ENSG00000116473 | *RAP1A* | 2 |
| ENSG00000116701 | *NCF2* | 1 |
| ENSG00000118260 | *CREB1* | 1 |
| ENSG00000118503 | *TNFAIP3* | 2 |
| ENSG00000119487 | *MAPKAP1* | 3 |
| ENSG00000121879 | *PIK3CA* | 1 |
| ENSG00000124486 | *USP9X* | 1 |
| ENSG00000125084 | *WNT1* | 1 |
| ENSG00000125347 | *IRF1* | 1 |
| ENSG00000127191 | *TRAF2* | 2 |
| ENSG00000127666 | *TICAM1* | 1 |
| ENSG00000131323 | *TRAF3* | 5 |
| ENSG00000131508 | *UBE2D2* | 1 |
| ENSG00000134070 | *IRAK2* | 2 |
| ENSG00000136689 | *IL1RN* | 1 |
| ENSG00000136807 | *CDK9* | 1 |
| ENSG00000136810 | *TXN* | 1 |
| ENSG00000137275 | *RIPK1* | 2 |
| ENSG00000141510 | *TP53* | 1 |
| ENSG00000142453 | *CARM1* | 1 |
| ENSG00000144802 | *NFKBIZ* | 2 |
| ENSG00000145675 | *PIK3R1* | 1 |
| ENSG00000146232 | *NFKBIE* | 1 |
| ENSG00000148053 | *NTRK2* | 1 |
| ENSG00000148737 | *TCF7L2* | 1 |
| ENSG00000150991 | *UBC* | 1 |
| ENSG00000154589 | *LY96* | 1 |
| ENSG00000157764 | *BRAF* | 1 |
| ENSG00000161011 | *SQSTM1* | 1 |
| ENSG00000162736 | *NCSTN* | 1 |
| ENSG00000162924 | *REL* | 2 |
| ENSG00000163932 | *PRKCD* | 3 |
| ENSG00000164327 | *RICTOR* | 1 |
| ENSG00000166167 | *BTRC* | 1 |
| ENSG00000168036 | *CTNNB1* | 1 |
| ENSG00000169967 | *MAP3K2* | 1 |
| ENSG00000170315 | *UBB* | 1 |
| ENSG00000171552 | *BCL2L1* | 2 |
| ENSG00000171608 | *PIK3CD* | 2 |
| ENSG00000172936 | *MYD88* | 1 |
| ENSG00000173039 | *RELA* | 1 |
| ENSG00000174130 | *TLR6* | 1 |
| ENSG00000177606 | *JUN* | 1 |
| ENSG00000183207 | *RUVBL2* | 1 |
| ENSG00000185338 | *SOCS1* | 1 |
| ENSG00000185507 | *IRF7* | 1 |
| ENSG00000185627 | *PSMD13* | 1 |
| ENSG00000186197 | *EDARADD* | 1 |
| ENSG00000196470 | *SIAH1* | 1 |
| ENSG00000197153 | *HIST1H3J* | 2 |
| ENSG00000197409 | *HIST1H3D* | 3 |
| ENSG00000197442 | *MAP3K5* | 3 |
| ENSG00000198400 | *NTRK1* | 1 |
| ENSG00000205155 | *PSENEN* | 2 |
| ENSG00000213281 | *NRAS* | 1 |
| ENSG00000226979 | *LTA* | 4 |
| ENSG00000227507 | *LTB* | 3 |
| ENSG00000232810 | *TNF* | 4 |
| ENSG00000243414 | *TICAM2* | 2 |
| ENSG00000263528 | *IKBKE* | 3 |

### Text mining to establish context proteins and genes for up-stream network analyses

1000 relevant abstracts were retrieved using the MedlineRanker tool^13^ from the PubMed database using the MeSH terms “Inflammatory bowel diseases” and “NF-κB”. Protein-protein interactions were identified in these abstracts using PESCADOR^14^. PESCADOR detects genes, proteins and their interactions, and rates them based on co-occurrences in an abstract. 827 interactions common for both *Homo sapiens* and *Mus musculus*, 2 for *Mus musculus* alone, and 26 uniquely for *Homo sapiens* were extracted (Supplementary Table S1). After querying the TRANSPATH^®^ database for known direct interactions, 127 novel co-occurrences of genes or proteins in the abstracts analysed. This table of interactions was used to provide context to subsequent up-stream network analysis to impute the key regulatory nodes for NF-κB signalling in IBD.

### Identifying master regulators of the NF-κB signalling network

An upstream search of the various molecular components of NF-κB complex including: NF-κB1-isoform1, NF-κB1-isoform2, NF-κB2-isoform4, NF-κB2-p100, NF-κB2-p49, RelA-p35, RelA-p65-delta, RelA-p65-delta2, RelA-p65-isoform1, RelA-p65-isoform4, RelB, c-Rel was performed. The network search extended to a maximal radius of 10 steps upstream of the NF-κB components and used a false discovery rate (FDR) cut-off of 0.05 and Z-score (reflecting how specific each master regulator was) cut-off of 1.0. The list of interacting proteins obtained by text-mining was used to provide “Context proteins” for the master-regulator acquisition algorithm^15^. This analysis revealed 325 controlling nodes predicted to exert signalling activity for the NF-κB components (Table S2).

### Developing a list of IBD associated genes and potential therapeutic targets

A list of IBD associated-genes and potential therapeutic targets was established by retrieving information from the HumanPSD database including known IBD biomarkers and drug targets, and two lists of IBD-related genes from genome-wide association studies, one focused on revealing genes of IBD prognosis^16^, and another on IBD susceptibility^17^. 159 IBD-targets were identified and summarised with an indication of the source of evidence about their relevance to IBD (Table S3).

Finally, to produce a list of proposed therapeutic targets the list of IBD-targets was intersected with the 325 controlling nodes to obtain 62 candidates (defined as IBD key-nodes, Table S4) that represent genes predicted to influence IBD associated NF-κB regulation, which had also independently been identified as IBD associated genes or potential therapeutic targets.

### Predicting drugs that may influence the IBD key-nodes

The PASS software package was used to predict drugs which may interact with IBD key-nodes: this software predicts the ability of a chemical structure to interact and influence the activity of defined molecular targets and biological activities.

The 62 key-nodes were translated into 46 PASS activities (Table S5). To identify compounds with potential for clinical re-positioning, the Top 200 drugs library was analysed to predict the probability that these established, and licenced agents may interfere with each PASS activity.

A cumulative score was calculated across all 46 PASS activities for each drug. We also required that the drugs were predicted to be active against the PASS activity “Transcription factor NF κB inhibitor”.

29 compounds achieved these criteria (Table 3); importantly, this table included several corticosteroids already used to treat IBD, supporting the validity of the discovery strategy.

**Table 3:** Drugs predicted to influence IBD.

| Ranking | Drug |
| --- | --- |
| 1 | Clarithromycin |
| 2 | Budesonide |
| 3 | Digoxin |
| 4 | Dexamethasone |
| 5 | Triamcinolone acetonide |
| 6 | Methylprednisolone |
| 7 | Prednisone |
| 8 | Azithromycin |
| 9 | Medroxyprogesterone |
| 10 | Estradiol |
| 11 | Nystatin |
| 12 | Progesterone |
| 13 | Albuterol |
| 14 | Ibuprofen |
| 15 | Dicyclomine |
| 16 | Naproxen |
| 17 | Nabumetone |
| 18 | Propranolol |
| 19 | Metoprolol |
| 20 | Atenolol |
| 21 | Acetaminophen |
| 22 | Phentermine |
| 23 | Divalproex |
| 24 | Gabapentin |
| 25 | Diclofenac |
| 26 | Lisdexamfetamine |
| 27 | Pregabalin |
| 28 | Methocarbamol |
| 29 | Metformin |

The highest-ranked drug was a macrolide antibiotic: clarithromycin. This agent is of particular interest because macrolides have an established role in treating aseptic inflammatory conditions including chronic rhinosinusitis^18^ and panbronchiolitis^19^ and because clarithromycin has previously been trialled in IBD with divergent outcomes^20 21^. We, therefore, considered it to be an excellent candidate for strategic repurposing and have used clarithromycin as a paradigm molecule for the development of a mechanism-led drug validation pathway.

### Clarithromycin suppresses stimulus-induced luciferase activity in primary cell cultures

To validate the efficacy of clarithromycin as an inhibitor of NF-κB mediated transcription, primary cultures from the hTNF.LucBAC mouse, a transgenic line that expresses firefly luciferase under the control of the entire human *TNF* promoter^22^, were used.

Luciferase activity was triggered by ligand binding to the TNF receptor and pattern recognition receptors including TLR4 (lipopolysaccharide), TLR5 (flagellin) and NOD2 (muramyl dipeptide, MDP) (Figures 2A-D) in peritoneal macrophages harvested from this mouse. When cells were pre-treated for 30 minutes with 10μM or 100μM clarithromycin, a dose-dependent reduction in luciferase activity was observed, independent of the stimulus applied.

**Figure 2.**
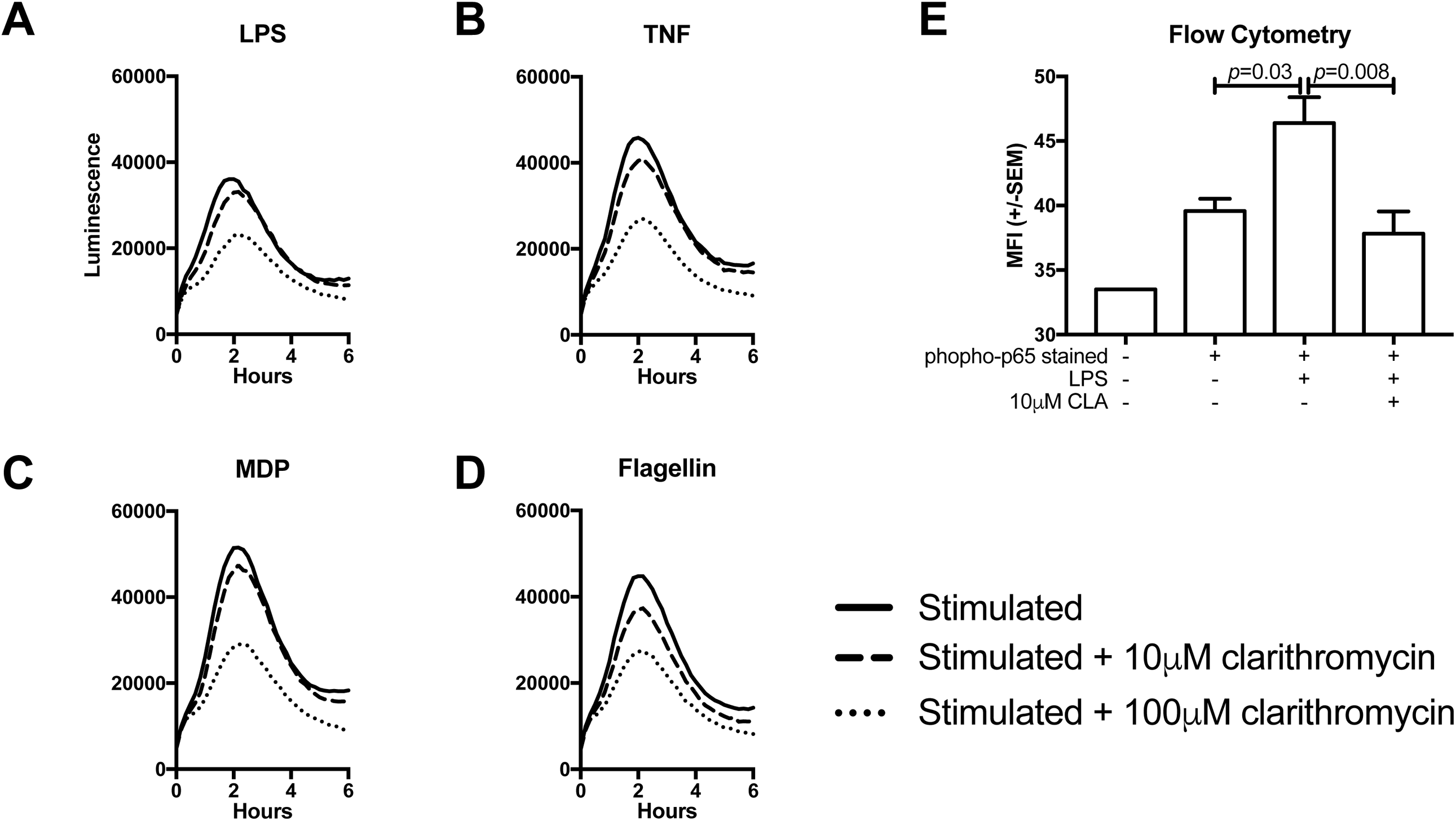
**A-D:** Luciferase activation curves for peritoneal macrophages extracted from HTNF.LucBAC mice stimulated with 10ng/mL LPS (**A**), 40 ng/mL TNF (**B**), 10μg/mL MDP (**C**) or 500ng/ml flagellin (**D**). Solid lines indicate cells response of pre-treated with drug vehicle (DMSO), dashed and dotted lines responses generated from cells pre-treated with 10μM or 100 μM clarithromycin, respectively. **E:** Median fluorescence intensity for anti-phopsho-p65 antibody stained peritoneal macrophages from C57BL/6 mice either unstimulated or stimulated with LPS and pre-treated with 10μM clarithromycin or DMSO vehicle. N=4 mice. Statistically significant differences tested by 1-way ANOVA and Dunnett’s post hoc test.

The effect of clarithromycin on NF-κB activation in murine peritoneal macrophages was confirmed by isolating peritoneal macrophages from 4 WT mice, and treating them with 10μM clarithromycin or vehicle before stimulation with 1μg/ml LPS. Cells were fixed, stained for phosphorylated-p65 and quantified by flow cytometry. Median fluorescence intensity (MFI) increased on stimulation (*p*=0.03, ANOVA); co-administration of LPS and clarithromycin suppressed this response (*p*=0.008, Figure 2E).

Small intestinal organoids (enteroids) were established from hTNF.LucBAC mice to determine whether clarithromycin could also alter NF-κB responses in gastrointestinal epithelial cell cultures. Luciferase activity was detectable in these cultures in response to 100ng/mL TNF administration (Figure 3A and B). Luciferase activity was significantly suppressed by pre-treatment for 30 minutes with either 1μM (*p*=0.014) or 10μM (*p*=0.001) clarithromycin (Figure 3B and C).

**Figure 3:**
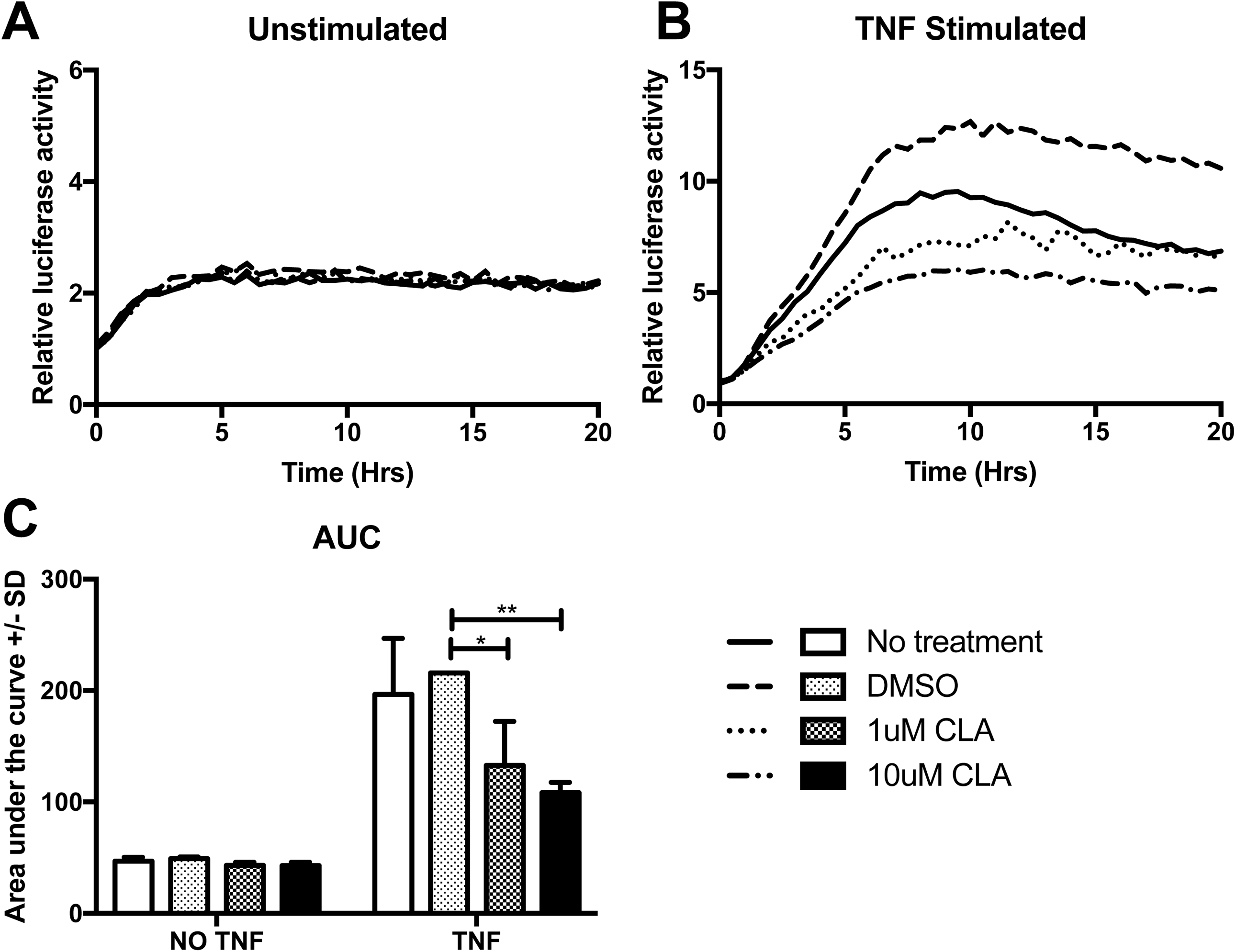
Representative luciferase activation curves for enteroids derived from HTNF.LucBAC mice either unstimulated (**A**) or stimulated with 100ng/mL TNF (**B**), and without pre-treatment (solid line) or 30 min pre-treatment with, DMSO vehicle (dashed line), 1μM clarithromycin (dotted line) or 10μM clarithromycin (dotted and dashed line). **C:** Area under the curve (AUC) calculations for the same experiment. N=3. Statistically significant differences tested by 2-way ANOVA and Dunnett’s posthoc test.

### Clarithromycin suppresses TNF-induced NF-κB(p65) shuttling in enteroids

To assess whether clarithromycin influenced NF-κB protein shuttling dynamics, enteroid cultures from reporter mice expressing p65-DsRedxp/IκBalpha-eGFP were established. This mouse expresses human p65-DsRedxp and human IκBalpha-eGFP fusion proteins.

Administration of 100ng/mL TNF-induced synchronised p65 translocation to the nucleus of cells within enteroids with a periodicity of approximately 50 minutes. This was observed as a highly damped shuttling response, with a single wave of synchronised nuclear translocation, followed by a second wave of partially synchronised translocation, after which further nuclear localisation occurred in an apparently stochastic fashion (Figure 4A and B and supplementary video). When cultures were pre-treated with 10µM clarithromycin for 30 minutes, p65-DsRedxp nuclear translocation was markedly suppressed.

**Figure 4.**
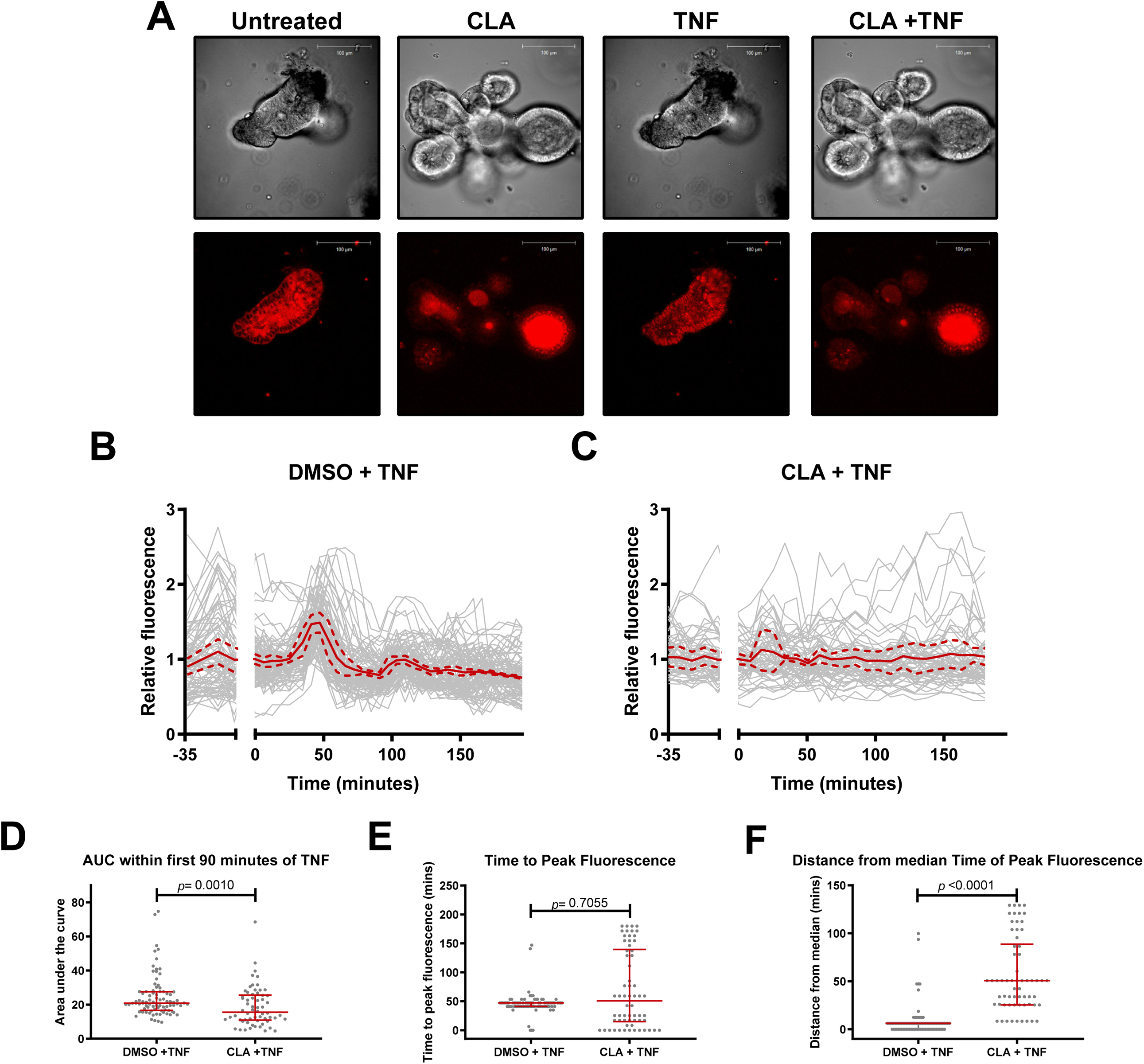
**A:** Representative images of bright field (upper panels) and red channel (lower panels) images of enteroids derived from p65-DsRedxp/IκBalpha-eGFP mice, either untreated, treated with clarithromycin alone, treated with 100ng/mL TNF alone, or pre-treated with 10μM clarithromycin and subsequently stimulated with TNF. **B and C:** Relative nuclear red fluorescence curves for individual cells (grey lines), mean (solid red line) and 1 SD above and below the mean (dashed red lines) over time. Cells pre-treated with DMSO vehicle **(B)** or clarithromycin **(C)** at time -35 min and stimulated with TNF at time 0 min. **D:** Area under the curve calculations for individual cells between time 0 and time 90 min for panels B and C. Statistically significant differences tested by Mann-Whitney U test. **E:** Time to peak fluorescence, post-TNF stimulation for individual cells, statistically significant differences tested by Mann-Whitney U test. **F:** Distance of individual cells peak fluorescence from the median value, statistically significant differences tested by Mann-Whitney U test. N=88 vehicle pre-treated, 62 clarithromycin pre-treated cells for panels D-F, lines represent median and IQR for these panels.

To quantify these events more precisely the mean area under the curve (AUC) during the first oscillatory wave (Figure 4D) was calculated, and demonstrated decreased nuclear intensity of p65-DsRed fluorescence in clarithromycin pre-treated enteroids (*p*=0.005). The time to peak nuclear fluorescence after 100ng/mL TNF administration was also quantified (figure 4E). In DMSO vehicle-treated cells, peak fluorescence occurred at a median time of 47.2 (IQR 41.0-47.2) minutes. This was not significantly different for clarithromycin treated enteroids, but there was significantly more cell-to-cell variability observed in clarithromycin treated than vehicle-treated cells (*p*<0.0001), demonstrating the loss of a synchronisation (Figures 4C and 4F).

### Clarithromycin suppresses LPS-induced NF-κB (p65) DNA binding in-vivo

To determine whether the effects of clarithromycin on stimulus-induced NF-κB activity observed *in-vitro* also occurred *in-vivo*, 0.125mg/kg LPS was administered to groups of six C57BL/6 male mice by intraperitoneal injection, either with or without clarithromycin co-administration. This stimulus induces small intestinal epithelial cell shedding, regulated by both NF-κB1 and NF-κB2 signalling pathways^23^.

LPS administration induced a 2-fold increase in p65 DNA binding, compared to saline vehicle control (*p*<0.001, 1-way ANOVA and Dunnett’s posthoc test); pre-treatment with clarithromycin suppressed this effect by approximately 51% (*p*=0.007, Figure 5A).

**Figure 5.**
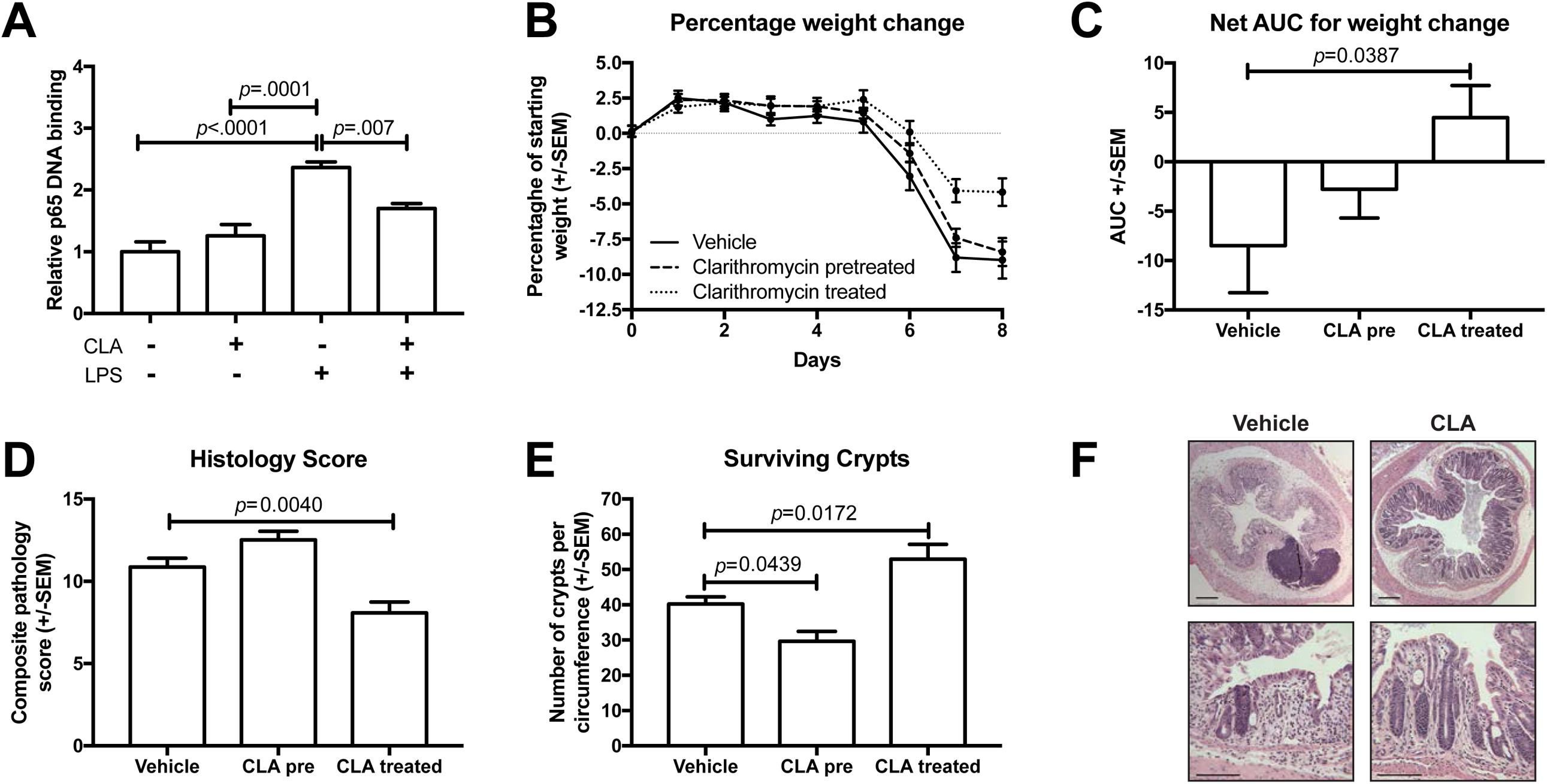
**A:** Relative p65 DNA binding activity in whole-cell lysates from proximal small intestine of C57BL/6 mice pretreated for three days with 50mg/kg clarithromycin or saline vehicle, and subsequently injected i.p. with 0.125mg/kg LPS or vehicle. N=5-6. **B-F:** Effect of clarithromycin on outcomes of DSS colitis in C57BL/6 mice. **B:** Weight loss plotted over time. **C:** Area under the curve analysis of weight loss. **D:** Histology severity scores. **E:** Number of surviving crypts per colonic circumference. **F:** Representative photomicrographs of the colonic mucosa of DSS treated mice co-treated with saline vehicle or 10mg/kg clarithromycin by oro-gastric gavage. N=9-10. Statistically significant differences tested by 1-way ANOVA and Dunnett’s post hoc test in all panels.

### Clarithromycin suppresses DSS-induced colitis

To determine whether clarithromycin affected murine colitis *in-vivo*, clarithromycin or vehicle were administered to mice receiving DSS to induce colitis. Animals received clarithromycin or vehicle daily for four days by oro-gastric gavage, after a four-day washout period 2.5% w/v DSS in drinking water *ad-libitum* was commenced for five days, followed by recovery for a further three days.

Mice co-administered DSS and clarithromycin lost significantly less weight than other groups (*p*=0.039, 1-way ANOVA and Dunnett’s posthoc test, Figures 5B and 5C) and had lower compound histology scores (*p*=0.004, Figures 5D and 5F) and a higher number of surviving colonic crypts than mice treated with vehicle (*p*=0.017, Figure 5E); this suggests that clarithromycin at least partially ameliorates this model of colitis.

### Clarithromycin suppresses TNF-induced NF-κB (p65) nuclear localisation in human enteroids

To determine whether the effects identified in murine primary culture and in-vivo experiments were also relevant to humans, passaged human ileal organoids from individuals with no evidence of IBD were pre-treated with 10µM clarithromycin or DMSO vehicle for 30 min before stimulation with 100ng/ml TNF. Paraformaldehyde fixed cultures were immunostained for p65, and the percentage of cells expressing nuclear-localised p65 was quantified. In untreated human enteroids, 0.6% (SEM 0.37) of cells demonstrated p65 nuclear localisation, administration of TNF induced a 57-fold increase in cells expressing nuclear p65 (33%, +/- 3.2 SEM, *p*<0.0001, 1-way ANOVA and Dunnett’s posthoc test, N=6 per group). Pre-treatment with clarithromycin suppressed TNF-induced nuclear localisation of p65 to a level comparable with DMSO vehicle-treated enteroids. (1%, +/- 0.36 SEM, *p*<0.0001, Figure 6).

**Figure 6.**
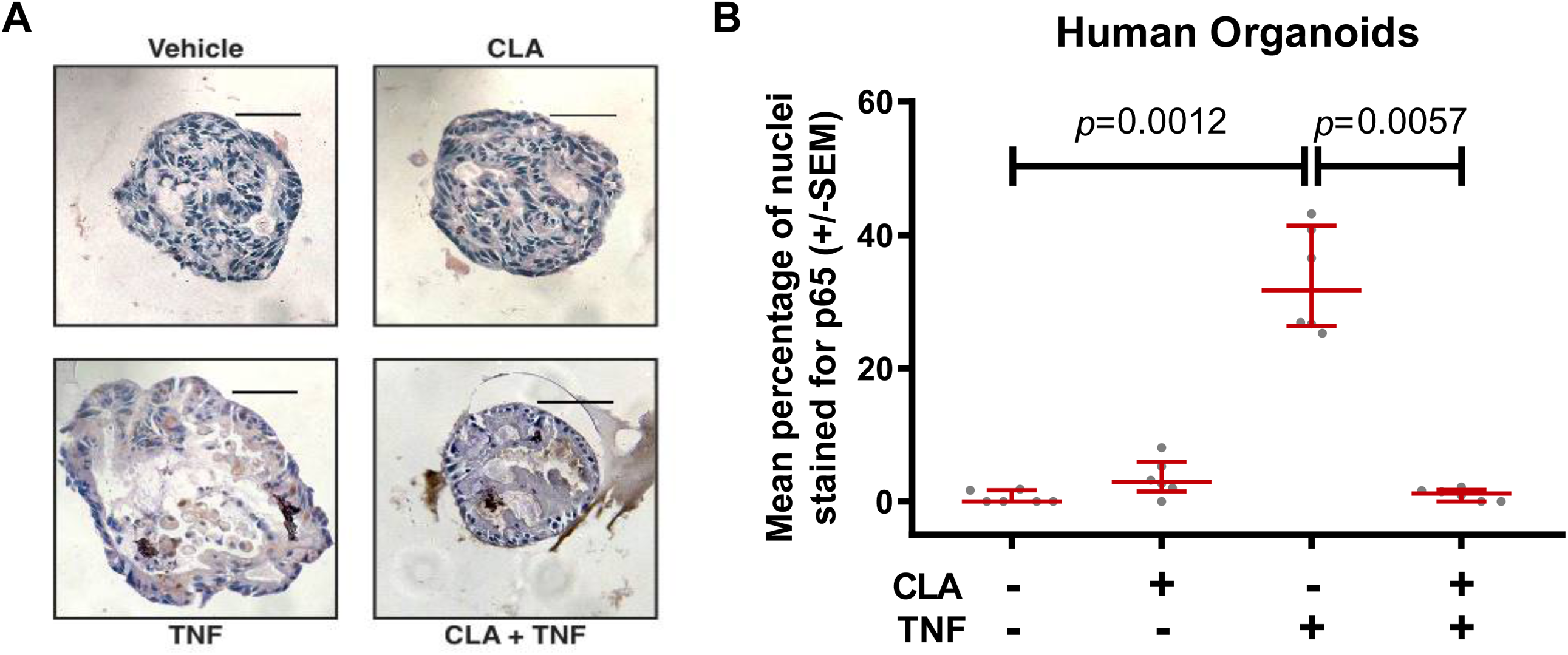
**A:** Representative photomicrographs of human ileal enteroids either unstimulated or stimulated with 100ng/ml TNF, and either co-administered <u>DMSO</u> vehicle or 10µM clarithromycin and immunostained for total p65. **B:** Quantification of nuclear p65 staining of human enteroids. Statistical testing by Kruskal-Wallis 1-way ANOVA and Dunn’s post hoc, N=6.

## Discussion

This article demonstrates that the macrolide antibiotic clarithromycin is a modifier of NF-κB signalling in the gastrointestinal tract. Identification of clarithromycin, using an *in-silico* screen of licensed drugs, demonstrates how novel bioinformatics can be used to progress drug-repurposing, a strategy that has the potential to reduce the cost of future drug development.

The SysmedIBD consortium integrated diverse skill sets to develop this approach which could be applied in different ways in the future. An identical bioinformatic analysis could be used to screen panels of small molecules to identify novel therapeutics for IBD. Similarly, the system could be adapted for use in other contexts where NF-κB signalling is of paramount importance or refocussed onto different signalling networks.

Our list of drugs predicted to influence IBD outcomes included several drugs in routine use for IBD, most prominently the corticosteroids, which are used to treat acute relapses of inflammatory bowel disease^24 25^. The analysis also identified sex hormones including medroxyprogesterone and estradiol which have been shown to modulate colitis^26^, and colitis-associated adenoma development *in vivo*^27^; and non-steroidal anti-inflammatory drugs, which, in clinical practice, are identified as agents that exacerbate IBD^28^. The harmful effects of NSAIDs result from inhibition of constitutively expressed COX-1 in the gastrointestinal epithelium, causing epithelial damage and ulceration; inhibition of COX-2, which is upregulated at sites of inflammation is a well-established anti-inflammatory mechanism, which influences NF-κB signalling. The drug discovery approach deliberately included a bias for agents that alter NF-κB signalling; it is likely that NSAIDs have been selected because of this bias.

These observations demonstrate the importance of interdisciplinary working. The *in-silico* drug discovery model is a powerful tool to identify drugs that may be repurposed, but decisions about which agents to pursue for further analysis can only be made in the context of existing clinical literature.

Laboratory evaluation of clarithromycin aimed to demonstrate proof-of-principle that a drug identified by *in-silico* testing would demonstrate the predicted mechanism of action, and show efficacy *in-vivo*.

Our strategy has potential for development into a higher throughput compound screening system: for example, the hTNF.LucBAC macrophage assay can be performed in a 96-well plate format, and peritoneal macrophages are abundant, simple to extract and highly sensitive to stimulus. Reporter enteroids generated from hTNF.LucBAC mice allowed us to validate the findings seen in peritoneal macrophages in a relevant, untransformed epithelial model using comparable technology, but are unlikely to be amenable to high-throughput assay development due to the challenges (and cost) of maintaining a 3D culture in a small well in a culture plate.

The visualisation of p65.DsRed translocation between the nucleus and cytoplasm in enteric organoids is technically challenging, and not immediately scalable; but it demonstrated fundamental differences in NF-κB signalling dynamics compared to cancer cell lines^7 29^, with TNF-induced p65 oscillations being heavily damped in organoids. This observation demonstrates the value of untransformed primary culture; the mechanisms underlying these differences will be subject to further investigation.

The observation that intestinal NF-κB signalling is altered by clarithromycin *in-vivo* is in keeping with earlier work using cancer cell lines^30^, but it is the first demonstration that macrolides alter this signalling pathway in either untransformed enteroids or gastrointestinal mucosae *in vivo*.

Previous studies demonstrated that a non-antibiotic macrolide, CSY0073, influenced acute DSS colitis in C57BL/6 mice^31^, but clarithromycin had not been studied.

Our final validation was to characterise whether clarithromycin influenced human epithelial NF-κB signalling. We investigated this using a HeLa reporter cell-line model, which is fast, inexpensive, commercially available and could be adapted as a high throughput screening test (figure S2), but it is inferior to the hTNF.LucBAC mouse primary culture models as NF-κB signalling is dysregulated in many cancer cell lines. By generating human enteroids, clarithromycin’s effect on a primary, untransformed human epithelial cell culture could be assessed. Unfortunately, the assay used for NF-κB activation in this model was necessarily less specific, but the results supported those obtained with other assays.

It was serendipitous that the highest-ranked agent identified for repurposing was a drug that had already been trialled in IBD. Importantly the outcomes of previous trials of clarithromycin in IBD have been heterogeneous^20 21 32-34^, suggesting that there may be context-dependent factors that determine whether clarithromycin is useful in a group of patients.

Four published papers, and a conference abstract^33^ have reported the effect of clarithromycin in IBD: they all focussed on Crohn’s disease and were predicated on an antibiotic effect of clarithromycin, either targeting intra-macrophage killing of *E. coli*^*20 21*^ or attempting to eradicate *Mycobacterium avium paratuberculosis*^*32-34*^.

Several factors may explain the discordant outcomes: Selby^32^ and Goodgame^34^ assessed the long-term effect of bacterial eradication; any anti-inflammatory effect would have been lost during the prolonged follow-up period before the primary endpoint was assessed.

In all studies of clarithromycin’s effect on IBD, patient inclusion was based on clinical definitions of active IBD and response assessed by clinical outcome measures. These measures lack objectivity and would not be acceptable endpoints or selection criteria for a current study.

The current study confirms that in addition to its antibiotic effect, clarithromycin has anti-inflammatory properties, which are relevant to gastrointestinal epithelia. Further carefully designed clinical studies will be needed to test the anti-inflammatory effects of treatment with clarithromycin.

Earlier discordant trial results raise questions about trial design and patient selection. In the current era of precision medicine, it is important that patients are carefully selected for treatments, and that agents with previous dichotomous clinical trial results are revisited. To effectively review drugs for precision use, their mechanisms of action need to be understood. This study has helped us to understand how clarithromycin affects NF-κB signalling dynamics, and we hypothesise that it should be possible to select a group of clarithromycin-responsive patients based on their NF-κB signalling status.

Our consortium has recently shown that patients with IBD cluster into several different cohorts based on a dynamic measure of NF-κB responses in peripheral blood monocyte derived macrophages^35^. We hypothesise that, by using this new measure of NF-κB activity, it will be possible to identify IBD patients most likely to respond to NF-κB targeted therapy in the form of clarithromycin, and thereby leverage a precision medicine trial.

In conclusion, our findings strongly suggest that clarithromycin may be a viable, anti-inflammatory therapeutic agent for IBD. In order to progress to a personalised medicine trial of clarithromycin in IBD a partner diagnostic test which can demonstrate altered NF-κB dynamics in patients’ peripheral blood monocyte-derived macrophages is under development^35^, once established this will be used to inform a personalised drug repurposing trial for clarithromycin in IBD.

## Supporting information

Supplementary results

## Conflicts of Interest

VDS is a shareholder and director of GeneXplain GmbH

AK is a shareholder and director of LifeGlimmer GmbH

## Author Contributions

KL performed experiments, acquired original data, analysed data and drafted part of the manuscript. SP performed experiments, acquired original data, analysed data and drafted part of the manuscript. ES performed experiments and acquired original data. PS developed analytical tools, analysed data and drafted part of the manuscript. FB analysed data and drafted part of the manuscript. LM analysed data and drafted part of the manuscript. MK developed analytical pathways and analysed data. HE performed experiments, acquired original data and analysed data. DS supported imaging experiments and reviewed the manuscript. MW devised the project and contributed to funding applications. CD supported organoid work and reviewed the manuscript. BC contributed to funding applications and reviewed the manuscript. VP developed analytical tools and analysed data. VMdS contributed to funding applications and supported the development of analytical pathways. AK contributed to funding applications, supported the development of novel analytical tools, performed data analyses and contributed to the drafting of the manuscript. WM contributed to funding applications and study design, reviewed the manuscript and supported laboratory work. DMP reviewed the manuscript and supported laboratory work. CP contributed to funding applications, designed the study, reviewed the manuscript and supported laboratory work. MB designed the research study, conducted experiments, analysed data, drafted the manuscript and coordinated the project.

## Acknowledgements

This work was supported by the European Union Seventh Framework Programme [FP7/2012–2017] under the SysmedIBD grant, agreement no. 305564

## Notes

#### Summary of Updates

This is a refinement of the original archived version.

