## Supplementary results for "Using systems medicine to identify a therapeutic agent with potential for repurposing in Inflammatory Bowel Disease"

**Supplementary Methods**

**Motif Enrichment Analysis using Logistic Regression** **(MEALR)**

Transcription regulatory feedback loops driven by NF-kappaB/RelA and cooperating transcription factors were inferred based on existing knowledge about signalling pathways involving NF-kappaB/RelA transcription factor binding motifs and ChIP-seq experiment data targeting NF-kappaB/RelA-bound sites in the human genome. The analyses were mainly conducted using the geneXplain platform^1^.

Our analysis aimed to identify feedback loops that encompass components of NF-kappaB/RelA pathways as well as cooperating transcriptional regulators that are themselves regulated by NF-kappaB/RelA (Figure S1).

First, we applied a novel method for motif enrichment analysis using logistic regression (MEALR). MEALR was developed to analyse binding site combinations in sets of DNA sequences that are likely to be bound by a common set of transcriptional regulators, as obtained through ChIP-seq experiments. The algorithm chooses a small set of DNA-sequence motifs from a possibly large library of such models. Given N sequences assigned to classes $y_{i}\in\left\{ 0,1 \right\}$and a library of M positional weight matrices (PWMs), MEALR estimates a sparse logistic regression (LR) model

$$P\left( y|x \right)=\frac{1}{1+e^{-\left( \beta_{0}+\sum\beta_{k}x_{k} \right)}}$$

using the vector of sequence scores calculated for the ith sequence

$$x_{\mathrm{ik}}=log\left( \frac{1}{L_{i}}\sum_{w} \exp\left( S_{w} \right) \right);i=1..N,k=1..M$$

where S_w_ is the log-odds score of the PWM assigned to the wth sequence window. The model coefficients can be used to prioritize motifs of transcriptional regulators with respect to their importance for experimentally observed binding events. Combinations of motifs prioritized by MEALR tend to coincide with transcription factors that are known to cooperate.

We applied MEALR to reveal discriminative DNA-sequence motifs in genomic regions bound by NF-kappaB/RelA after TNF-alpha stimulus. Sequence scores were calculated for a subset from the TRANSFAC(R) database, release 2014.4, consisting of 1429 motifs for transcription factors from vertebrate organisms. Genomic NF-kappaB/RelA binding sites were identified by analysis of ChIP-seq experiments carried out by the ENCODE project [Encode] and deposited in GEO [Gm10847, Gm12878, Gm12891, Gm12892, Gm15510, Gm18505, Gm18526, Gm18951, Gm19099, and Gm19193 (GEO series GSE31477)]. MEALR models were estimated to distinguish ChIP-seq peak sequences from genomic background sequences that were randomly sampled from gene promoter regions not overlapping with peaks of the respective experiment. The analysis was conducted five times for each ChIP-seq data set with different background sequence sets. We retained motifs that were incorporated in the model in all five runs. As MEALR may select similar binding motifs with different identifiers in separate runs, we considered DNA-sequence motif similarity quantified by our method m2match^2^.

A total of 62 TRANSFAC(R) motifs were selected by MEALR in at least 9 of the 10 ChIP-seq data sets. These motifs mapped to 38 transcription factor genes including NF-κB-type factors as well as other transcription factors which our analysis suggests cooperate with NF-κB (Supplementary Table S7).

To identify potential consensus target genes of NF-kappaB/RelA we grouped overlapping peak regions and mapped groups with peaks in at least 6 out of 10 ChIP-seq experiments to nearby genes. This part of the analysis focused on peaks with a maximal length of 3000 bases (>99.1% of all peak regions in the ten ChIP-seq experiments). 4835 consensus ChIP-seq regions were identified in the vicinity of 6329 genes. These genes were defined as consensus target genes.

Among the target genes we identified 24 transcription factors inferred by MEALR as well as 90 of 284 genes encoding components of signalling pathways and cascades involving NF-kappaB/RelA collected from the TRANSPATH(R) database, release 2013.3. These comprise the feedback loops with NF-kappaB/RelA consensus targets described in Figure S1.

### Wnt-3a conditioned media

L Wnt-3a cells (ATCC® CRL-2647™) were seeded into a T75 flask and passaged when ~80% confluent into another T75 flask with 10mL selection media of Dulbecco’s modified eagle’s medium (DMEM) supplemented with 10% foetal calf serum (FCS), 2mM penicillin-streptomycin and 300μg/ml zeocin (Invivogen). When confluent, cells were passaged at a 1:4 split ratio into T150 flasks and 15ml selection media was added. After 24hrs, cells were washed with PBS and the media was changed to 10 mL reduced serum media of DMEM supplemented with 5% FCS and 2 mM penicillin-streptomycin and grown until over-confluent (approximately 4-5 days). The media was collected and centrifuged at 2000 x *g* for 5mins at 4°C. The supernatant was filtered through a 0.22 μm syringe filter (Millipore) and stored at -80°C until use.

### Basal murine organoid media

| Reagent | Final concentration | Supplier |
| --- | --- | --- |
| Advanced DMEM/F12 media | 500 mL | Sigma |
| Glutamax-I | 2 mM | Thermofisher |
| Hepes | 10 mM | Sigma |
| N2 | 1x | Invitrogen |
| B27 | 1x | Invitrogen |
| Antibiotic/antimycotic | 1x | Thermofisher |

### Human seeding media

Seeding media contained 50% human basal media and 50% Wnt-3a conditioned media with the addition of growth factors and inhibitors outlined below.

| Reagent | Final concentration | Supplier |
| --- | --- | --- |
| hEGF | 50 ng/mL | R&D systems |
| hNoggin | 100 ng/mL | R&D systems |
| hR-spondin-1 | 1 μg/mL | R&D systems |
| N-acetylcysteine | 1 mM | Sigma |
| Nicotinamide | 10 mM | Sigma |
| Gastrin | 10 nM | Bachem |
| SB202190 | 10 μM | Sigma |
| PGE2 | 0.01 μM | Sigma |
| LY2157299 | 0.5 μM | Sigma |

### Human organoid culturing media

Seeding media contained 50% human basal media and 50% Wnt-3a conditioned media with the addition of growth factors and inhibitors outlined below.

| Reagent | Final concentration | Supplier |
| --- | --- | --- |
| hEGF | 50 ng/mL | R&D systems |
| hNoggin | 100 ng/mL | R&D systems |
| hR-spondin-1 | 0.5 μg/mL | R&D systems |
| Nicotinamide | 10 mM | Sigma |
| Gastrin | 10 nM | Bachem |
| SB202190 | 10 μM | Sigma |
| PGE2 | 0.01 μM | Sigma |

### Human organoid culture

Five ileal biopsies were taken from non-IBD patients that were enrolled onto the SysMedIBD study and placed into 1 ml PBS containing 1x antibiotic-antimycotic (PBSAA) for transport to the laboratory. In a fume hood, biopsies were washed 10 times in 2 mL PBSAA. Biopsies were places in 4 mL ice cold chelation buffer (3 mM EDTA in PBS) and incubated at room temperature for 30 mins without agitation. Chelation buffer was discarded and replaced with 4 mL shaking buffer (43.3 mM sucrose and 59.4 mM sorbitol in PBS). Biopsies were mechanically agitated in the shaking buffer for 3 mins or until the crypts were no longer seen on the biopsy surface. The crypt suspension was pelleted by centrifugation at 200g for 5mins at 4°C, resuspended in 500 μL Matrigel (Corning, UK) and 50μL was plated out per well of a 24 well plate. Matrigel was polymerised at 37°C before applying 500 μL of human seeding medium per well. After 3 days, human seeding medium was changed to fresh human culturing media which was replaced every 3 days. Organoids were passaged every 7 days.

Terminal ileal organoids were passaged and cultured for 5 days before being removed from the Matrigel using 400 μL cell recovery solution (Corning) on ice for 40 mins. The organoid suspension was transferred to a 15 mL Falcon tube and centrifuged 200 x *g* for 5 mins. The organoids were washed in PBS and resuspended in fresh human culturing media. Pre-treatment with or without 10 μM clarithromycin or 1% DMSO (vehicle control) was applied in suspension for 30 mins. Following pre-treatment, 100 ng/mL recombinant human TNF (Peprotech) was applied for a further 30 mins and then fixed with 4% paraformaldehyde for 20 mins. Fixed organoids were washed with PBS twice and transferred into 300 μL Richard-Allan™ Histogel™ (Thermofisher) and left to polymerise on ice for 30 mins. Once set, the organoid-containing Histogel sample was processed as normal and paraffin embedded.

**Supplementary figures**

**Supplementary figure S1**
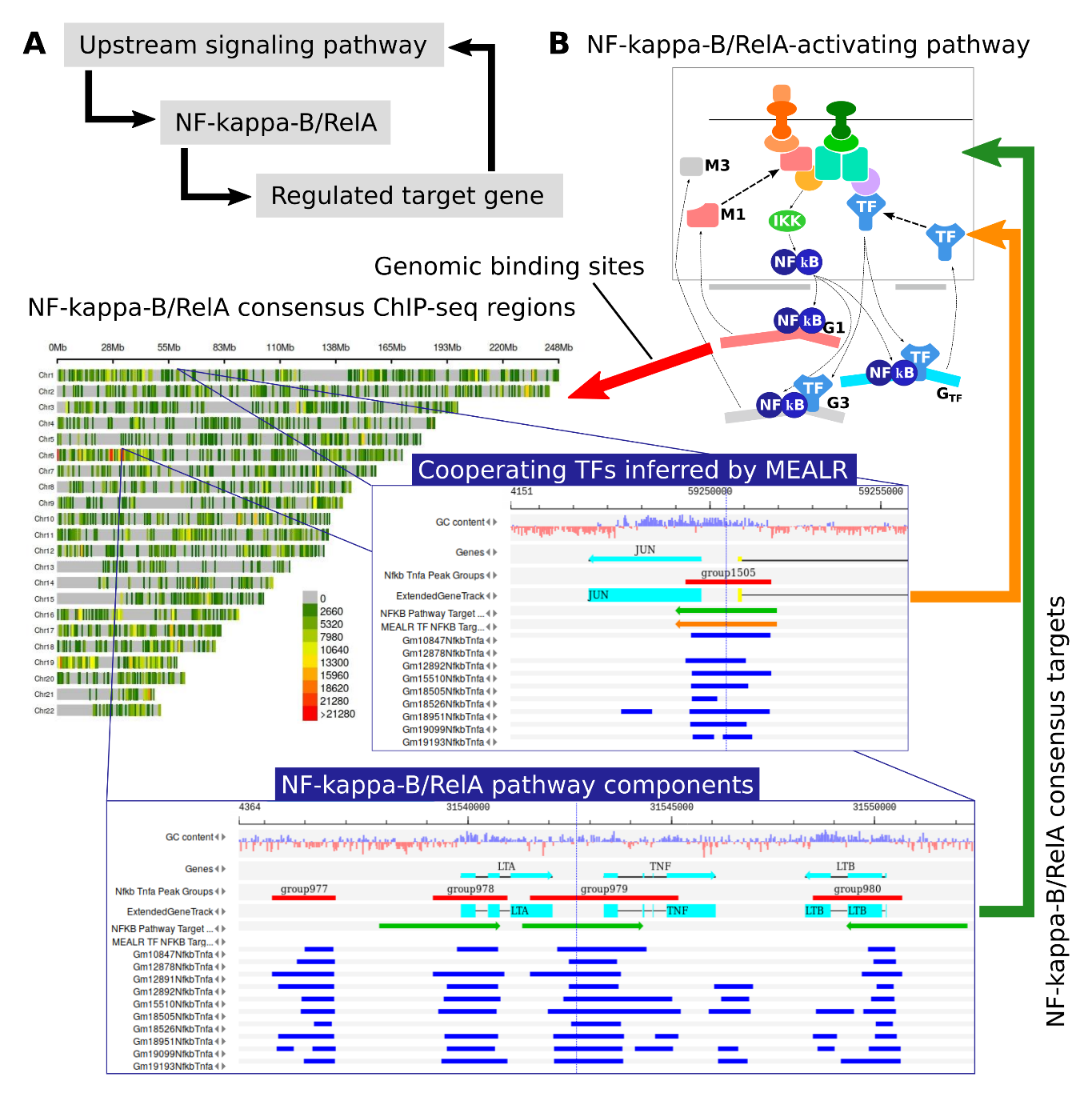
**:** Schematic of the transcriptional feedback loops sought by our analysis. Upon activation through upstream signalling cascades the transcriptional regulator NF-kappa-B/RelA (and cooperating transcription factors) control expression of target genes which may themselves play a role in signalling cascades regulating the activity of NF-kappa-B/RelA thereby establishing a positive or negative feedback loop depending on whether NF-kappa-B/RelA enhances or interferes with the transcription of respective target genes. **B:** More detailed illustration of identified feedback loops. Using consensus NF-kappa-B/RelA binding sites collected from 10 ChIP-seq experiments we analysed their genomic locations with respect to nearby potential target genes which were known components of relevant pathways and/or transcription factors whose motifs played a role for target sequence recognition as inferred by MEALR. Details of the compiled and analysed data are exemplified for pathway target genes TNF, LTA and LTB on chromosome 6 and for the cooperating transcription factor JUN on chromosome 1. The consensus peak density presentation was calculated using the CMplot package. The detailed views of genomic regions were created using the genome browser of the geneXplain platform.

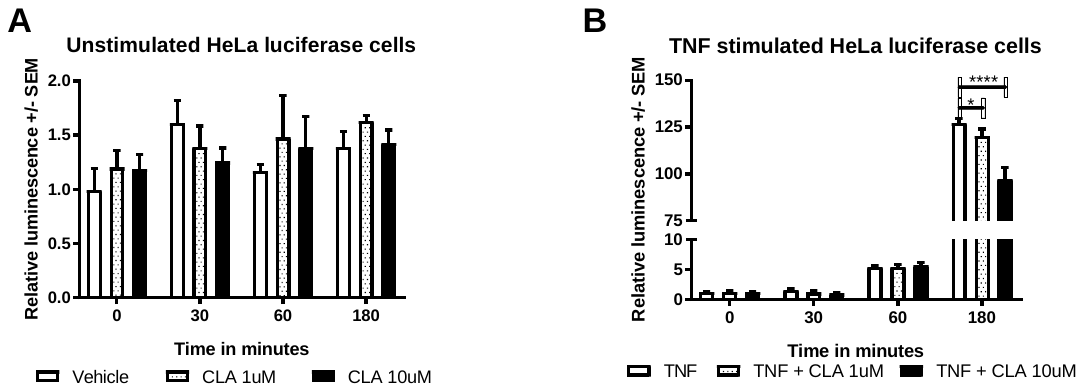

Supplementary Figure S2: Relative luciferase activity in unstimulated HeLa cells treated with DMSO vehicle (open bar), 1uM (hatched bar) or 10uM (solid bar) clarithromycin. B: Relative luciferase activity in TNF stimulated HeLa cells treated with DMSO vehicle, 1uM or 10uM CLA. * denotes p<.05, ****p<.0001, by 2-way ANOVA and Dunnett’s posthoc test.

**Supplementary Tables**

| Gene Symbol 1 | Gene Symbol 2 | Type of interaction | Mouse / Human |
| --- | --- | --- | --- |
| Abl | c-Jun | Process | Human |
| AKT-1 | ASK1 | Process | Human |
| ASK1 | MKK6 | Process | Human |
| Caspase-8 | parkin | Process | Human |
| ERK2 | TAB1 | Process | Human |
| IKK | NF-kappaB1 | Process | Human |
| JNK1 | JNK1alpha1 | Process | Human |
| JNK1 | JNK2 | Process | Human |
| JNK1alpha1 | TAB1 | Process | Human |
| JNK2 | MKK7 | Process | Human |
| MEK1 | MEKK1 | Process | Human |
| MEKK1 | STAT3 | Process | Human |
| MKK6 | TRAF2 | Process | Human |
| MSK1 | p38alpha | Process | Human |
| MyD88 | TLR9 | Process | Human |
| NF-kappaB1 | NF-kappaB1-p50 | Association | Human |
| NF-kappaB1 | p50 | Association | Human |
| NF-kappaB1 | RelA-p65 | Association | Human |
| NF-kappaB1-p50 | p50 | Association | Human |
| NF-kappaB1-p50 | RelA-p65 | Association | Human |
| Nod2 | TRAF2 | Association | Human |
| Nod2 | ubiquitin | Association | Human |
| parkin | ubiquitin | Association | Human |
| TAK1 | TAK1a | Association | Human |
| TAK1a | TRAF2 | Association | Human |
| TAK1a | ubiquitin | Association | Human |
| JIP1 | JNK1 | Process | Mouse |
| PKCzeta | proCaspase-3 | Process | Mouse |
| 26S proteasome | IkappaB-alpha | Process | Both |
| 26S proteasome | p105 | Dissociation | Both |
| 26S proteasome | p50 | Dissociation | Both |
| 26S proteasome | p50 | Process | Both |
| 26S proteasome | RelA-p65 | Process | Both |
| 26S proteasome | ubiquitin | Dissociation | Both |
| 26S proteasome | ubiquitin | Process | Both |
| A20 | IKK | Process | Both |
| A20 | IKK-beta | Process | Both |
| A20 | RIP | Dissociation | Both |
| A20 | TRADD | Dissociation | Both |
| A20 | TRAF2 | Dissociation | Both |
| A20 | ubiquitin | Dissociation | Both |
| Abl | Caspase-8 | Dissociation | Both |
| Abl | IkappaB-alpha | Process | Both |
| acetyl-CoA | carm1 | Process | Both |
| acetyl-CoA | CBP | Process | Both |
| acetyl-CoA | CoA | Process | Both |
| acetyl-CoA | histone H3 | Process | Both |
| acetyl-CoA | p50 | Process | Both |
| acetyl-CoA | RelA-p65 | Process | Both |
| acetyl-CoA | SRC-1 | Process | Both |
| ASK1 | MKK4 | Process | Both |
| ASK1 | TNF-alpha | Process | Both |
| ASK1 | TNFR1 | Process | Both |
| ASK1 | TRADD | Process | Both |
| ASK1 | TRAF2 | Process | Both |
| ASK1 | Trx1 | Association | Both |
| ASK1 | Trx1 | Dissociation | Both |
| Bak | Bid | Association | Both |
| Bak | Bid | Process | Both |
| Bak | Cytochrome C | Process | Both |
| Bak | tBid | Association | Both |
| Bak | tBid | Process | Both |
| Bid | Caspase-8 | Dissociation | Both |
| Bid | Cytochrome C | Process | Both |
| Bid | tBid | Association | Both |
| Bid | tBid | Dissociation | Both |
| Bid | tBid | Process | Both |
| carm1 | CBP | Association | Both |
| carm1 | CBP | Process | Both |
| carm1 | Cdk9 | Association | Both |
| carm1 | Cdk9 | Process | Both |
| carm1 | CoA | Process | Both |
| carm1 | cyclinT1 | Association | Both |
| carm1 | cyclinT1 | Process | Both |
| carm1 | histone H3 | Process | Both |
| carm1 | IL8 | Process | Both |
| carm1 | p50 | Association | Both |
| carm1 | p50 | Process | Both |
| carm1 | RelA-p65 | Association | Both |
| carm1 | RelA-p65 | Process | Both |
| carm1 | S-adenosylhomocysteine | Process | Both |
| carm1 | S-adenosylmethionine | Process | Both |
| carm1 | SRC-1 | Association | Both |
| carm1 | SRC-1 | Process | Both |
| Caspase-2 | CRADD | Association | Both |
| Caspase-2 | CRADD | Dissociation | Both |
| Caspase-2 | proCaspase-2 | Association | Both |
| Caspase-2 | proCaspase-2 | Dissociation | Both |
| Caspase-2 | RIP | Association | Both |
| Caspase-2 | RIP | Dissociation | Both |
| Caspase-2 | TNF-alpha | Association | Both |
| Caspase-2 | TNF-alpha | Dissociation | Both |
| Caspase-2 | TNFR1 | Association | Both |
| Caspase-2 | TNFR1 | Dissociation | Both |
| Caspase-2 | TRADD | Association | Both |
| Caspase-2 | TRADD | Dissociation | Both |
| Caspase-8 | FADD | Association | Both |
| Caspase-8 | FADD | Dissociation | Both |
| Caspase-8 | proCaspase-3 | Process | Both |
| Caspase-8 | tBid | Dissociation | Both |
| Caspase-8 | TNF-alpha | Association | Both |
| Caspase-8 | TNF-alpha | Dissociation | Both |
| Caspase-8 | TNFR1 | Association | Both |
| Caspase-8 | TNFR1 | Dissociation | Both |
| Caspase-8 | TRADD | Association | Both |
| Caspase-8 | TRADD | Dissociation | Both |
| CBP | Cdk9 | Association | Both |
| CBP | Cdk9 | Process | Both |
| CBP | CoA | Process | Both |
| CBP | cyclinT1 | Association | Both |
| CBP | cyclinT1 | Process | Both |
| CBP | histone H3 | Process | Both |
| CBP | IKK | Process | Both |
| CBP | IKK-alpha | Process | Both |
| CBP | IL8 | Process | Both |
| CBP | p50 | Association | Both |
| CBP | p50 | Process | Both |
| CBP | RelA-p65 | Association | Both |
| CBP | RelA-p65 | Process | Both |
| CBP | S-adenosylhomocysteine | Process | Both |
| CBP | S-adenosylmethionine | Process | Both |
| CBP | SRC-1 | Association | Both |
| CBP | SRC-1 | Process | Both |
| CD14 | IRAK-1 | Association | Both |
| CD14 | IRAK-1 | Process | Both |
| CD14 | lbp | Association | Both |
| CD14 | lbp | Process | Both |
| CD14 | LPS | Association | Both |
| CD14 | LPS | Process | Both |
| CD14 | MD-2 | Association | Both |
| CD14 | MD-2 | Process | Both |
| CD14 | MEKK1 | Association | Both |
| CD14 | MyD88 | Association | Both |
| CD14 | MyD88 | Process | Both |
| CD14 | SITPEC | Association | Both |
| CD14 | tab2 | Process | Both |
| CD14 | TLR4 | Association | Both |
| CD14 | TLR4 | Process | Both |
| CD14 | traf6 | Association | Both |
| CD14 | traf6 | Process | Both |
| Cdc34 | Cul-1 | Process | Both |
| Cdc34 | IkappaB-alpha | Process | Both |
| Cdc34 | p50 | Process | Both |
| Cdc34 | RelA-p65 | Process | Both |
| Cdc34 | Ubc5 | Process | Both |
| Cdc34 | ubiquitin | Process | Both |
| Cdk9 | cyclinT1 | Association | Both |
| Cdk9 | cyclinT1 | Process | Both |
| Cdk9 | IL8 | Process | Both |
| Cdk9 | p50 | Association | Both |
| Cdk9 | p50 | Process | Both |
| Cdk9 | RelA-p65 | Association | Both |
| Cdk9 | RelA-p65 | Process | Both |
| Cdk9 | SRC-1 | Association | Both |
| Cdk9 | SRC-1 | Process | Both |
| ceramide | N-SMase | Process | Both |
| ceramide | SM | Process | Both |
| c-Jun | JNK1 | Process | Both |
| c-Jun | JNK2 | Process | Both |
| c-Jun | MKK7 | Process | Both |
| CoA | histone H3 | Process | Both |
| CoA | p50 | Process | Both |
| CoA | RelA-p65 | Process | Both |
| CoA | SRC-1 | Process | Both |
| CRADD | proCaspase-2 | Association | Both |
| CRADD | proCaspase-2 | Dissociation | Both |
| CRADD | RIP | Association | Both |
| CRADD | RIP | Dissociation | Both |
| CRADD | TNF-alpha | Association | Both |
| CRADD | TNF-alpha | Dissociation | Both |
| CRADD | TNFR1 | Association | Both |
| CRADD | TNFR1 | Dissociation | Both |
| CRADD | TRADD | Association | Both |
| CRADD | TRADD | Dissociation | Both |
| Cul-1 | IkappaB-alpha | Association | Both |
| Cul-1 | IkappaB-alpha | Process | Both |
| Cul-1 | p105 | Association | Both |
| Cul-1 | p50 | Association | Both |
| Cul-1 | p50 | Process | Both |
| Cul-1 | RelA-p65 | Association | Both |
| Cul-1 | RelA-p65 | Process | Both |
| Cul-1 | Ubc5 | Process | Both |
| Cul-1 | Ubc5C | Process | Both |
| Cul-1 | ubiquitin | Association | Both |
| Cul-1 | ubiquitin | Process | Both |
| cyclinT1 | IL8 | Process | Both |
| cyclinT1 | p50 | Association | Both |
| cyclinT1 | p50 | Process | Both |
| cyclinT1 | RelA-p65 | Association | Both |
| cyclinT1 | RelA-p65 | Process | Both |
| cyclinT1 | SRC-1 | Association | Both |
| cyclinT1 | SRC-1 | Process | Both |
| Cytochrome C | tBid | Process | Both |
| E1 | tab2 | Process | Both |
| E1 | traf6 | Process | Both |
| E1 | ubiquitin | Process | Both |
| EAC | IKK | Process | Both |
| EAC | IKK-gamma | Process | Both |
| EAC | RIP | Association | Both |
| EAC | RIP | Dissociation | Both |
| EAC | TNF-alpha | Association | Both |
| EAC | TNF-alpha | Dissociation | Both |
| EAC | TNFR1 | Association | Both |
| EAC | TNFR1 | Dissociation | Both |
| EAC | TRADD | Association | Both |
| EAC | TRADD | Dissociation | Both |
| EAC | TRAF2 | Association | Both |
| EAC | TRAF2 | Dissociation | Both |
| EAC | ubiquitin | Dissociation | Both |
| FADD | TNF-alpha | Association | Both |
| FADD | TNF-alpha | Dissociation | Both |
| FADD | TNFR1 | Association | Both |
| FADD | TNFR1 | Dissociation | Both |
| FADD | TRADD | Association | Both |
| FADD | TRADD | Dissociation | Both |
| FAN | N-SMase | Process | Both |
| FAN | RACK1 | Association | Both |
| FAN | RACK1 | Process | Both |
| FAN | SM | Process | Both |
| FAN | TNF-alpha | Association | Both |
| FAN | TNF-alpha | Process | Both |
| FAN | TNFR1 | Association | Both |
| FAN | TNFR1 | Process | Both |
| histone H3 | p50 | Process | Both |
| histone H3 | RelA-p65 | Process | Both |
| histone H3 | S-adenosylhomocysteine | Process | Both |
| histone H3 | S-adenosylmethionine | Process | Both |
| histone H3 | SRC-1 | Process | Both |
| ICAM1 | miR-221/222/222ab/1928 | Process | Both |
| ICAM1 | p50 | Process | Both |
| ICAM1 | RelA-p65 | Process | Both |
| IkappaB-alpha | IKK | Process | Both |
| IkappaB-alpha | IKK-alpha | Process | Both |
| IkappaB-alpha | IKK-beta | Process | Both |
| IkappaB-alpha | IKK-gamma | Process | Both |
| IkappaB-alpha | p50 | Association | Both |
| IkappaB-alpha | p50 | Dissociation | Both |
| IkappaB-alpha | p50 | Process | Both |
| IkappaB-alpha | RelA-p65 | Association | Both |
| IkappaB-alpha | RelA-p65 | Dissociation | Both |
| IkappaB-alpha | RelA-p65 | Process | Both |
| IkappaB-alpha | Ubc5 | Process | Both |
| IkappaB-alpha | Ubc5C | Process | Both |
| IkappaB-alpha | ubiquitin | Process | Both |
| IKK | IKK-alpha | Association | Both |
| IKK | IKK-alpha | Dissociation | Both |
| IKK | IKK-alpha | Process | Both |
| IKK | IKK-beta | Association | Both |
| IKK | IKK-beta | Dissociation | Both |
| IKK | IKK-beta | Process | Both |
| IKK | IKK-gamma | Association | Both |
| IKK | IKK-gamma | Dissociation | Both |
| IKK | IKK-gamma | Process | Both |
| IKK | MEKK1 | Process | Both |
| IKK | NIK | Association | Both |
| IKK | NIK | Dissociation | Both |
| IKK | NIK | Process | Both |
| IKK | p105 | Process | Both |
| IKK | p50 | Process | Both |
| IKK | p62 | Association | Both |
| IKK | p62 | Dissociation | Both |
| IKK | p62 | Process | Both |
| IKK | PKCzeta | Association | Both |
| IKK | PKCzeta | Dissociation | Both |
| IKK | PKCzeta | Process | Both |
| IKK | RelA-p65 | Process | Both |
| IKK | RIP | Association | Both |
| IKK | RIP | Dissociation | Both |
| IKK | RIP | Process | Both |
| IKK | SITPEC | Process | Both |
| IKK | TAB1 | Process | Both |
| IKK | tab2 | Association | Both |
| IKK | tab2 | Dissociation | Both |
| IKK | tab2 | Process | Both |
| IKK | tab3 | Association | Both |
| IKK | tab3 | Dissociation | Both |
| IKK | tab3 | Process | Both |
| IKK | TAK1 | Association | Both |
| IKK | TAK1 | Dissociation | Both |
| IKK | TAK1 | Process | Both |
| IKK | TNF-alpha | Association | Both |
| IKK | TNF-alpha | Dissociation | Both |
| IKK | TNF-alpha | Process | Both |
| IKK | TNFR1 | Association | Both |
| IKK | TNFR1 | Dissociation | Both |
| IKK | TNFR1 | Process | Both |
| IKK | TRADD | Association | Both |
| IKK | TRADD | Dissociation | Both |
| IKK | TRADD | Process | Both |
| IKK | TRAF2 | Association | Both |
| IKK | TRAF2 | Dissociation | Both |
| IKK | TRAF2 | Process | Both |
| IKK | traf6 | Process | Both |
| IKK-alpha | IKK-beta | Association | Both |
| IKK-alpha | IKK-beta | Dissociation | Both |
| IKK-alpha | IKK-beta | Process | Both |
| IKK-alpha | IKK-gamma | Association | Both |
| IKK-alpha | IKK-gamma | Dissociation | Both |
| IKK-alpha | IKK-gamma | Process | Both |
| IKK-alpha | MEKK1 | Process | Both |
| IKK-alpha | NIK | Association | Both |
| IKK-alpha | NIK | Dissociation | Both |
| IKK-alpha | NIK | Process | Both |
| IKK-alpha | p50 | Process | Both |
| IKK-alpha | p62 | Association | Both |
| IKK-alpha | p62 | Dissociation | Both |
| IKK-alpha | p62 | Process | Both |
| IKK-alpha | PKCzeta | Association | Both |
| IKK-alpha | PKCzeta | Dissociation | Both |
| IKK-alpha | PKCzeta | Process | Both |
| IKK-alpha | RelA-p65 | Process | Both |
| IKK-alpha | RIP | Association | Both |
| IKK-alpha | RIP | Dissociation | Both |
| IKK-alpha | RIP | Process | Both |
| IKK-alpha | SITPEC | Process | Both |
| IKK-alpha | TAB1 | Process | Both |
| IKK-alpha | tab2 | Association | Both |
| IKK-alpha | tab2 | Dissociation | Both |
| IKK-alpha | tab2 | Process | Both |
| IKK-alpha | tab3 | Association | Both |
| IKK-alpha | tab3 | Dissociation | Both |
| IKK-alpha | tab3 | Process | Both |
| IKK-alpha | TAK1 | Association | Both |
| IKK-alpha | TAK1 | Dissociation | Both |
| IKK-alpha | TAK1 | Process | Both |
| IKK-alpha | TNF-alpha | Association | Both |
| IKK-alpha | TNF-alpha | Dissociation | Both |
| IKK-alpha | TNF-alpha | Process | Both |
| IKK-alpha | TNFR1 | Association | Both |
| IKK-alpha | TNFR1 | Dissociation | Both |
| IKK-alpha | TNFR1 | Process | Both |
| IKK-alpha | TRADD | Association | Both |
| IKK-alpha | TRADD | Dissociation | Both |
| IKK-alpha | TRADD | Process | Both |
| IKK-alpha | TRAF2 | Association | Both |
| IKK-alpha | TRAF2 | Dissociation | Both |
| IKK-alpha | TRAF2 | Process | Both |
| IKK-alpha | traf6 | Process | Both |
| IKK-beta | IKK-gamma | Association | Both |
| IKK-beta | IKK-gamma | Dissociation | Both |
| IKK-beta | IKK-gamma | Process | Both |
| IKK-beta | MEKK1 | Process | Both |
| IKK-beta | NIK | Association | Both |
| IKK-beta | NIK | Dissociation | Both |
| IKK-beta | NIK | Process | Both |
| IKK-beta | p105 | Process | Both |
| IKK-beta | p50 | Process | Both |
| IKK-beta | p62 | Association | Both |
| IKK-beta | p62 | Dissociation | Both |
| IKK-beta | p62 | Process | Both |
| IKK-beta | PKCzeta | Association | Both |
| IKK-beta | PKCzeta | Dissociation | Both |
| IKK-beta | PKCzeta | Process | Both |
| IKK-beta | RelA-p65 | Process | Both |
| IKK-beta | RIP | Association | Both |
| IKK-beta | RIP | Dissociation | Both |
| IKK-beta | RIP | Process | Both |
| IKK-beta | SITPEC | Process | Both |
| IKK-beta | TAB1 | Process | Both |
| IKK-beta | tab2 | Association | Both |
| IKK-beta | tab2 | Dissociation | Both |
| IKK-beta | tab2 | Process | Both |
| IKK-beta | tab3 | Association | Both |
| IKK-beta | tab3 | Dissociation | Both |
| IKK-beta | tab3 | Process | Both |
| IKK-beta | TAK1 | Association | Both |
| IKK-beta | TAK1 | Dissociation | Both |
| IKK-beta | TAK1 | Process | Both |
| IKK-beta | TNF-alpha | Association | Both |
| IKK-beta | TNF-alpha | Dissociation | Both |
| IKK-beta | TNF-alpha | Process | Both |
| IKK-beta | TNFR1 | Association | Both |
| IKK-beta | TNFR1 | Dissociation | Both |
| IKK-beta | TNFR1 | Process | Both |
| IKK-beta | TRADD | Association | Both |
| IKK-beta | TRADD | Dissociation | Both |
| IKK-beta | TRADD | Process | Both |
| IKK-beta | TRAF2 | Association | Both |
| IKK-beta | TRAF2 | Dissociation | Both |
| IKK-beta | TRAF2 | Process | Both |
| IKK-beta | traf6 | Process | Both |
| IKK-gamma | MEKK1 | Process | Both |
| IKK-gamma | NIK | Association | Both |
| IKK-gamma | NIK | Dissociation | Both |
| IKK-gamma | NIK | Process | Both |
| IKK-gamma | p50 | Process | Both |
| IKK-gamma | p62 | Association | Both |
| IKK-gamma | p62 | Dissociation | Both |
| IKK-gamma | p62 | Process | Both |
| IKK-gamma | PKCzeta | Association | Both |
| IKK-gamma | PKCzeta | Dissociation | Both |
| IKK-gamma | PKCzeta | Process | Both |
| IKK-gamma | RelA-p65 | Process | Both |
| IKK-gamma | RIP | Association | Both |
| IKK-gamma | RIP | Dissociation | Both |
| IKK-gamma | RIP | Process | Both |
| IKK-gamma | SITPEC | Process | Both |
| IKK-gamma | TAB1 | Process | Both |
| IKK-gamma | tab2 | Association | Both |
| IKK-gamma | tab2 | Dissociation | Both |
| IKK-gamma | tab2 | Process | Both |
| IKK-gamma | tab3 | Association | Both |
| IKK-gamma | tab3 | Dissociation | Both |
| IKK-gamma | tab3 | Process | Both |
| IKK-gamma | TAK1 | Association | Both |
| IKK-gamma | TAK1 | Dissociation | Both |
| IKK-gamma | TAK1 | Process | Both |
| IKK-gamma | TNF-alpha | Association | Both |
| IKK-gamma | TNF-alpha | Dissociation | Both |
| IKK-gamma | TNF-alpha | Process | Both |
| IKK-gamma | TNFR1 | Association | Both |
| IKK-gamma | TNFR1 | Dissociation | Both |
| IKK-gamma | TNFR1 | Process | Both |
| IKK-gamma | TRADD | Association | Both |
| IKK-gamma | TRADD | Dissociation | Both |
| IKK-gamma | TRADD | Process | Both |
| IKK-gamma | TRAF2 | Association | Both |
| IKK-gamma | TRAF2 | Dissociation | Both |
| IKK-gamma | TRAF2 | Process | Both |
| IKK-gamma | traf6 | Process | Both |
| IL-10 | IL-1RI | Process | Both |
| IL-10 | IL-1RII | Process | Both |
| IL-1beta-p17 | IL-1RA | Association | Both |
| IL-1beta-p17 | IL-1RA | Process | Both |
| IL-1beta-p17 | IL-1RAcP | Association | Both |
| IL-1beta-p17 | IL-1RAcP | Process | Both |
| IL-1beta-p17 | IL-1RI | Association | Both |
| IL-1beta-p17 | IL-1RI | Process | Both |
| IL-1beta-p17 | IL-1RII | Association | Both |
| IL-1beta-p17 | IRAK-1 | Association | Both |
| IL-1beta-p17 | IRAK-1 | Process | Both |
| IL-1beta-p17 | IRAK-2 | Association | Both |
| IL-1beta-p17 | IRAK-2 | Process | Both |
| IL-1beta-p17 | IRAK-4 | Association | Both |
| IL-1beta-p17 | IRAK-4 | Process | Both |
| IL-1beta-p17 | MEKK1 | Association | Both |
| IL-1beta-p17 | MyD88 | Association | Both |
| IL-1beta-p17 | MyD88 | Process | Both |
| IL-1beta-p17 | Pellino1 | Process | Both |
| IL-1beta-p17 | SITPEC | Association | Both |
| IL-1beta-p17 | tab2 | Process | Both |
| IL-1beta-p17 | traf6 | Association | Both |
| IL-1beta-p17 | traf6 | Process | Both |
| IL-1RA | IL-1RAcP | Association | Both |
| IL-1RA | IL-1RAcP | Process | Both |
| IL-1RA | IL-1RI | Association | Both |
| IL-1RA | IL-1RI | Process | Both |
| IL-1RA | IL-1RII | Association | Both |
| IL-1RA | IRAK-1 | Association | Both |
| IL-1RA | IRAK-1 | Process | Both |
| IL-1RA | IRAK-2 | Association | Both |
| IL-1RA | IRAK-2 | Process | Both |
| IL-1RA | IRAK-4 | Association | Both |
| IL-1RA | IRAK-4 | Process | Both |
| IL-1RA | MEKK1 | Association | Both |
| IL-1RA | MyD88 | Association | Both |
| IL-1RA | MyD88 | Process | Both |
| IL-1RA | Pellino1 | Process | Both |
| IL-1RA | SITPEC | Association | Both |
| IL-1RA | tab2 | Process | Both |
| IL-1RA | traf6 | Association | Both |
| IL-1RA | traf6 | Process | Both |
| IL-1RAcP | IL-1RI | Association | Both |
| IL-1RAcP | IL-1RI | Process | Both |
| IL-1RAcP | IL-1RII | Association | Both |
| IL-1RAcP | IRAK-1 | Association | Both |
| IL-1RAcP | IRAK-1 | Process | Both |
| IL-1RAcP | IRAK-2 | Association | Both |
| IL-1RAcP | IRAK-2 | Process | Both |
| IL-1RAcP | IRAK-4 | Association | Both |
| IL-1RAcP | IRAK-4 | Process | Both |
| IL-1RAcP | MEKK1 | Association | Both |
| IL-1RAcP | MyD88 | Association | Both |
| IL-1RAcP | MyD88 | Process | Both |
| IL-1RAcP | Pellino1 | Process | Both |
| IL-1RAcP | SITPEC | Association | Both |
| IL-1RAcP | tab2 | Process | Both |
| IL-1RAcP | traf6 | Association | Both |
| IL-1RAcP | traf6 | Process | Both |
| IL-1RI | IL-1RII | Association | Both |
| IL-1RI | IL-1RII | Process | Both |
| IL-1RI | IRAK-1 | Association | Both |
| IL-1RI | IRAK-1 | Process | Both |
| IL-1RI | IRAK-2 | Association | Both |
| IL-1RI | IRAK-2 | Process | Both |
| IL-1RI | IRAK-4 | Association | Both |
| IL-1RI | IRAK-4 | Process | Both |
| IL-1RI | MEKK1 | Association | Both |
| IL-1RI | MyD88 | Association | Both |
| IL-1RI | MyD88 | Process | Both |
| IL-1RI | Pellino1 | Process | Both |
| IL-1RI | SITPEC | Association | Both |
| IL-1RI | tab2 | Process | Both |
| IL-1RI | traf6 | Association | Both |
| IL-1RI | traf6 | Process | Both |
| IL8 | p50 | Process | Both |
| IL8 | RelA-p65 | Process | Both |
| IL8 | SRC-1 | Process | Both |
| importin-alpha3 | p50 | Association | Both |
| importin-alpha3 | RelA-p65 | Association | Both |
| IRAK-1 | IRAK-2 | Association | Both |
| IRAK-1 | IRAK-2 | Process | Both |
| IRAK-1 | IRAK-4 | Association | Both |
| IRAK-1 | IRAK-4 | Dissociation | Both |
| IRAK-1 | IRAK-4 | Process | Both |
| IRAK-1 | lbp | Association | Both |
| IRAK-1 | lbp | Process | Both |
| IRAK-1 | LPS | Association | Both |
| IRAK-1 | LPS | Process | Both |
| IRAK-1 | MD-2 | Association | Both |
| IRAK-1 | MD-2 | Process | Both |
| IRAK-1 | MEKK1 | Association | Both |
| IRAK-1 | MyD88 | Association | Both |
| IRAK-1 | MyD88 | Process | Both |
| IRAK-1 | Pellino1 | Dissociation | Both |
| IRAK-1 | Pellino1 | Process | Both |
| IRAK-1 | SITPEC | Association | Both |
| IRAK-1 | tab2 | Dissociation | Both |
| IRAK-1 | tab2 | Process | Both |
| IRAK-1 | TLR4 | Association | Both |
| IRAK-1 | TLR4 | Process | Both |
| IRAK-1 | traf6 | Association | Both |
| IRAK-1 | traf6 | Dissociation | Both |
| IRAK-1 | traf6 | Process | Both |
| IRAK-2 | IRAK-4 | Association | Both |
| IRAK-2 | IRAK-4 | Process | Both |
| IRAK-2 | MEKK1 | Association | Both |
| IRAK-2 | MyD88 | Association | Both |
| IRAK-2 | MyD88 | Process | Both |
| IRAK-2 | Pellino1 | Process | Both |
| IRAK-2 | SITPEC | Association | Both |
| IRAK-2 | tab2 | Process | Both |
| IRAK-2 | traf6 | Association | Both |
| IRAK-2 | traf6 | Process | Both |
| IRAK-4 | MEKK1 | Association | Both |
| IRAK-4 | MyD88 | Association | Both |
| IRAK-4 | MyD88 | Process | Both |
| IRAK-4 | Pellino1 | Dissociation | Both |
| IRAK-4 | Pellino1 | Process | Both |
| IRAK-4 | SITPEC | Association | Both |
| IRAK-4 | tab2 | Dissociation | Both |
| IRAK-4 | tab2 | Process | Both |
| IRAK-4 | traf6 | Association | Both |
| IRAK-4 | traf6 | Dissociation | Both |
| IRAK-4 | traf6 | Process | Both |
| JNK1 | MKK4 | Process | Both |
| JNK1 | MKK7 | Association | Both |
| JNK1 | MKK7 | Process | Both |
| JNK1 | TAB1 | Process | Both |
| JNK2 | MKK4 | Process | Both |
| lbp | LPS | Association | Both |
| lbp | LPS | Process | Both |
| lbp | MD-2 | Association | Both |
| lbp | MD-2 | Process | Both |
| lbp | MEKK1 | Association | Both |
| lbp | MyD88 | Association | Both |
| lbp | MyD88 | Process | Both |
| lbp | SITPEC | Association | Both |
| lbp | tab2 | Process | Both |
| lbp | TLR4 | Association | Both |
| lbp | TLR4 | Process | Both |
| lbp | traf6 | Association | Both |
| lbp | traf6 | Process | Both |
| LPS | MD-2 | Association | Both |
| LPS | MD-2 | Process | Both |
| LPS | MEKK1 | Association | Both |
| LPS | MyD88 | Association | Both |
| LPS | MyD88 | Process | Both |
| LPS | SITPEC | Association | Both |
| LPS | tab2 | Process | Both |
| LPS | TLR4 | Association | Both |
| LPS | TLR4 | Process | Both |
| LPS | traf6 | Association | Both |
| LPS | traf6 | Process | Both |
| MD-2 | MEKK1 | Association | Both |
| MD-2 | MyD88 | Association | Both |
| MD-2 | MyD88 | Process | Both |
| MD-2 | SITPEC | Association | Both |
| MD-2 | tab2 | Process | Both |
| MD-2 | TLR4 | Association | Both |
| MD-2 | TLR4 | Process | Both |
| MD-2 | traf6 | Association | Both |
| MD-2 | traf6 | Process | Both |
| MEKK1 | MKK7 | Process | Both |
| MEKK1 | MyD88 | Association | Both |
| MEKK1 | SITPEC | Association | Both |
| MEKK1 | SITPEC | Process | Both |
| MEKK1 | TLR4 | Association | Both |
| MEKK1 | TNF-alpha | Association | Both |
| MEKK1 | TNF-alpha | Process | Both |
| MEKK1 | TNFR1 | Association | Both |
| MEKK1 | TNFR1 | Process | Both |
| MEKK1 | TRADD | Association | Both |
| MEKK1 | TRADD | Process | Both |
| MEKK1 | TRAF2 | Association | Both |
| MEKK1 | TRAF2 | Process | Both |
| MEKK1 | traf6 | Association | Both |
| MEKK1 | traf6 | Process | Both |
| miR-126-3p | VCAM1 | Process | Both |
| MKK4 | MKK7 | Process | Both |
| MKK4 | TRAF2 | Process | Both |
| MKK6 | p38alpha | Process | Both |
| MKK6 | TAB1 | Process | Both |
| MKK6 | tab2 | Process | Both |
| MKK6 | TAK1 | Process | Both |
| MKK6 | traf6 | Process | Both |
| MKK7 | p38alpha | Process | Both |
| MKK7 | TNF-alpha | Process | Both |
| MKK7 | TNFR1 | Process | Both |
| MKK7 | TRADD | Process | Both |
| MKK7 | TRAF2 | Process | Both |
| MSK1 | p50 | Process | Both |
| MSK1 | RelA-p65 | Process | Both |
| MyD88 | Pellino1 | Process | Both |
| MyD88 | SITPEC | Association | Both |
| MyD88 | ST2 | Association | Both |
| MyD88 | tab2 | Process | Both |
| MyD88 | TLR4 | Association | Both |
| MyD88 | TLR4 | Process | Both |
| MyD88 | TLR5 | Process | Both |
| MyD88 | traf6 | Association | Both |
| MyD88 | traf6 | Process | Both |
| NFKBIA | p50 | Process | Both |
| NFKBIA | RelA-p65 | Process | Both |
| NIK | p62 | Association | Both |
| NIK | p62 | Dissociation | Both |
| NIK | p62 | Process | Both |
| NIK | PKCzeta | Association | Both |
| NIK | PKCzeta | Dissociation | Both |
| NIK | PKCzeta | Process | Both |
| NIK | RIP | Association | Both |
| NIK | RIP | Dissociation | Both |
| NIK | RIP | Process | Both |
| NIK | TNF-alpha | Association | Both |
| NIK | TNF-alpha | Dissociation | Both |
| NIK | TNF-alpha | Process | Both |
| NIK | TNFR1 | Association | Both |
| NIK | TNFR1 | Dissociation | Both |
| NIK | TNFR1 | Process | Both |
| NIK | TRADD | Association | Both |
| NIK | TRADD | Dissociation | Both |
| NIK | TRADD | Process | Both |
| NIK | TRAF2 | Association | Both |
| NIK | TRAF2 | Dissociation | Both |
| NIK | TRAF2 | Process | Both |
| N-SMase | RACK1 | Process | Both |
| N-SMase | SM | Process | Both |
| N-SMase | TNF-alpha | Process | Both |
| N-SMase | TNFR1 | Process | Both |
| p105 | p50 | Dissociation | Both |
| p105 | ubiquitin | Association | Both |
| p105 | ubiquitin | Dissociation | Both |
| p38alpha | TAB1 | Process | Both |
| p50 | Pin1 | Association | Both |
| p50 | PKAc | Process | Both |
| p50 | RelA-p65 | Association | Both |
| p50 | RelA-p65 | Dissociation | Both |
| p50 | RelA-p65 | Process | Both |
| p50 | S-adenosylhomocysteine | Process | Both |
| p50 | S-adenosylmethionine | Process | Both |
| p50 | SELE | Process | Both |
| p50 | SOCS-1 | Association | Both |
| p50 | SOCS-1 | Dissociation | Both |
| p50 | SRC-1 | Association | Both |
| p50 | SRC-1 | Process | Both |
| p50 | TNFAIP3 | Process | Both |
| p50 | Ubc5 | Association | Both |
| p50 | Ubc5 | Process | Both |
| p50 | Ubc5A | Association | Both |
| p50 | Ubc5C | Process | Both |
| p50 | ubiquitin | Association | Both |
| p50 | ubiquitin | Dissociation | Both |
| p50 | ubiquitin | Process | Both |
| p50 | VCAM1 | Process | Both |
| p50 | Wip1 | Process | Both |
| p62 | PKCzeta | Association | Both |
| p62 | PKCzeta | Dissociation | Both |
| p62 | PKCzeta | Process | Both |
| p62 | RIP | Association | Both |
| p62 | RIP | Dissociation | Both |
| p62 | RIP | Process | Both |
| p62 | TNF-alpha | Association | Both |
| p62 | TNF-alpha | Dissociation | Both |
| p62 | TNF-alpha | Process | Both |
| p62 | TNFR1 | Association | Both |
| p62 | TNFR1 | Dissociation | Both |
| p62 | TNFR1 | Process | Both |
| p62 | TRADD | Association | Both |
| p62 | TRADD | Dissociation | Both |
| p62 | TRADD | Process | Both |
| p62 | TRAF2 | Association | Both |
| p62 | TRAF2 | Dissociation | Both |
| p62 | TRAF2 | Process | Both |
| Pellino1 | tab2 | Dissociation | Both |
| Pellino1 | tab2 | Process | Both |
| Pellino1 | traf6 | Dissociation | Both |
| Pellino1 | traf6 | Process | Both |
| Pin1 | RelA-p65 | Association | Both |
| PKAc | RelA-p65 | Process | Both |
| PKCzeta | RIP | Association | Both |
| PKCzeta | RIP | Dissociation | Both |
| PKCzeta | RIP | Process | Both |
| PKCzeta | TNF-alpha | Association | Both |
| PKCzeta | TNF-alpha | Dissociation | Both |
| PKCzeta | TNF-alpha | Process | Both |
| PKCzeta | TNFR1 | Association | Both |
| PKCzeta | TNFR1 | Dissociation | Both |
| PKCzeta | TNFR1 | Process | Both |
| PKCzeta | TRADD | Association | Both |
| PKCzeta | TRADD | Dissociation | Both |
| PKCzeta | TRADD | Process | Both |
| PKCzeta | TRAF2 | Association | Both |
| PKCzeta | TRAF2 | Dissociation | Both |
| PKCzeta | TRAF2 | Process | Both |
| proCaspase-2 | RIP | Association | Both |
| proCaspase-2 | RIP | Dissociation | Both |
| proCaspase-2 | TNF-alpha | Association | Both |
| proCaspase-2 | TNF-alpha | Dissociation | Both |
| proCaspase-2 | TNFR1 | Association | Both |
| proCaspase-2 | TNFR1 | Dissociation | Both |
| proCaspase-2 | TRADD | Association | Both |
| proCaspase-2 | TRADD | Dissociation | Both |
| RACK1 | SM | Process | Both |
| RACK1 | TNF-alpha | Association | Both |
| RACK1 | TNF-alpha | Process | Both |
| RACK1 | TNFR1 | Association | Both |
| RACK1 | TNFR1 | Process | Both |
| RelA-p65 | S-adenosylhomocysteine | Process | Both |
| RelA-p65 | S-adenosylmethionine | Process | Both |
| RelA-p65 | SELE | Process | Both |
| RelA-p65 | SOCS-1 | Association | Both |
| RelA-p65 | SOCS-1 | Dissociation | Both |
| RelA-p65 | SRC-1 | Association | Both |
| RelA-p65 | SRC-1 | Process | Both |
| RelA-p65 | TNFAIP3 | Process | Both |
| RelA-p65 | Ubc5 | Association | Both |
| RelA-p65 | Ubc5 | Process | Both |
| RelA-p65 | Ubc5A | Association | Both |
| RelA-p65 | Ubc5C | Process | Both |
| RelA-p65 | ubiquitin | Association | Both |
| RelA-p65 | ubiquitin | Dissociation | Both |
| RelA-p65 | ubiquitin | Process | Both |
| RelA-p65 | VCAM1 | Process | Both |
| RelA-p65 | Wip1 | Process | Both |
| RIP | tab2 | Association | Both |
| RIP | tab2 | Dissociation | Both |
| RIP | tab2 | Process | Both |
| RIP | tab3 | Association | Both |
| RIP | tab3 | Dissociation | Both |
| RIP | tab3 | Process | Both |
| RIP | TAK1 | Association | Both |
| RIP | TAK1 | Dissociation | Both |
| RIP | TAK1 | Process | Both |
| RIP | TNF-alpha | Association | Both |
| RIP | TNF-alpha | Dissociation | Both |
| RIP | TNF-alpha | Process | Both |
| RIP | TNFR1 | Association | Both |
| RIP | TNFR1 | Dissociation | Both |
| RIP | TNFR1 | Process | Both |
| RIP | TRADD | Association | Both |
| RIP | TRADD | Dissociation | Both |
| RIP | TRADD | Process | Both |
| RIP | TRAF2 | Association | Both |
| RIP | TRAF2 | Dissociation | Both |
| RIP | TRAF2 | Process | Both |
| RIP | ubiquitin | Association | Both |
| RIP | ubiquitin | Dissociation | Both |
| S-adenosylhomocysteine | S-adenosylmethionine | Process | Both |
| S-adenosylhomocysteine | SRC-1 | Process | Both |
| S-adenosylmethionine | SRC-1 | Process | Both |
| SITPEC | TLR4 | Association | Both |
| SITPEC | traf6 | Association | Both |
| SITPEC | traf6 | Process | Both |
| SM | TNF-alpha | Process | Both |
| SM | TNFR1 | Process | Both |
| SOCS-1 | Ubc5 | Association | Both |
| SOCS-1 | Ubc5A | Association | Both |
| SOCS-1 | ubiquitin | Association | Both |
| SOCS-1 | ubiquitin | Dissociation | Both |
| TAB1 | tab2 | Association | Both |
| TAB1 | tab2 | Process | Both |
| TAB1 | TAK1 | Association | Both |
| TAB1 | TAK1 | Process | Both |
| TAB1 | traf6 | Association | Both |
| TAB1 | traf6 | Process | Both |
| tab2 | tab3 | Association | Both |
| tab2 | tab3 | Dissociation | Both |
| tab2 | tab3 | Process | Both |
| tab2 | TAK1 | Association | Both |
| tab2 | TAK1 | Dissociation | Both |
| tab2 | TAK1 | Process | Both |
| tab2 | TLR4 | Process | Both |
| tab2 | TRADD | Association | Both |
| tab2 | TRADD | Dissociation | Both |
| tab2 | TRADD | Process | Both |
| tab2 | TRAF2 | Association | Both |
| tab2 | TRAF2 | Dissociation | Both |
| tab2 | TRAF2 | Process | Both |
| tab2 | traf6 | Association | Both |
| tab2 | traf6 | Dissociation | Both |
| tab2 | traf6 | Process | Both |
| tab2 | ubiquitin | Process | Both |
| tab3 | TAK1 | Association | Both |
| tab3 | TAK1 | Dissociation | Both |
| tab3 | TAK1 | Process | Both |
| tab3 | TRADD | Association | Both |
| tab3 | TRADD | Dissociation | Both |
| tab3 | TRADD | Process | Both |
| tab3 | TRAF2 | Association | Both |
| tab3 | TRAF2 | Dissociation | Both |
| tab3 | TRAF2 | Process | Both |
| TAK1 | TRADD | Association | Both |
| TAK1 | TRADD | Dissociation | Both |
| TAK1 | TRADD | Process | Both |
| TAK1 | TRAF2 | Association | Both |
| TAK1 | TRAF2 | Dissociation | Both |
| TAK1 | TRAF2 | Process | Both |
| TAK1 | traf6 | Association | Both |
| TAK1 | traf6 | Process | Both |
| TAK1 | ubiquitin | Association | Both |
| TLR4 | traf6 | Association | Both |
| TLR4 | traf6 | Process | Both |
| TNF-alpha | TNFR1 | Association | Both |
| TNF-alpha | TNFR1 | Dissociation | Both |
| TNF-alpha | TNFR1 | Process | Both |
| TNF-alpha | TRADD | Association | Both |
| TNF-alpha | TRADD | Dissociation | Both |
| TNF-alpha | TRADD | Process | Both |
| TNF-alpha | TRAF2 | Association | Both |
| TNF-alpha | TRAF2 | Dissociation | Both |
| TNF-alpha | TRAF2 | Process | Both |
| TNF-alpha | ubiquitin | Association | Both |
| TNF-alpha | ubiquitin | Dissociation | Both |
| TNFR1 | TRADD | Association | Both |
| TNFR1 | TRADD | Dissociation | Both |
| TNFR1 | TRADD | Process | Both |
| TNFR1 | TRAF2 | Association | Both |
| TNFR1 | TRAF2 | Dissociation | Both |
| TNFR1 | TRAF2 | Process | Both |
| TNFR1 | ubiquitin | Association | Both |
| TNFR1 | ubiquitin | Dissociation | Both |
| TRADD | TRAF2 | Association | Both |
| TRADD | TRAF2 | Dissociation | Both |
| TRADD | TRAF2 | Process | Both |
| TRADD | ubiquitin | Association | Both |
| TRADD | ubiquitin | Dissociation | Both |
| TRAF2 | ubiquitin | Association | Both |
| TRAF2 | ubiquitin | Dissociation | Both |
| traf6 | ubiquitin | Association | Both |
| traf6 | ubiquitin | Process | Both |
| Ubc5 | Ubc5A | Association | Both |
| Ubc5 | Ubc5C | Process | Both |
| Ubc5 | ubiquitin | Association | Both |
| Ubc5 | ubiquitin | Process | Both |
| Ubc5A | ubiquitin | Association | Both |

Table S1: Interactions between genes and/or proteins identified by text mining algorithm

| ID | Gene symbol | Maximal radius | Reached from set | Reachable total | Score | FDR | Z-Score | Ranks sum |
| --- | --- | --- | --- | --- | --- | --- | --- | --- |
| ENSG00000111537 | *IFNG* | 3.6 | 12 | 71893 | 0.90934 | 0 | 25.515682 | 2 |
| ENSG00000166888 | *STAT6* | 3.6 | 12 | 71893 | 0.90934 | 0 | 25.515682 | 4 |
| ENSG00000106799 | *TGFBR1* | 3.6 | 12 | 71893 | 0.90934 | 0 | 25.24933 | 10 |
| ENSG00000008294 | *SPAG9* | 3.6 | 12 | 71893 | 0.90934 | 0 | 25.109789 | 16 |
| ENSG00000065559 | *MAP2K4* | 3.6 | 12 | 71893 | 0.90934 | 0 | 25.109789 | 16 |
| ENSG00000198909 | *MAP3K3* | 3.6 | 12 | 71893 | 0.90934 | 0 | 25.109789 | 16 |
| ENSG00000148344 | *PTGES* | 3.6 | 12 | 71893 | 0.90934 | 0 | 25.109789 | 20 |
| ENSG00000101665 | *SMAD7* | 3.6 | 12 | 71893 | 0.90934 | 0 | 25.109789 | 22 |
| ENSG00000198742 | *SMURF1* | 3.6 | 12 | 71893 | 0.90934 | 0 | 25.109789 | 22 |
| ENSG00000109471 | *IL2* | 3.6 | 12 | 71893 | 0.90934 | 0 | 25.109789 | 24 |
| ENSG00000120899 | *PTK2B* | 3.6 | 12 | 71893 | 0.90934 | 0 | 23.785404 | 32 |
| ENSG00000138798 | *EGF* | 3.6 | 12 | 71893 | 0.90934 | 0 | 23.785404 | 32 |
| ENSG00000146648 | *EGFR* | 3.6 | 12 | 71893 | 0.90934 | 0 | 23.785404 | 32 |
| ENSG00000197122 | *SRC* | 3.6 | 12 | 71893 | 0.90934 | 0 | 23.785404 | 32 |
| ENSG00000174175 | *SELP* | 3.6 | 12 | 71893 | 0.90934 | 0 | 23.785404 | 34 |
| ENSG00000137462 | *TLR2* | 3.6 | 12 | 71893 | 0.90934 | 0 | 17.603453 | 51 |
| ENSG00000136869 | *TLR4* | 3.6 | 12 | 71893 | 0.90934 | 0 | 17.603453 | 53 |
| ENSG00000076984 | *MAP2K7* | 3.6 | 12 | 71893 | 0.90934 | 0 | 17.603453 | 55 |
| ENSG00000169967 | *MAP3K2* | 3.6 | 12 | 71893 | 0.90934 | 0 | 17.603453 | 55 |
| ENSG00000100030 | *MAPK1* | 3.6 | 12 | 71893 | 0.90934 | 0 | 17.347782 | 58 |
| ENSG00000102882 | *MAPK3* | 3.6 | 12 | 71893 | 0.90934 | 0 | 17.347782 | 60 |
| ENSG00000006210 | *CX3CL1* | 3.6 | 12 | 71893 | 0.90934 | 0 | 17.347782 | 62 |
| ENSG00000130427 | *EPO* | 3.6 | 12 | 71893 | 0.90934 | 0 | 16.325098 | 68 |
| ENSG00000026508 | *CD44* | 3.6 | 12 | 71893 | 0.90934 | 0 | 15.6443615 | 71 |
| ENSG00000136689 | *IL1RN* | 3.6 | 12 | 71893 | 0.90934 | 0 | 15.395652 | 79 |
| ENSG00000197181 | *PIWIL2* | 3.6 | 12 | 71893 | 0.90934 | 0 | 15.6443615 | 81 |
| ENSG00000169429 | *CXCL8* | 3.6 | 12 | 71893 | 0.90934 | 0 | 14.8982315 | 84 |
| ENSG00000107643 | *MAPK8* | 3.6 | 12 | 71893 | 0.90934 | 0 | 14.649521 | 86 |
| ENSG00000115009 | *CCL20* | 3.6 | 12 | 71893 | 0.90934 | 0 | 14.649521 | 88 |
| ENSG00000115353 | *TACR1* | 3.6 | 12 | 71893 | 0.90934 | 0 | 14.649521 | 92 |
| ENSG00000113302 | *IL12B* | 3.6 | 12 | 71893 | 0.90934 | 0 | 14.649521 | 94 |
| ENSG00000128052 | *KDR* | 3.6 | 12 | 71893 | 0.90934 | 0 | 12.908551 | 102 |
| ENSG00000112715 | *VEGFA* | 3.6 | 12 | 71893 | 0.90934 | 0 | 12.908551 | 104 |
| ENSG00000142208 | *AKT1* | 3.6 | 12 | 71893 | 0.90934 | 0 | 12.411131 | 108 |
| ENSG00000163464 | *CXCR1* | 3.6 | 12 | 71893 | 0.90934 | 0 | 9.924029 | 114 |
| ENSG00000136244 | *IL6* | 3.6 | 12 | 71893 | 0.90934 | 0 | 9.177899 | 123 |
| ENSG00000087245 | *MMP2* | 3.6 | 12 | 71893 | 0.90934 | 0 | 8.92919 | 126 |
| ENSG00000125538 | *IL1B* | 3.6 | 12 | 71893 | 0.90934 | 0 | 9.177899 | 126 |
| ENSG00000100985 | *MMP9* | 3.6 | 12 | 71893 | 0.90934 | 0 | 8.680479 | 132 |
| ENSG00000137752 | *CASP1* | 3.6 | 12 | 71893 | 0.90934 | 0 | 8.92919 | 134 |
| ENSG00000149269 | *PAK1* | 3.6 | 12 | 71893 | 0.90934 | 0 | 8.444952 | 137 |
| ENSG00000002549 | *LAP3* | 3.6 | 12 | 71893 | 0.90934 | 0 | 8.444952 | 139 |
| ENSG00000092969 | *TGFB2* | 3.6 | 12 | 71893 | 0.90934 | 0 | 8.444952 | 139 |
| ENSG00000119699 | *TGFB3* | 3.6 | 12 | 71893 | 0.90934 | 0 | 8.444952 | 139 |
| ENSG00000143768 | *LEFTY2* | 3.6 | 12 | 71893 | 0.90934 | 0 | 8.444952 | 139 |
| ENSG00000049323 | *LTBP1* | 3.6 | 12 | 71893 | 0.90934 | 0 | 8.444952 | 141 |
| ENSG00000090006 | *LTBP4* | 3.6 | 12 | 71893 | 0.90934 | 0 | 8.444952 | 141 |
| ENSG00000119681 | *LTBP2* | 3.6 | 12 | 71893 | 0.90934 | 0 | 8.444952 | 141 |
| ENSG00000168056 | *LTBP3* | 3.6 | 12 | 71893 | 0.90934 | 0 | 8.444952 | 141 |
| ENSG00000134759 | *ELP2* | 3.6 | 12 | 71893 | 0.90934 | 0 | 7.4784894 | 157 |
| ENSG00000115415 | *STAT1* | 3.6 | 12 | 71893 | 0.90934 | 0 | 7.4784894 | 161 |
| ENSG00000105397 | *TYK2* | 3.6 | 12 | 71893 | 0.90934 | 0 | 7.4784894 | 165 |
| ENSG00000110324 | *IL10RA* | 3.6 | 12 | 71893 | 0.90934 | 0 | 7.4784894 | 165 |
| ENSG00000136634 | *IL10* | 3.6 | 12 | 71893 | 0.90934 | 0 | 7.4784894 | 165 |
| ENSG00000162434 | *JAK1* | 3.6 | 12 | 71893 | 0.90934 | 0 | 7.4784894 | 165 |
| ENSG00000243646 | *IL10RB* | 3.6 | 12 | 71893 | 0.90934 | 0 | 7.4784894 | 165 |
| ENSG00000164400 | *CSF2* | 3.6 | 12 | 71893 | 0.90934 | 0 | 6.7536426 | 174 |
| ENSG00000115008 | *IL1A* | 3.6 | 12 | 71893 | 0.90934 | 0 | 6.7536426 | 180 |
| ENSG00000110944 | *IL23A* | 3.6 | 12 | 71893 | 0.90934 | 0 | 6.7536426 | 182 |
| ENSG00000138378 | *STAT4* | 3.6 | 12 | 71893 | 0.90934 | 0 | 5.862493 | 190 |
| ENSG00000113580 | *NR3C1* | 3.6 | 12 | 71893 | 0.90934 | 0 | 5.862493 | 192 |
| ENSG00000134352 | *IL6ST* | 3.6 | 12 | 71893 | 0.90934 | 0 | 5.393531 | 196 |
| ENSG00000160712 | *IL6R* | 3.6 | 12 | 71893 | 0.90934 | 0 | 5.393531 | 198 |
| ENSG00000069702 | *TGFBR3* | 3.6 | 12 | 71893 | 0.90934 | 0 | 5.393531 | 200 |
| ENSG00000163513 | *TGFBR2* | 3.6 | 12 | 71893 | 0.90934 | 0 | 5.393531 | 200 |
| ENSG00000017427 | *IGF1* | 3.6 | 12 | 71893 | 0.90934 | 0 | 5.393531 | 202 |
| ENSG00000168036 | *CTNNB1* | 3.6 | 12 | 71893 | 0.90934 | 0 | 5.393531 | 204 |
| ENSG00000164761 | *TNFRSF11B* | 3.6 | 12 | 71893 | 0.90934 | 0 | 5.1354814 | 208 |
| ENSG00000164136 | *IL15* | 3.6 | 12 | 71893 | 0.90934 | 0 | 5.1354814 | 210 |
| ENSG00000118689 | *FOXO3* | 3.6 | 12 | 71893 | 0.90934 | 0 | 3.9684856 | 214 |
| ENSG00000175387 | *SMAD2* | 3.6 | 12 | 71893 | 0.90934 | 0 | 3.7350864 | 216 |
| ENSG00000108691 | *CCL2* | 3.6 | 12 | 71893 | 0.90934 | 0 | 3.7350864 | 218 |
| ENSG00000157404 | *KIT* | 3.6 | 12 | 71893 | 0.90934 | 0 | 3.7350864 | 220 |
| ENSG00000160791 | *CCR5* | 3.6 | 12 | 71893 | 0.90934 | 0 | 3.7350864 | 224 |
| ENSG00000039068 | *CDH1* | 3.6 | 12 | 71893 | 0.90934 | 0 | 3.7350864 | 226 |
| ENSG00000112116 | *IL17F* | 3.6 | 12 | 71893 | 0.90934 | 0 | 3.7350864 | 230 |
| ENSG00000112115 | *IL17A* | 3.6 | 12 | 71893 | 0.90934 | 0 | 3.4077682 | 233 |
| ENSG00000090339 | *ICAM1* | 3.6 | 12 | 71893 | 0.90934 | 0 | 3.1813836 | 239 |
| ENSG00000102245 | *CD40LG* | 3.6 | 12 | 71893 | 0.90934 | 0 | 3.1813836 | 243 |
| ENSG00000173327 | *MAP3K11* | 3.6 | 12 | 71893 | 0.90934 | 0 | 3.1813836 | 245 |
| ENSG00000144381 | *HSPD1* | 3.6 | 12 | 71893 | 0.90934 | 0 | 2.5022295 | 269 |
| ENSG00000168610 | *STAT3* | 3.6 | 12 | 71893 | 0.90934 | 0 | 2.5022295 | 271 |
| ENSG00000149968 | *MMP3* | 3.6 | 12 | 71893 | 0.90934 | 0 | 2.4914422 | 275 |
| ENSG00000006062 | *MAP3K14* | 5.46 | 12 | 70106 | 0.74578536 | 0 | 3.4883113 | 277 |
| ENSG00000106683 | *LIMK1* | 3.6 | 12 | 71893 | 0.90934 | 0 | 2.4914422 | 279 |
| ENSG00000111321 | *LTBR* | 5.46 | 12 | 70106 | 0.745759 | 0 | 3.3829823 | 281 |
| ENSG00000226979 | *LTA* | 5.46 | 12 | 70106 | 0.745759 | 0 | 3.3829823 | 281 |
| ENSG00000227507 | *LTB* | 5.46 | 12 | 70106 | 0.745759 | 0 | 3.3829823 | 281 |
| ENSG00000115541 | *HSPE1* | 3.6 | 12 | 71893 | 0.90934 | 0 | 2.4806724 | 283 |
| ENSG00000164305 | *CASP3* | 3.6 | 12 | 71893 | 0.90934 | 0 | 2.4806724 | 283 |
| ENSG00000113525 | *IL5* | 3.6 | 12 | 71893 | 0.90934 | 0 | 2.2562785 | 288 |
| ENSG00000064012 | *CASP8* | 3.6 | 12 | 71893 | 0.90934 | 0 | 2.2562785 | 292 |
| ENSG00000072062 | *PRKACA* | 5.6 | 12 | 70973 | 0.807201 | 0.003 | 2.7416925 | 292 |
| ENSG00000232810 | *TNF* | 3.6 | 12 | 71893 | 0.90934 | 0 | 2.2562785 | 296 |
| ENSG00000067182 | *TNFRSF1A* | 3.6 | 12 | 71893 | 0.90934 | 0 | 2.031885 | 322 |
| ENSG00000108094 | *CUL2* | 6.12 | 12 | 70584 | 0.7465518 | 0.012 | 2.1583393 | 326 |
| ENSG00000115457 | *IGFBP2* | 6.75 | 12 | 70055 | 0.61936754 | 0.006 | 2.8468504 | 338 |
| ENSG00000173039 | *RELA* | 6.75 | 12 | 70046 | 0.6259042 | 0.017 | 2.6685882 | 339 |
| ENSG00000105810 | *CDK6* | 6.75 | 12 | 70044 | 0.6250095 | 0.028 | 2.542868 | 345 |
| ENSG00000147162 | *OGT* | 6.75 | 12 | 70045 | 0.6155797 | 0.014 | 2.6078267 | 348 |
| ENSG00000164088 | *PPM1M* | 6.75 | 12 | 70029 | 0.6124364 | 0.007 | 2.5822856 | 351 |
| ENSG00000005381 | *MPO* | 5.3999996 | 12 | 71017 | 0.82925606 | 0.005 | 1.8166571 | 352 |
| ENSG00000107968 | *MAP3K8* | 6.6 | 12 | 70129 | 0.71335024 | 0.001 | 1.8473707 | 358 |
| ENSG00000110330 | *BIRC2* | 5.5 | 12 | 70517 | 0.7119078 | 0.017 | 1.8184592 | 361 |
| ENSG00000067900 | *ROCK1* | 6.75 | 12 | 70037 | 0.6123565 | 0.036 | 2.451285 | 364 |
| ENSG00000104365 | *IKBKB* | 6.75 | 12 | 70119 | 0.6526765 | 0.015 | 2.1128314 | 366 |
| ENSG00000134058 | *CDK7* | 6.75 | 12 | 70030 | 0.6123358 | 0.043 | 2.346676 | 366 |
| ENSG00000269335 | *IKBKG* | 5.75 | 12 | 70869 | 0.71404743 | 0.03 | 1.6740329 | 384 |
| ENSG00000131788 | *PIAS3* | 5.76 | 12 | 70778 | 0.71407497 | 0.04 | 1.6588471 | 391 |
| ENSG00000163702 | *IL17RC* | 6.75 | 12 | 70099 | 0.612196 | 0.047 | 2.0695095 | 393 |
| ENSG00000177663 | *IL17RA* | 6.75 | 12 | 70099 | 0.612196 | 0.047 | 2.0695095 | 393 |
| ENSG00000213341 | *CHUK* | 6.75 | 12 | 70125 | 0.6153272 | 0.041 | 2.021259 | 394 |
| ENSG00000055208 | *TAB2* | 6.75 | 12 | 70099 | 0.612196 | 0.047 | 2.0686812 | 394 |
| ENSG00000056972 | *TRAF3IP2* | 6.75 | 12 | 70099 | 0.612196 | 0.047 | 2.069004 | 395 |
| ENSG00000131323 | *TRAF3* | 7.035 | 12 | 65994 | 0.5407338 | 0.006 | 2.5663161 | 396 |
| ENSG00000175104 | *TRAF6* | 6.75 | 12 | 70099 | 0.612196 | 0.047 | 2.0683603 | 398 |
| ENSG00000162924 | *REL* | 9.9 | 12 | 59326 | 0.46628165 | 0.001 | 6.650155 | 402 |
| ENSG00000135341 | *MAP3K7* | 5.76 | 12 | 70043 | 0.6245046 | 0.036 | 1.7717301 | 406 |
| ENSG00000129559 | *NEDD8* | 8.28 | 12 | 66741 | 0.54859954 | 0.027 | 2.1377385 | 423 |
| ENSG00000173801 | *JUP* | 7.56 | 12 | 68114 | 0.59836376 | 0.003 | 1.7430931 | 428 |
| ENSG00000105329 | *TGFB1* | 3.6 | 12 | 71893 | 0.90934 | 0 | 1.3587039 | 429 |
| ENSG00000149591 | *TAGLN* | 7.6 | 12 | 68077 | 0.59836614 | 0.005 | 1.7195715 | 432 |
| ENSG00000167749 | *KLK4* | 7.6 | 12 | 68070 | 0.5983616 | 0.004 | 1.731232 | 433 |
| ENSG00000115170 | *ACVR1* | 7.5950003 | 12 | 68069 | 0.5983511 | 0.006 | 1.6844014 | 439 |
| ENSG00000152484 | *USP12* | 7.6 | 12 | 68084 | 0.5983578 | 0.006 | 1.6811485 | 442 |
| ENSG00000131653 | *TRAF7* | 6.8 | 12 | 69963 | 0.63348943 | 0.05 | 1.5657457 | 445 |
| ENSG00000121989 | *ACVR2A* | 7.5600004 | 12 | 68115 | 0.5983318 | 0.007 | 1.693381 | 445 |
| ENSG00000122641 | *INHBA* | 7.5600004 | 12 | 68115 | 0.5983318 | 0.007 | 1.693381 | 445 |
| ENSG00000135503 | *ACVR1B* | 7.5600004 | 12 | 68115 | 0.5983318 | 0.007 | 1.693381 | 445 |
| ENSG00000172349 | *IL16* | 7.6 | 12 | 68079 | 0.5983425 | 0.007 | 1.681531 | 446 |
| ENSG00000160633 | *SAFB* | 7.6 | 12 | 68067 | 0.598348 | 0.007 | 1.6682975 | 449 |
| ENSG00000141552 | *ANAPC11* | 7.6 | 12 | 68076 | 0.598261 | 0.006 | 1.6981487 | 449 |
| ENSG00000172840 | *PDP2* | 7.5950003 | 12 | 68080 | 0.5983457 | 0.007 | 1.6640131 | 454 |
| ENSG00000139567 | *ACVRL1* | 7.5950003 | 12 | 68070 | 0.5983349 | 0.007 | 1.6671054 | 455 |
| ENSG00000136807 | *CDK9* | 7.5950003 | 12 | 68107 | 0.60283357 | 0.012 | 1.6284322 | 457 |
| ENSG00000100234 | *TIMP3* | 7.5599995 | 12 | 68137 | 0.5983164 | 0.006 | 1.6641697 | 459 |
| ENSG00000164850 | *GPER1* | 3.6 | 12 | 71894 | 0.909338 | 0 | 1.2649971 | 460 |
| ENSG00000112799 | *LY86* | 3.6 | 12 | 71894 | 0.909338 | 0 | 1.2615731 | 462 |
| ENSG00000134061 | *CD180* | 3.6 | 12 | 71894 | 0.909338 | 0 | 1.2615731 | 462 |
| ENSG00000169118 | *CSNK1G1* | 6.8499994 | 12 | 69866 | 0.63015866 | 0.02 | 1.424171 | 465 |
| ENSG00000115718 | *PROC* | 5.91 | 12 | 70681 | 0.71089166 | 0.037 | 1.2780058 | 471 |
| ENSG00000178568 | *ERBB4* | 7.5950003 | 12 | 68082 | 0.5982364 | 0.007 | 1.6562477 | 471 |
| ENSG00000065361 | *ERBB3* | 7.5950003 | 12 | 68082 | 0.59823585 | 0.007 | 1.6563139 | 471 |
| ENSG00000141736 | *ERBB2* | 7.5950003 | 12 | 68082 | 0.5982351 | 0.007 | 1.6563655 | 471 |
| ENSG00000104856 | *RELB* | 4.32 | 2 | 78 | 0.25144747 | 0.002 | 9.694386 | 474 |
| ENSG00000135960 | *EDAR* | 5.76 | 12 | 69263 | 0.62076813 | 0.001 | 1.4052681 | 476 |
| ENSG00000089041 | *P2RX7* | 6.6 | 12 | 70129 | 0.7012792 | 0.001 | 1.3085307 | 477 |
| ENSG00000100079 | *LGALS2* | 6.6 | 12 | 70130 | 0.70127815 | 0.001 | 1.3114457 | 478 |
| ENSG00000164951 | *PDP1* | 7.5950003 | 12 | 68069 | 0.5983471 | 0.011 | 1.5548629 | 479 |
| ENSG00000180772 | *AGTR2* | 5.91 | 12 | 70682 | 0.7108893 | 0.044 | 1.2524608 | 481 |
| ENSG00000095739 | *BAMBI* | 6.6 | 12 | 70129 | 0.70127857 | 0.002 | 1.3057415 | 481 |
| ENSG00000107014 | *RLN2* | 6.6 | 12 | 70130 | 0.7012774 | 0.001 | 1.308338 | 482 |
| ENSG00000107779 | *BMPR1A* | 7.5950003 | 12 | 68083 | 0.5980402 | 0.012 | 1.5984213 | 482 |
| ENSG00000150093 | *ITGB1* | 9.585 | 12 | 60076 | 0.41584295 | 0.005 | 2.849528 | 482 |
| ENSG00000105647 | *PIK3R2* | 7.9999995 | 12 | 66901 | 0.5182761 | 0.031 | 1.6607733 | 483 |
| ENSG00000128342 | *LIF* | 6.6 | 12 | 70129 | 0.70127815 | 0.001 | 1.3043623 | 484 |
| ENSG00000176797 | *DEFB103A* | 6.6 | 12 | 70130 | 0.7012766 | 0.001 | 1.3081578 | 484 |
| ENSG00000177243 | *DEFB103B* | 6.6 | 12 | 70130 | 0.7012766 | 0.001 | 1.3081578 | 484 |
| ENSG00000115687 | *PASK* | 6.75 | 12 | 70172 | 0.6347166 | 0.032 | 1.3217078 | 484 |
| ENSG00000112425 | *EPM2A* | 6.75 | 12 | 70180 | 0.6347163 | 0.032 | 1.3190727 | 486 |
| ENSG00000043093 | *DCUN1D1* | 9.27 | 12 | 63476 | 0.45494947 | 0.014 | 2.119744 | 486 |
| ENSG00000171855 | *IFNB1* | 6.6 | 12 | 70138 | 0.7012685 | 0.002 | 1.3028532 | 488 |
| ENSG00000145675 | *PIK3R1* | 7.9999995 | 12 | 66901 | 0.5182743 | 0.032 | 1.6557599 | 491 |
| ENSG00000104814 | *MAP4K1* | 7.5600004 | 12 | 65212 | 0.51734453 | 0.044 | 2.010294 | 494 |
| ENSG00000136279 | *DBNL* | 7.5600004 | 12 | 65212 | 0.51734453 | 0.044 | 2.010294 | 494 |
| ENSG00000010671 | *BTK* | 9.599999 | 12 | 60063 | 0.4158373 | 0.007 | 2.618889 | 495 |
| ENSG00000101977 | *MCF2* | 7.6 | 12 | 68110 | 0.59817326 | 0.018 | 1.5310167 | 496 |
| ENSG00000108389 | *MTMR4* | 7.5950003 | 12 | 68134 | 0.59828806 | 0.015 | 1.4546616 | 499 |
| ENSG00000157450 | *RNF111* | 9.38 | 12 | 62732 | 0.45513147 | 0.017 | 1.8615283 | 504 |
| ENSG00000204209 | *DAXX* | 7.5599995 | 12 | 68194 | 0.5978195 | 0.02 | 1.4358637 | 510 |
| ENSG00000100393 | *EP300* | 8 | 12 | 67011 | 0.5181211 | 0.045 | 1.5372753 | 511 |
| ENSG00000110958 | *PTGES3* | 9.599999 | 12 | 60057 | 0.41586527 | 0.009 | 2.2056735 | 516 |
| ENSG00000126351 | *THRA* | 9.599999 | 12 | 60093 | 0.41587228 | 0.006 | 2.1791701 | 517 |
| ENSG00000174775 | *HRAS* | 9.57 | 12 | 60167 | 0.41561633 | 0.015 | 2.4829588 | 517 |
| ENSG00000100983 | *GSS* | 9.599999 | 12 | 60601 | 0.41573972 | 0.012 | 2.2672687 | 518 |
| ENSG00000180370 | *PAK2* | 7.5950003 | 12 | 68408 | 0.59760123 | 0.023 | 1.4013838 | 522 |
| ENSG00000118503 | *TNFAIP3* | 3.6 | 12 | 71893 | 0.90934 | 0 | 1.1343101 | 528 |
| ENSG00000197081 | *IGF2R* | 9 | 12 | 63755 | 0.42712894 | 0.01 | 1.8112867 | 528 |
| ENSG00000131196 | *NFATC1* | 7.95 | 12 | 67021 | 0.517943 | 0.022 | 1.4142506 | 529 |
| ENSG00000169194 | *IL13* | 3.6 | 12 | 71893 | 0.90934 | 0 | 1.1343101 | 530 |
| ENSG00000113520 | *IL4* | 3.6 | 12 | 71893 | 0.90934 | 0 | 1.1343101 | 532 |
| ENSG00000068903 | *SIRT2* | 7.8 | 12 | 67853 | 0.554528 | 0.02 | 1.3698407 | 532 |
| ENSG00000147099 | *HDAC8* | 7.95 | 12 | 67030 | 0.5179252 | 0.023 | 1.3974826 | 534 |
| ENSG00000150782 | *IL18* | 3.6 | 12 | 71893 | 0.90934 | 0 | 1.1343101 | 535 |
| ENSG00000159110 | *IFNAR2* | 9 | 12 | 63761 | 0.42710644 | 0.013 | 1.7704278 | 535 |
| ENSG00000007171 | *NOS2* | 3.6 | 12 | 71893 | 0.90934 | 0 | 1.1343101 | 537 |
| ENSG00000048052 | *HDAC9* | 7.95 | 12 | 67033 | 0.51792264 | 0.023 | 1.3973957 | 537 |
| ENSG00000115594 | *IL1R1* | 3.6 | 12 | 71893 | 0.90934 | 0 | 1.1343101 | 539 |
| ENSG00000185338 | *SOCS1* | 6.91 | 12 | 69124 | 0.6298157 | 0.026 | 1.2146442 | 539 |
| ENSG00000082146 | *STRADB* | 9.595 | 12 | 60290 | 0.41571394 | 0.014 | 2.0868638 | 539 |
| ENSG00000061273 | *HDAC7* | 7.95 | 12 | 67065 | 0.51789963 | 0.026 | 1.3862103 | 541 |
| ENSG00000094631 | *HDAC6* | 7.95 | 12 | 67032 | 0.51791775 | 0.025 | 1.3618859 | 546 |
| ENSG00000108840 | *HDAC5* | 7.95 | 12 | 67030 | 0.5179232 | 0.026 | 1.3528996 | 547 |
| ENSG00000156711 | *MAPK13* | 7.5950003 | 12 | 68171 | 0.59766006 | 0.034 | 1.285942 | 549 |
| ENSG00000117399 | *CDC20* | 9.599999 | 12 | 60352 | 0.41544947 | 0.013 | 1.9345831 | 556 |
| ENSG00000188130 | *MAPK12* | 7.5950003 | 12 | 68166 | 0.59763104 | 0.037 | 1.2603947 | 559 |
| ENSG00000108443 | *RPS6KB1* | 9.585 | 12 | 61285 | 0.41531238 | 0.019 | 1.9173793 | 559 |
| ENSG00000117984 | *CTSD* | 7.95 | 12 | 67071 | 0.51790935 | 0.029 | 1.2937016 | 565 |
| ENSG00000213281 | *NRAS* | 9.57 | 12 | 60140 | 0.4156918 | 0.013 | 1.7966001 | 566 |
| ENSG00000185104 | *FAF1* | 5.76 | 12 | 70777 | 0.7104392 | 0.026 | 1.0897201 | 570 |
| ENSG00000129422 | *MTUS1* | 9 | 12 | 63755 | 0.4271303 | 0.01 | 1.5686253 | 570 |
| ENSG00000263528 | *IKBKE* | 7.75 | 12 | 68025 | 0.5711993 | 0.041 | 1.2332013 | 574 |
| ENSG00000177606 | *JUN* | 9.285 | 10 | 49709 | 0.32360092 | 0.014 | 2.2259452 | 576 |
| ENSG00000198900 | *TOP1* | 9.285 | 10 | 49709 | 0.32360092 | 0.014 | 2.2259452 | 576 |
| ENSG00000137834 | *SMAD6* | 5.76 | 12 | 70777 | 0.710434 | 0.024 | 1.0866207 | 579 |
| ENSG00000111653 | *ING4* | 6.91 | 12 | 69105 | 0.62207466 | 0.031 | 1.1652129 | 579 |
| ENSG00000011275 | *RNF216* | 8.91 | 12 | 59454 | 0.34724462 | 0.022 | 1.7870173 | 581 |
| ENSG00000171206 | *TRIM8* | 9.495 | 12 | 60595 | 0.43240014 | 0.014 | 1.4411689 | 582 |
| ENSG00000187391 | *MAGI2* | 5.76 | 12 | 70784 | 0.7104409 | 0.03 | 1.0605493 | 583 |
| ENSG00000144802 | *NFKBIZ* | 7.92 | 12 | 67218 | 0.51772976 | 0.023 | 1.2481622 | 583 |
| ENSG00000176697 | *BDNF* | 5.76 | 12 | 70777 | 0.7104282 | 0.033 | 1.0690925 | 586 |
| ENSG00000138794 | *CASP6* | 5.76 | 12 | 70777 | 0.7104279 | 0.031 | 1.0713297 | 586 |
| ENSG00000106624 | *AEBP1* | 5.76 | 12 | 70785 | 0.7104448 | 0.045 | 1.0506582 | 588 |
| ENSG00000104408 | *EIF3E* | 9 | 12 | 63838 | 0.42702994 | 0.018 | 1.5188372 | 590 |
| ENSG00000131080 | *EDA2R* | 8.85 | 12 | 65024 | 0.43020087 | 0.023 | 1.4212871 | 591 |
| ENSG00000129988 | *LBP* | 5.76 | 12 | 70790 | 0.71043706 | 0.021 | 1.0498791 | 593 |
| ENSG00000157227 | *MMP14* | 5.76 | 12 | 70777 | 0.71042585 | 0.03 | 1.0598683 | 594 |
| ENSG00000049130 | *KITLG* | 7.92 | 12 | 67218 | 0.5177293 | 0.03 | 1.2322732 | 594 |
| ENSG00000173163 | *COMMD1* | 8.610001 | 12 | 59328 | 0.3733167 | 0.01 | 1.6696557 | 594 |
| ENSG00000177426 | *TGIF1* | 7.92 | 12 | 67220 | 0.5177254 | 0.024 | 1.245107 | 595 |
| ENSG00000100380 | *ST13* | 8.969999 | 12 | 63859 | 0.42679292 | 0.028 | 1.4982194 | 595 |
| ENSG00000153233 | *PTPRR* | 5.76 | 12 | 70777 | 0.7104353 | 0.041 | 1.0350901 | 598 |
| ENSG00000141753 | *IGFBP4* | 7.92 | 12 | 67218 | 0.5177292 | 0.031 | 1.2259469 | 598 |
| ENSG00000082074 | *FYB1* | 9.599999 | 12 | 60086 | 0.41582003 | 0.01 | 1.5812954 | 598 |
| ENSG00000275302 | *CCL4* | 7.92 | 12 | 67218 | 0.5177297 | 0.031 | 1.2243251 | 599 |
| ENSG00000126583 | *PRKCG* | 9 | 12 | 63769 | 0.42712083 | 0.01 | 1.4074908 | 599 |
| ENSG00000154229 | *PRKCA* | 9 | 12 | 63769 | 0.42712083 | 0.01 | 1.4074908 | 599 |
| ENSG00000166501 | *PRKCB* | 9 | 12 | 63769 | 0.42712083 | 0.01 | 1.4074908 | 599 |
| ENSG00000095015 | *MAP3K1* | 8.85 | 12 | 65059 | 0.43016466 | 0.035 | 1.3734448 | 602 |
| ENSG00000130159 | *ECSIT* | 8.85 | 12 | 65059 | 0.43016466 | 0.035 | 1.3734448 | 602 |
| ENSG00000213658 | *LAT* | 9.599999 | 12 | 60116 | 0.41576496 | 0.01 | 1.5740044 | 602 |
| ENSG00000150630 | *VEGFC* | 8.969999 | 12 | 63866 | 0.42678547 | 0.026 | 1.4386523 | 603 |
| ENSG00000106615 | *RHEB* | 9.57 | 12 | 60071 | 0.4158295 | 0.008 | 1.5662229 | 603 |
| ENSG00000225950 | *NTF4* | 5.76 | 12 | 70777 | 0.71043473 | 0.038 | 1.0128971 | 606 |
| ENSG00000146678 | *IGFBP1* | 5.76 | 12 | 70777 | 0.710435 | 0.025 | 1.0027324 | 609 |
| ENSG00000271503 | *CCL5* | 7.92 | 12 | 67218 | 0.5177286 | 0.036 | 1.2171644 | 609 |
| ENSG00000075213 | *SEMA3A* | 7.92 | 12 | 67218 | 0.51772624 | 0.035 | 1.2180241 | 609 |
| ENSG00000157933 | *SKI* | 7.92 | 12 | 67218 | 0.5177291 | 0.037 | 1.2151346 | 610 |
| ENSG00000183691 | *NOG* | 5.76 | 12 | 70778 | 0.71040905 | 0.047 | 1.0179756 | 611 |
| ENSG00000043462 | *LCP2* | 9.599999 | 12 | 60148 | 0.41575575 | 0.012 | 1.5598011 | 611 |
| ENSG00000051523 | *CYBA* | 8.985001 | 12 | 63770 | 0.4270362 | 0.024 | 1.3809861 | 612 |
| ENSG00000100365 | *NCF4* | 8.985001 | 12 | 63770 | 0.4270362 | 0.024 | 1.3809861 | 612 |
| ENSG00000116701 | *NCF2* | 8.985001 | 12 | 63770 | 0.4270362 | 0.024 | 1.3809861 | 612 |
| ENSG00000136238 | *RAC1* | 8.985001 | 12 | 63770 | 0.4270362 | 0.024 | 1.3809861 | 612 |
| ENSG00000142765 | *SYTL1* | 8.985001 | 12 | 63770 | 0.4270362 | 0.024 | 1.3809861 | 612 |
| ENSG00000158517 | *NCF1* | 8.985001 | 12 | 63770 | 0.4270362 | 0.024 | 1.3809861 | 612 |
| ENSG00000165168 | *CYBB* | 8.985001 | 12 | 63770 | 0.4270362 | 0.024 | 1.3809861 | 612 |
| ENSG00000113721 | *PDGFRB* | 5.76 | 12 | 70778 | 0.71040905 | 0.044 | 1.0032418 | 614 |
| ENSG00000197635 | *DPP4* | 7.92 | 12 | 67222 | 0.517722 | 0.031 | 1.2208937 | 614 |
| ENSG00000134853 | *PDGFRA* | 5.76 | 12 | 70778 | 0.7104186 | 0.05 | 1.0005448 | 616 |
| ENSG00000121653 | *MAPK8IP1* | 7.92 | 12 | 67218 | 0.5177258 | 0.03 | 1.2096031 | 618 |
| ENSG00000102524 | *TNFSF13B* | 8.91 | 11 | 37216 | 0.2654545 | 0.006 | 1.7353129 | 620 |
| ENSG00000213928 | *IRF9* | 7.92 | 12 | 67219 | 0.5177216 | 0.032 | 1.2152371 | 621 |
| ENSG00000121879 | *PIK3CA* | 7.92 | 12 | 67220 | 0.5177101 | 0.032 | 1.2247334 | 621 |
| ENSG00000078369 | *GNB1* | 7.98 | 9 | 31781 | 0.24260649 | 0.021 | 1.9368281 | 622 |
| ENSG00000115461 | *IGFBP5* | 7.92 | 12 | 67218 | 0.5177259 | 0.035 | 1.200358 | 623 |
| ENSG00000185499 | *MUC1* | 7.92 | 12 | 67218 | 0.5177259 | 0.035 | 1.2001833 | 623 |
| ENSG00000170581 | *STAT2* | 7.92 | 12 | 67219 | 0.517722 | 0.032 | 1.2059002 | 625 |
| ENSG00000142166 | *IFNAR1* | 7.92 | 12 | 67219 | 0.51772016 | 0.029 | 1.2098798 | 626 |
| ENSG00000158813 | *EDA* | 7.92 | 12 | 63549 | 0.4400373 | 0.005 | 1.2597772 | 628 |
| ENSG00000079385 | *CEACAM1* | 8.969999 | 12 | 63900 | 0.42671067 | 0.047 | 1.3360722 | 632 |
| ENSG00000187098 | *MITF* | 8.969999 | 12 | 63908 | 0.4267764 | 0.028 | 1.3022013 | 638 |
| ENSG00000129315 | *CCNT1* | 7.92 | 12 | 67222 | 0.51771617 | 0.036 | 1.1955749 | 641 |
| ENSG00000196083 | *IL1RAP* | 7.92 | 12 | 67221 | 0.5176931 | 0.042 | 1.1984386 | 649 |
| ENSG00000099942 | *CRKL* | 10 | 12 | 57637 | 0.335366 | 0.035 | 1.4703811 | 650 |
| ENSG00000107263 | *RAPGEF1* | 10 | 12 | 57637 | 0.335366 | 0.035 | 1.4703811 | 650 |
| ENSG00000187266 | *EPOR* | 10 | 12 | 57637 | 0.335366 | 0.035 | 1.4703811 | 650 |
| ENSG00000124762 | *CDKN1A* | 7.92 | 12 | 67219 | 0.5177215 | 0.05 | 1.1724962 | 652 |
| ENSG00000277632 | *CCL3* | 7.92 | 12 | 67219 | 0.51770365 | 0.039 | 1.1933305 | 652 |
| ENSG00000120868 | *APAF1* | 10 | 12 | 57686 | 0.33529928 | 0.048 | 1.5147556 | 652 |
| ENSG00000132906 | *CASP9* | 10 | 12 | 57686 | 0.33529928 | 0.048 | 1.5147556 | 652 |
| ENSG00000172115 | *CYCS* | 10 | 12 | 57686 | 0.33529928 | 0.048 | 1.5147556 | 652 |
| ENSG00000154310 | *TNIK* | 7.92 | 12 | 67220 | 0.51771235 | 0.036 | 1.175839 | 655 |
| ENSG00000131089 | *ARHGEF9* | 7.92 | 12 | 67236 | 0.5176874 | 0.045 | 1.1959542 | 657 |
| ENSG00000102547 | *CAB39L* | 9.835001 | 10 | 34919 | 0.24541828 | 0.033 | 1.6348159 | 657 |
| ENSG00000118046 | *STK11* | 9.835001 | 10 | 34919 | 0.24541828 | 0.033 | 1.6348159 | 657 |
| ENSG00000135932 | *CAB39* | 9.835001 | 10 | 34919 | 0.24541828 | 0.033 | 1.6348159 | 657 |
| ENSG00000266173 | *STRADA* | 9.835001 | 10 | 34919 | 0.24541828 | 0.033 | 1.6348159 | 657 |
| ENSG00000140299 | *BNIP2* | 7.92 | 12 | 67235 | 0.5176878 | 0.045 | 1.1950666 | 658 |
| ENSG00000197461 | *PDGFA* | 7.92 | 12 | 67233 | 0.5176887 | 0.04 | 1.192116 | 659 |
| ENSG00000127948 | *POR* | 9.795 | 10 | 43756 | 0.24303655 | 0.02 | 1.6571015 | 660 |
| ENSG00000109320 | *NFKB1* | 7.695 | 9 | 42750 | 0.24338467 | 0.012 | 1.5972216 | 662 |
| ENSG00000088387 | *DOCK9* | 7.92 | 12 | 67237 | 0.5176855 | 0.046 | 1.1904196 | 663 |
| ENSG00000085276 | *MECOM* | 7.92 | 12 | 67218 | 0.517708 | 0.047 | 1.1685082 | 665 |
| ENSG00000180228 | *PRKRA* | 7.92 | 12 | 67232 | 0.51767194 | 0.049 | 1.1877697 | 666 |
| ENSG00000017797 | *RALBP1* | 7.92 | 12 | 67239 | 0.51766765 | 0.048 | 1.1836513 | 669 |
| ENSG00000105974 | *CAV1* | 8.969999 | 12 | 63876 | 0.4267289 | 0.048 | 1.2320014 | 669 |
| ENSG00000140575 | *IQGAP1* | 7.92 | 12 | 67222 | 0.51771337 | 0.031 | 1.1524434 | 671 |
| ENSG00000171560 | *FGA* | 9.889999 | 10 | 43781 | 0.24304931 | 0.028 | 1.5700408 | 671 |
| ENSG00000166949 | *SMAD3* | 9 | 12 | 63756 | 0.42712975 | 0.012 | 1.1992165 | 676 |
| ENSG00000070831 | *CDC42* | 7.92 | 12 | 67220 | 0.5176907 | 0.046 | 1.1606978 | 678 |
| ENSG00000158092 | *NCK1* | 7.92 | 12 | 67219 | 0.51770896 | 0.042 | 1.0892324 | 691 |
| ENSG00000168067 | *MAP4K2* | 7.995 | 9 | 31776 | 0.24262741 | 0.017 | 1.4689363 | 694 |
| ENSG00000002330 | *BAD* | 9.93 | 10 | 43578 | 0.24308331 | 0.021 | 1.3219874 | 712 |
| ENSG00000111276 | *CDKN1B* | 7.9749994 | 9 | 31837 | 0.24258348 | 0.021 | 1.3293632 | 722 |
| ENSG00000123374 | *CDK2* | 7.9749994 | 9 | 31837 | 0.24258348 | 0.021 | 1.3293632 | 722 |
| ENSG00000105141 | *CASP14* | 9.9 | 12 | 59349 | 0.33481878 | 0.04 | 1.2508427 | 723 |
| ENSG00000196642 | *RABL6* | 8.91 | 12 | 64087 | 0.42654973 | 0.05 | 1.1622107 | 724 |
| ENSG00000163935 | *SFMBT1* | 9.9 | 12 | 59330 | 0.33482593 | 0.043 | 1.2230949 | 729 |
| ENSG00000157625 | *TAB3* | 7.92 | 12 | 64880 | 0.44248962 | 0.033 | 1.0469893 | 736 |
| ENSG00000178522 | *AMBN* | 9.9 | 12 | 59330 | 0.33482286 | 0.048 | 1.2187074 | 740 |
| ENSG00000204628 | *RACK1* | 9 | 12 | 63756 | 0.4271173 | 0.015 | 1.0624357 | 741 |
| ENSG00000088832 | *FKBP1A* | 8.969999 | 12 | 63860 | 0.42677236 | 0.035 | 1.0888858 | 741 |
| ENSG00000108622 | *ICAM2* | 9.889999 | 10 | 43734 | 0.24306636 | 0.016 | 1.2505156 | 741 |
| ENSG00000137802 | *MAPKBP1* | 9.9 | 12 | 59330 | 0.33482626 | 0.044 | 1.2007021 | 743 |
| ENSG00000111802 | *TDP2* | 9.225 | 12 | 55447 | 0.33578593 | 0.019 | 1.17512 | 750 |
| ENSG00000171522 | *PTGER4* | 9.9 | 12 | 59330 | 0.33482534 | 0.048 | 1.1962769 | 751 |
| ENSG00000067082 | *KLF6* | 8.91 | 12 | 64088 | 0.42654967 | 0.047 | 1.065588 | 756 |
| ENSG00000162733 | *DDR2* | 8.91 | 12 | 64102 | 0.42653918 | 0.042 | 1.0592271 | 763 |
| ENSG00000110651 | *CD81* | 9.9 | 12 | 59332 | 0.33482435 | 0.046 | 1.1714522 | 769 |
| ENSG00000109458 | *GAB1* | 8.969999 | 12 | 63910 | 0.42671496 | 0.049 | 1.026577 | 770 |
| ENSG00000019991 | *HGF* | 9.9 | 12 | 59334 | 0.33482125 | 0.049 | 1.1727505 | 774 |
| ENSG00000127914 | *AKAP9* | 9.9 | 12 | 59335 | 0.3348233 | 0.046 | 1.1691952 | 775 |
| ENSG00000091879 | *ANGPT2* | 9.91 | 10 | 43975 | 0.24301484 | 0.05 | 1.1873254 | 793 |
| ENSG00000103056 | *SMPD3* | 9.9 | 12 | 59336 | 0.33482343 | 0.033 | 1.1287276 | 796 |
| ENSG00000132024 | *CC2D1A* | 9.9 | 12 | 59333 | 0.3348239 | 0.037 | 1.1056803 | 797 |
| ENSG00000138668 | *HNRNPD* | 8.91 | 12 | 59459 | 0.35119116 | 0.04 | 1.0414926 | 800 |
| ENSG00000103490 | *PYCARD* | 9.9 | 12 | 59334 | 0.33482435 | 0.044 | 1.0793577 | 805 |
| ENSG00000264522 | *OTUD7B* | 8.61 | 12 | 59380 | 0.36004668 | 0.016 | 1.0028156 | 808 |
| ENSG00000141867 | *BRD4* | 9.455 | 10 | 48628 | 0.25022906 | 0.04 | 1.0719167 | 822 |
| ENSG00000143437 | *ARNT* | 9.780001 | 10 | 43703 | 0.24304433 | 0.028 | 1.1202061 | 823 |
| ENSG00000020426 | *MNAT1* | 9.38 | 10 | 48880 | 0.2501911 | 0.042 | 1.0640048 | 829 |
| ENSG00000134480 | *CCNH* | 9.38 | 10 | 48880 | 0.2501911 | 0.042 | 1.0640048 | 829 |
| ENSG00000184990 | *SIVA1* | 9.9 | 12 | 59367 | 0.33481124 | 0.042 | 1.0061295 | 840 |

Table S2: Controlling node genes that may regulate signalling activity on the NF-κB network

| ID | IBD importance | IBD Causal | IBD Correlative | IBD Mechanism | IBD Negative | IBD Preventative | IBD Prognosis | IBD Target | IBD GWAS prognosis | IBD GWAS susceptablity |
| --- | --- | --- | --- | --- | --- | --- | --- | --- | --- | --- |
| *TNF* | 10 | + | + |  | + | + | + | + | + | + |
| *TNFRSF11B* | 10 | + | + |  | + | + | + | + | + | + |
| *TNFRSF18* | 10 | + | + |  | + | + | + | + | + | + |
| *TNFRSF1A* | 10 | + | + |  | + | + | + | + | + | + |
| *TNFRSF4* | 10 | + | + |  | + | + | + | + | + | + |
| *TNFRSF6B* | 10 | + | + |  | + | + | + | + | + | + |
| *TNFRSF9* | 10 | + | + |  | + | + | + | + | + | + |
| *TNFSF11* | 10 | + | + |  | + | + | + | + | + | + |
| *TNFSF14* | 10 | + | + |  | + | + | + | + | + | + |
| *TNFSF15* | 10 | + | + |  | + | + | + | + | + | + |
| *TNFSF18* | 10 | + | + |  | + | + | + | + | + | + |
| *TNFSF4* | 10 | + | + |  | + | + | + | + | + | + |
| *IL10* | 10 | + | + | + | + | + | + | + |  | + |
| *IL13* | 10 | + | + | + | + | + | + | + |  | + |
| *IL15* | 10 | + | + | + | + | + | + | + |  | + |
| *IL15RA* | 10 | + | + | + | + | + | + | + |  | + |
| *IL17A* | 10 | + | + | + | + | + | + | + |  | + |
| *IL17F* | 10 | + | + | + | + | + | + | + |  | + |
| *IL18* | 10 | + | + | + | + | + | + | + |  | + |
| *IL18RAP* | 10 | + | + | + | + | + | + | + |  | + |
| *IL1A* | 10 | + | + | + | + | + | + | + |  | + |
| *IL1B* | 10 | + | + | + | + | + | + | + |  | + |
| *IL1R1* | 10 | + | + | + | + | + | + | + |  | + |
| *IL1RN* | 10 | + | + | + | + | + | + | + |  | + |
| *IL21* | 10 | + | + | + | + | + | + | + |  | + |
| *IL22* | 10 | + | + | + | + | + | + | + |  | + |
| *IL23R* | 10 | + | + | + | + | + | + | + |  | + |
| *IL25* | 10 | + | + | + | + | + | + | + |  | + |
| *IL26* | 10 | + | + | + | + | + | + | + |  | + |
| *IL27* | 10 | + | + | + | + | + | + | + |  | + |
| *IL31RA* | 10 | + | + | + | + | + | + | + |  | + |
| *IL4* | 10 | + | + | + | + | + | + | + |  | + |
| *IL5* | 10 | + | + | + | + | + | + | + |  | + |
| *IL6* | 10 | + | + | + | + | + | + | + |  | + |
| *IL6ST* | 10 | + | + | + | + | + | + | + |  | + |
| *CCL11* | 8 | + | + | + | + |  | + |  |  | + |
| *CCL13* | 8 | + | + | + | + |  | + |  |  | + |
| *CCL17* | 8 | + | + | + | + |  | + |  |  | + |
| *CCL2* | 8 | + | + | + | + |  | + |  |  | + |
| *CCL20* | 8 | + | + | + | + |  | + |  |  | + |
| *CCL26* | 8 | + | + | + | + |  | + |  |  | + |
| *CCR5* | 8 | + | + | + | + |  | + |  |  | + |
| *CCR6* | 8 | + | + | + | + |  | + |  |  | + |
| *CX3CL1* | 8 | + | + | + | + |  | + |  |  | + |
| *CXCL16* | 8 | + | + | + | + |  | + |  |  | + |
| *CXCL6* | 8 | + | + | + | + |  | + |  |  | + |
| *CXCL8* | 8 | + | + | + | + |  | + |  |  | + |
| *CXCR5* | 8 | + | + | + | + |  | + |  |  | + |
| *MAP3K8* | 5 | + | + |  |  |  |  |  | + | + |
| *MAPK1* | 5 | + | + |  |  |  |  |  | + | + |
| *MAPK3* | 5 | + | + |  |  |  |  |  | + | + |
| *MAPK8* | 5 | + | + |  |  |  |  |  | + | + |
| *SELP* | 5 | + | + |  |  |  |  |  | + | + |
| *SELPLG* | 5 | + | + |  |  |  |  |  | + | + |
| *STAT1* | 5 | + | + |  |  |  |  |  | + | + |
| *STAT3* | 5 | + | + |  |  |  |  |  | + | + |
| *STAT4* | 5 | + | + |  |  |  |  |  | + | + |
| *STAT6* | 5 | + | + |  |  |  |  |  | + | + |
| *IFNAR1* | 4 | + | + |  |  |  |  |  | + | + |
| *IFNG* | 4 | + | + |  |  |  |  |  | + | + |
| *IFNGR2* | 4 | + | + |  |  |  |  |  | + | + |
| *NOD1* | 4 | + | + |  |  |  |  |  | + | + |
| *NOD2* | 4 | + | + |  |  |  |  |  | + | + |
| *VEGFA* | 4 | + | + |  |  |  |  |  | + | + |
| *HSPA6* | 3 | + | + |  |  |  |  |  | + |  |
| *HSPD1* | 3 | + | + |  |  |  |  |  | + |  |
| *ICAM1* | 3 | + | + |  |  |  |  |  | + |  |
| *IL12A* | 3 | + | + |  |  |  |  |  | + |  |
| *IL12B* | 3 | + | + |  |  |  |  |  | + |  |
| *MMP1* | 3 | + | + |  |  |  |  |  | + |  |
| *MMP10* | 3 | + | + |  |  |  |  |  | + |  |
| *MMP2* | 3 | + | + |  |  |  |  |  | + |  |
| *MMP3* | 3 | + | + |  |  |  |  |  | + |  |
| *MMP7* | 3 | + | + |  |  |  |  |  | + |  |
| *MMP9* | 3 | + | + |  |  |  |  |  | + |  |
| *TGFB1* | 3 | + | + |  |  |  |  |  | + |  |
| *TLR2* | 3 | + | + |  |  |  |  |  | + |  |
| *TLR3* | 3 | + | + |  |  |  |  |  | + |  |
| *TLR4* | 3 | + | + |  |  |  |  |  | + |  |
| *ADAM15* | 3 |  | + |  |  |  | + |  |  | + |
| *ADAM30* | 3 |  | + |  |  |  | + |  |  | + |
| *ALPI* | 3 |  |  |  |  | + | + | + |  |  |
| *BMP7* | 3 | + |  | + |  |  | + |  |  |  |
| *EPO* | 3 |  |  |  |  | + |  | + |  | + |
| *CSF2* | 2 | + |  | + |  |  |  |  |  |  |
| *CTNNB1* | 2 |  | + |  |  |  |  |  |  |  |
| *DUSP1* | 2 |  |  |  |  |  |  |  |  | + |
| *DUSP22* | 2 |  |  |  |  |  |  |  |  | + |
| *FASLG* | 2 |  |  |  |  |  |  |  |  | + |
| *GPX2* | 2 | + | + |  |  |  |  |  |  |  |
| *GPX4* | 2 | + | + |  |  |  |  |  |  |  |
| *IGF1* | 2 | + | + |  |  |  |  |  |  |  |
| *IGSF6* | 2 | + | + |  |  |  |  |  |  |  |
| *IL2* | 2 | + | + |  |  |  |  |  |  |  |
| *IL2RA* | 2 | + | + |  |  |  |  |  |  |  |
| *KIT* | 2 | + | + |  |  |  |  |  |  |  |
| *MPO* | 2 | + | + |  |  |  |  |  |  |  |
| *NFKB2* | 2 | + | + |  |  |  |  |  |  |  |
| *NOS2* | 2 | + | + |  |  |  |  |  |  |  |
| *PRKCB* | 2 | + | + |  |  |  |  |  |  |  |
| *PTK2B* | 2 | + | + |  |  |  |  |  |  |  |
| *PTPN22* | 2 | + | + |  |  |  |  |  |  |  |
| *REL* | 2 | + | + |  |  |  |  |  |  |  |
| *RELA* | 2 | + | + |  |  |  |  |  |  |  |
| *SELE* | 2 | + | + |  |  |  |  |  |  |  |
| *SELL* | 2 | + | + |  |  |  |  |  |  |  |
| *SMAD3* | 2 | + | + |  |  |  |  |  |  |  |
| *TAB1* | 2 | + | + |  |  |  |  |  |  |  |
| *TPMT* | 2 | + | + |  |  |  |  |  |  |  |
| *ADCY3* | 1 |  |  |  |  |  |  |  |  | + |
| *ALDH2* | 1 |  |  |  |  |  |  |  |  | + |
| *CREB5* | 1 |  |  |  |  |  |  |  |  | + |
| *CTSZ* | 1 |  |  |  |  |  |  |  |  | + |
| *DAP* | 1 |  |  |  |  |  |  |  |  | + |
| *DNMT3B* | 1 |  |  |  |  |  |  |  |  | + |
| *EPHX2* | 1 |  |  |  |  |  |  |  |  | + |
| *F2* | 1 |  |  |  | + |  |  |  |  |  |
| *ABCB1* | 1 |  | + |  |  |  |  |  |  |  |
| *AOX1* | 1 |  | + |  |  |  |  |  |  |  |
| *CASP8* | 1 |  | + |  |  |  |  |  |  |  |
| *CDH1* | 1 |  | + |  |  |  |  |  |  |  |
| *CPB2* | 1 |  | + |  |  |  |  |  |  |  |
| *FGF7* | 1 | + |  |  |  |  |  |  |  |  |
| *GALC* | 1 | + |  |  |  |  |  |  |  |  |
| *GPR18* | 1 | + |  |  |  |  |  |  |  |  |
| *GPR183* | 1 | + |  |  |  |  |  |  |  |  |
| *GPR35* | 1 | + |  |  |  |  |  |  |  |  |
| *GPR65* | 1 | + |  |  |  |  |  |  |  |  |
| *HCK* | 1 | + |  |  |  |  |  |  |  |  |
| *IPMK* | 1 | + |  |  |  |  |  |  |  |  |
| *IRGM* | 1 | + |  |  |  |  |  |  |  |  |
| *ITPA* | 1 | + |  |  |  |  |  |  |  |  |
| *JAK2* | 1 | + |  |  |  |  |  |  |  |  |
| *KCNN3* | 1 | + |  |  |  |  |  |  |  |  |
| *KIF21B* | 1 | + |  |  |  |  |  |  |  |  |
| *LACC1* | 1 | + |  |  |  |  |  |  |  |  |
| *MST1* | 1 | + |  |  |  |  |  |  |  |  |
| *NDUFA13* | 1 | + |  |  |  |  |  |  |  |  |
| *PAK1* | 1 | + |  |  |  |  |  |  |  |  |
| *PFKFB4* | 1 | + |  |  |  |  |  |  |  |  |
| *PHACTR2* | 1 | + |  |  |  |  |  |  |  |  |
| *PLA2G4A* | 1 | + |  |  |  |  |  |  |  |  |
| *PLA2R1* | 1 | + |  |  |  |  |  |  |  |  |
| *PMM1* | 1 | + |  |  |  |  |  |  |  |  |
| *PRKAB1* | 1 | + |  |  |  |  |  |  |  |  |
| *PTGER4* | 1 | + |  |  |  |  |  |  |  |  |
| *PTGES* | 1 | + |  |  |  |  |  |  |  |  |
| *PTGS2* | 1 | + |  |  |  |  |  |  |  |  |
| *PTH* | 1 | + |  |  |  |  |  |  |  |  |
| *PTPRC* | 1 | + |  |  |  |  |  |  |  |  |
| *RASGRP1* | 1 | + |  |  |  |  |  |  |  |  |
| *RPS6KA2* | 1 | + |  |  |  |  |  |  |  |  |
| *RPS6KB1* | 1 | + |  |  |  |  |  |  |  |  |
| *S1PR2* | 1 | + |  |  |  |  |  |  |  |  |
| *TBXAS1* | 1 | + |  |  |  |  |  |  |  |  |
| *TST* | 1 | + |  |  |  |  |  |  |  |  |
| *TUBD1* | 1 | + |  |  |  |  |  |  |  |  |
| *TYK2* | 1 | + |  |  |  |  |  |  |  |  |
| *VDR* | 1 | + |  |  |  |  |  |  |  |  |

Table S3: IBD target genes, with source of data

| ID | Gene symbol | Score | FDR | Z-Score | Ranks sum |
| --- | --- | --- | --- | --- | --- |
| ENSG00000111537 | *IFNG* | 0.90934 | 0 | 25.515682 | 2 |
| ENSG00000148344 | *PTGES* | 0.90934 | 0 | 25.109789 | 20 |
| ENSG00000109471 | *IL2* | 0.90934 | 0 | 25.109789 | 24 |
| ENSG00000174175 | *SELP* | 0.90934 | 0 | 23.785404 | 34 |
| ENSG00000137462 | *TLR2* | 0.90934 | 0 | 17.603453 | 51 |
| ENSG00000136869 | *TLR4* | 0.90934 | 0 | 17.603453 | 53 |
| ENSG00000006210 | *CX3CL1* | 0.90934 | 0 | 25.109789 | 62 |
| ENSG00000130427 | *EPO* | 0.90934 | 0 | 16.325098 | 68 |
| ENSG00000026508 | *CD44* | 0.90934 | 0 | 15.6443615 | 73 |
| ENSG00000136689 | *IL1RN* | 0.90934 | 0 | 15.893072 | 79 |
| ENSG00000169429 | *CXCL8* | 0.90934 | 0 | 14.8982315 | 84 |
| ENSG00000115009 | *CCL20* | 0.90934 | 0 | 14.649521 | 88 |
| ENSG00000115353 | *TACR1* | 0.90934 | 0 | 14.649521 | 92 |
| ENSG00000113302 | *IL12B* | 0.90934 | 0 | 14.649521 | 94 |
| ENSG00000125538 | *IL1B* | 0.90934 | 0 | 9.924029 | 126 |
| ENSG00000100985 | *MMP9* | 0.90934 | 0 | 8.680479 | 132 |
| ENSG00000149269 | *PAK1* | 0.90934 | 0 | 8.444952 | 143 |
| ENSG00000166888 | *STAT6* | 0.90934 | 0 | 25.515682 | 153 |
| ENSG00000136244 | *IL6* | 0.90934 | 0 | 9.177899 | 155 |
| ENSG00000164400 | *CSF2* | 0.90934 | 0 | 6.7536426 | 174 |
| ENSG00000115008 | *IL1A* | 0.90934 | 0 | 6.7536426 | 180 |
| ENSG00000113580 | *NR3C1* | 0.90934 | 0 | 19.889582 | 192 |
| ENSG00000017427 | *IGF1* | 0.90934 | 0 | 6.512027 | 202 |
| ENSG00000112715 | *VEGFA* | 0.90934 | 0 | 14.649521 | 206 |
| ENSG00000164761 | *TNFRSF11B* | 0.90934 | 0 | 12.16242 | 208 |
| ENSG00000164136 | *IL15* | 0.90934 | 0 | 5.1354814 | 210 |
| ENSG00000118689 | *FOXO3* | 0.90934 | 0 | 8.444952 | 214 |
| ENSG00000175387 | *SMAD2* | 0.90934 | 0 | 10.42145 | 216 |
| ENSG00000108691 | *CCL2* | 0.90934 | 0 | 13.903391 | 218 |
| ENSG00000160791 | *CCR5* | 0.90934 | 0 | 22.46102 | 224 |
| ENSG00000039068 | *CDH1* | 0.90934 | 0 | 3.7350864 | 226 |
| ENSG00000112115 | *IL17A* | 0.90934 | 0 | 9.924029 | 233 |
| ENSG00000137752 | *CASP1* | 0.90934 | 0 | 3.4077682 | 235 |
| ENSG00000090339 | *ICAM1* | 0.90934 | 0 | 3.1813836 | 239 |
| ENSG00000102245 | *CD40LG* | 0.90934 | 0 | 3.1813836 | 243 |
| ENSG00000100030 | *MAPK1* | 0.90934 | 0 | 17.347782 | 251 |
| ENSG00000102882 | *MAPK3* | 0.90934 | 0 | 24.315157 | 260 |
| ENSG00000144381 | *HSPD1* | 0.90934 | 0 | 2.5022295 | 269 |
| ENSG00000115415 | *STAT1* | 0.90934 | 0 | 25.515682 | 273 |
| ENSG00000149968 | *MMP3* | 0.90934 | 0 | 2.4914422 | 275 |
| ENSG00000136634 | *IL10* | 0.90934 | 0 | 2.4914422 | 277 |
| ENSG00000113525 | *IL5* | 0.90934 | 0 | 2.2562785 | 288 |
| ENSG00000168036 | *CTNNB1* | 0.90934 | 0 | 25.515682 | 294 |
| ENSG00000232810 | *TNF* | 0.90934 | 0 | 2.2562785 | 296 |
| ENSG00000064012 | *CASP8* | 0.90934 | 0 | 24.050282 | 300 |
| ENSG00000067182 | *TNFRSF1A* | 0.90934 | 0 | 2.031885 | 322 |
| ENSG00000087245 | *MMP2* | 0.90934 | 0 | 8.92919 | 324 |
| ENSG00000138378 | *STAT4* | 0.90934 | 0 | 5.862493 | 326 |
| ENSG00000108094 | *CUL2* | 0.7465518 | 0.013 | 2.1583393 | 330 |
| ENSG00000005381 | *MPO* | 0.82925606 | 0.006 | 1.8166571 | 354 |
| ENSG00000105329 | *TGFB1* | 0.90934 | 0 | 3.1813836 | 429 |
| ENSG00000168610 | *STAT3* | 0.90934 | 0 | 13.903391 | 458 |
| ENSG00000107643 | *MAPK8* | 0.90934 | 0 | 19.889582 | 460 |
| ENSG00000164850 | *GPER1* | 0.909338 | 0 | 1.2649971 | 460 |
| ENSG00000118503 | *TNFAIP3* | 0.90934 | 0 | 1.1343101 | 528 |
| ENSG00000169194 | *IL13* | 0.90934 | 0 | 5.862493 | 530 |
| ENSG00000113520 | *IL4* | 0.90934 | 0 | 8.203337 | 532 |
| ENSG00000150782 | *IL18* | 0.90934 | 0 | 3.7350864 | 535 |
| ENSG00000007171 | *NOS2* | 0.90934 | 0 | 1.1343101 | 537 |
| ENSG00000115594 | *IL1R1* | 0.90934 | 0 | 2.2562785 | 539 |
| ENSG00000142208 | *AKT1* | 0.90934 | 0 | 15.6443615 | 541 |
| ENSG00000157404 | *KIT* | 0.90934 | 0.039 | 6.7536426 | 573 |

Table S4: List of IBD key-nodes, potential therapeutic targets for IBD

| Gene symbol | ID | Maximal radius | Score | Z-Score | FDR | Ranks sum | Target Significance | PASS Mechanisms |
| --- | --- | --- | --- | --- | --- | --- | --- | --- |
| *IFNG* | ENSG00000111537 | 3.6 | 0.90934 | 25.515682 | 0 | 2 | 6 | Interferon agonist,Interferon antagonist,Interferon gamma antagonist,Interferon inducer |
| *PTGES* | ENSG00000148344 | 3.6 | 0.90934 | 25.109789 | 0 | 20 | 1 | Prostaglandin-E synthase inhibitor,Prostaglandin-E2 synthase 1 inhibitor |
| *IL2* | ENSG00000109471 | 3.6 | 0.90934 | 25.109789 | 0 | 24 | 5 | Interleukin 2 agonist,Interleukin 2 antagonist,Interleukin agonist,Interleukin antagonist |
| *SELP* | ENSG00000174175 | 3.6 | 0.90934 | 23.785404 | 0 | 34 | 1 | Cell adhesion molecule inhibitor,Selectin P antagonist,Selectin antagonist |
| *TLR2* | ENSG00000137462 | 3.6 | 0.90934 | 17.603453 | 0 | 51 | 3 | Toll-Like receptor 2 agonist,Toll-Like receptor 2 antagonist,Toll-Like receptor agonist,Toll-Like receptor antagonist |
| *TLR4* | ENSG00000136869 | 3.6 | 0.90934 | 17.603453 | 0 | 53 | 1 | Toll-Like receptor 4 antagonist,Toll-Like receptor agonist,Toll-Like receptor antagonist |
| *CD44* | ENSG00000026508 | 3.6 | 0.90934 | 15.6443615 | 0 | 73 | 2 | Cell adhesion molecule inhibitor |
| *CXCL8* | ENSG00000169429 | 3.6 | 0.90934 | 14.8982315 | 0 | 84 | 6 | Interleukin 8 antagonist,Interleukin agonist,Interleukin antagonist |
| *IL12B* | ENSG00000113302 | 3.6 | 0.90934 | 14.649521 | 0 | 94 | 1 | Interleukin 12 agonist,Interleukin agonist,Interleukin antagonist |
| *IL1B* | ENSG00000125538 | 3.6 | 0.90934 | 9.924029 | 0 | 126 | 9 | Interleukin 1 antagonist,Interleukin 1b antagonist,Interleukin agonist,Interleukin antagonist |
| *MMP9* | ENSG00000100985 | 3.6 | 0.90934 | 8.680479 | 0 | 132 | 4 | Collagenase inhibitor,Gelatinase inhibitor,Matrix metalloproteinase inhibitor,Metalloproteinase inhibitor,Metalloproteinase-9 inhibitor |
| *PAK1* | ENSG00000149269 | 3.6 | 0.90934 | 8.444952 | 0 | 143 | 10 | Protein-serine-threonine kinase inhibitor,p21-activated kinase 1 inhibitor,p21-activated kinase inhibitor |
| *STAT6* | ENSG00000166888 | 3.6 | 0.90934 | 25.515682 | 0 | 153 | 4 | Transcription factor STAT inhibitor,Transcription factor STAT6 inhibitor |
| *IL6* | ENSG00000136244 | 3.6 | 0.90934 | 9.177899 | 0 | 155 | 5 | Interleukin 6 antagonist,Interleukin agonist,Interleukin antagonist |
| *CSF2* | ENSG00000164400 | 3.6 | 0.90934 | 6.7536426 | 0 | 174 | 1 | Colony stimulating factor agonist,Colony stimulating factor antagonist,Granulocyte macrophage colony stimulating factor agonist,Granulocyte macrophage colony stimulating factor antagonist,Hydrolase inhibitor |
| *IL1A* | ENSG00000115008 | 3.6 | 0.90934 | 6.7536426 | 0 | 180 | 1 | Interleukin 1 antagonist,Interleukin 1a antagonist,Interleukin agonist,Interleukin antagonist |
| *IGF1* | ENSG00000017427 | 3.6 | 0.90934 | 6.512027 | 0 | 202 | 2 | Insulin like growth factor 1 agonist |
| *VEGFA* | ENSG00000112715 | 3.6 | 0.90934 | 14.649521 | 0 | 206 | 4 | Endothelial growth factor antagonist |
| *IL15* | ENSG00000164136 | 3.6 | 0.90934 | 5.1354814 | 0 | 210 | 5 | Interleukin agonist,Interleukin antagonist |
| *CCR5* | ENSG00000160791 | 3.6 | 0.90934 | 22.46102 | 0 | 224 | 1 | CC chemokine 5 receptor agonist,CC chemokine 5 receptor antagonist,CC chemokine receptor agonist,CC chemokine receptor antagonist,Chemokine receptor agonist,Chemokine receptor antagonist |
| *CDH1* | ENSG00000039068 | 3.6 | 0.90934 | 3.7350864 | 0 | 226 | 1 | Cadherin antagonist,Cell adhesion molecule inhibitor |
| *IL17A* | ENSG00000112115 | 3.6 | 0.90934 | 9.924029 | 0 | 233 | 4 | Interleukin agonist,Interleukin antagonist |
| *CASP1* | ENSG00000137752 | 3.6 | 0.90934 | 3.4077682 | 0 | 235 | 3 | Interleukin 1 beta converting enzyme inhibitor |
| *ICAM1* | ENSG00000090339 | 3.6 | 0.90934 | 3.1813836 | 0 | 239 | 4 | Cell adhesion molecule inhibitor,ICAM 1 antagonist |
| *MAPK1* | ENSG00000100030 | 3.6 | 0.90934 | 17.347782 | 0 | 251 | 5 | MAP kinase 1 inhibitor,MAP kinase inhibitor,MAP kinase stimulant |
| *MAPK3* | ENSG00000102882 | 3.6 | 0.90934 | 24.315157 | 0 | 260 | 4 | MAP kinase 3 inhibitor,MAP kinase inhibitor,MAP kinase stimulant |
| *HSPD1* | ENSG00000144381 | 3.6 | 0.90934 | 2.5022295 | 0 | 269 | 3 | Chaperonin ATPase inhibitor |
| *STAT1* | ENSG00000115415 | 3.6 | 0.90934 | 25.515682 | 0 | 273 | 3 | Transcription factor STAT inhibitor,Transcription factor STAT1 inhibitor |
| *MMP3* | ENSG00000149968 | 3.6 | 0.90934 | 2.4914422 | 0 | 275 | 3 | Matrix metalloproteinase 3 (membrane-type) inhibitor,Matrix metalloproteinase inhibitor,Metalloproteinase inhibitor,Metalloproteinase-3 inhibitor |
| *IL10* | ENSG00000136634 | 3.6 | 0.90934 | 2.4914422 | 0 | 277 | 6 | Interleukin 10 agonist,Interleukin 10 antagonist,Interleukin agonist,Interleukin antagonist |
| *IL5* | ENSG00000113525 | 3.6 | 0.90934 | 2.2562785 | 0 | 288 | 1 | Interleukin 5 antagonist,Interleukin agonist,Interleukin antagonist |
| *CTNNB1* | ENSG00000168036 | 3.6 | 0.90934 | 25.515682 | 0 | 294 | 2 | Catenin beta inhibitor |
| *TNF* | ENSG00000232810 | 3.6 | 0.90934 | 2.2562785 | 0 | 296 | 10 | Tumour necrosis factor alpha antagonist,Tumour necrosis factor alpha release inhibitor,Tumour necrosis factor antagonist |
| *CASP8* | ENSG00000064012 | 3.6 | 0.90934 | 24.050282 | 0 | 300 | 1 | Caspase 8 inhibitor,Caspase 8 stimulant |
| *MMP2* | ENSG00000087245 | 3.6 | 0.90934 | 8.92919 | 0 | 324 | 3 | Collagenase inhibitor,Gelatinase inhibitor,Matrix metalloproteinase 2 (membrane-type) inhibitor,Matrix metalloproteinase inhibitor,Metalloproteinase inhibitor,Metalloproteinase-2 inhibitor |
| *STAT4* | ENSG00000138378 | 3.6 | 0.90934 | 5.862493 | 0 | 326 | 1 | Transcription factor STAT inhibitor |
| *MPO* | ENSG00000005381 | 5.3999996 | 0.82925606 | 1.8166571 | 0.006 | 354 | 3 | Myeloperoxidase inhibitor,Peroxidase inhibitor |
| *TGFB1* | ENSG00000105329 | 3.6 | 0.90934 | 3.1813836 | 0 | 429 | 2 | Transforming growth factor agonist,Transforming growth factor antagonist,Transforming growth factor beta 1 agonist,Transforming growth factor beta 1 antagonist,Transforming growth factor beta agonist,Transforming growth factor beta antagonist |
| *STAT3* | ENSG00000168610 | 3.6 | 0.90934 | 13.903391 | 0 | 458 | 6 | Transcription factor STAT inhibitor,Transcription factor STAT3 inhibitor |
| *MAPK8* | ENSG00000107643 | 3.6 | 0.90934 | 19.889582 | 0 | 460 | 2 | JNK mitogen-activated protein kinase inhibitor,MAP kinase 8 inhibitor,MAP kinase inhibitor,MAP kinase stimulant |
| *IL13* | ENSG00000169194 | 3.6 | 0.90934 | 5.862493 | 0 | 530 | 1 | Interleukin agonist,Interleukin antagonist |
| *IL4* | ENSG00000113520 | 3.6 | 0.90934 | 8.203337 | 0 | 532 | 1 | Interleukin 4 antagonist,Interleukin agonist,Interleukin antagonist |
| *IL18* | ENSG00000150782 | 3.6 | 0.90934 | 3.7350864 | 0 | 535 | 2 | Interleukin agonist,Interleukin antagonist |
| *NOS2* | ENSG00000007171 | 3.6 | 0.90934 | 1.1343101 | 0 | 537 | 1 | Inducible nitric oxide synthase inhibitor,Inducible nitric-oxide synthase inhibitor,Nitric-oxide synthase inhibitor,Nitric-oxide synthase stimulant |
| *AKT1* | ENSG00000142208 | 3.6 | 0.90934 | 15.6443615 | 0 | 541 | 2 | Protein kinase (PKA_comma_ PKC_comma_ AKT_comma_ GRK_comma_ AGC-related_comma_ RSK_comma_ DBF2_comma_ SGK) inhibitor,Protein kinase B alpha inhibitor,Protein kinase B inhibitor,Protein kinase B stimulant,Protein-serine-threonine kinase inhibitor |
| *KIT* | ENSG00000157404 | 6.91 | 0.90934 | 6.7536426 | 0.039 | 573 | 2 | Protein-tyrosine kinase (PTK_comma_ not ETK_comma_ WZC) inhibitor,Proto-oncogene tyrosine-protein kinase Kit inhibitor,Tyrosine kinase stimulant,Tyrosine-protein kinase receptor antagonist |

Table S5: IBD key-nodes and their associated PASS activities

| Gene symbol | ID | Site model ID | Selected in #ChIP-seq data sets |
| --- | --- | --- | --- |
| *BACH1* | ENSG00000156273 | V$BACH1_01 | 9 |
| *BACH2* | ENSG00000112182 | V$BACH2_01 | 10 |
| *CBFB* | ENSG00000067955 | V$AML_Q6,V$PEBP_Q6 | 9 |
| *DEAF1* | ENSG00000177030 | V$DEAF1_02 | 10 |
| *EBF1* | ENSG00000164330 | V$COE1_Q6 | 9 |
| *ELF1* | ENSG00000120690 | V$ELF1_Q6 | 10 |
| *ELK1* | ENSG00000126767 | V$ELK1_01,V$ELK1_02 | 10 |
| *ETS1* | ENSG00000134954 | V$CETS1P54_03,V$ETS1_B,V$ETS_B | 10 |
| *ETS2* | ENSG00000157557 | V$ETS_B | 10 |
| *FOS* | ENSG00000170345 | V$AP1_01 | 10 |
| *FOSB* | ENSG00000125740 | V$AP1_01 | 9 |
| *FOSL1* | ENSG00000175592 | V$AP1_01,V$FRA1_Q5,V$FRA1_Q6 | 10 |
| *GABPA* | ENSG00000154727 | V$GADP_01 | 9 |
| *GABPB1* | ENSG00000104064 | V$GABPBETA_Q3,V$GADP_01 | 10 |
| *IRF1* | ENSG00000125347 | V$IRF1_01,V$IRF1_Q6,V$IRF1_Q6_01,V$IRF_Q6,V$IRF_Q6_01 | 10 |
| *IRF2* | ENSG00000168310 | V$IRF2_01,V$IRF_Q6,V$IRF_Q6_01 | 10 |
| *IRF3* | ENSG00000126456 | V$IRF_Q6,V$IRF_Q6_01 | 9 |
| *IRF4* | ENSG00000137265 | V$IRF_Q6,V$IRF_Q6_01 | 10 |
| *IRF5* | ENSG00000128604 | V$IRF_Q6,V$IRF_Q6_01 | 10 |
| *IRF6* | ENSG00000117595 | V$IRF_Q6 | 9 |
| *IRF7* | ENSG00000185507 | V$IRF_Q6,V$IRF_Q6_01 | 10 |
| *IRF8* | ENSG00000140968 | V$ICSBP_Q6,V$IRF8_Q6,V$IRF_Q6,V$IRF_Q6_01 | 10 |
| *IRF9* | ENSG00000213928 | V$IRF_Q6_01,V$ISRE_01 | 9 |
| *JUN* | ENSG00000177606 | V$AP1_01 | 10 |
| *JUNB* | ENSG00000171223 | V$AP1_01,V$JUNB_Q6 | 9 |
| *JUND* | ENSG00000130522 | V$AP1_01,V$JUND_Q6 | 9 |
| *MAFK* | ENSG00000198517 | V$MAFK_Q3,V$NFE2_01,V$NFE2_Q6 | 10 |
| *NFE2* | ENSG00000123405 | V$NFE2_01,V$NFE2_Q6 | 9 |
| *NFKB1* | ENSG00000109320 | V$NFKAPPAB50_01,V$NFKAPPAB_01,V$NFKB_C,V$NFKB_Q6,V$NFKB_Q6_01,V$P50P50_Q3,V$P50RELAP65_Q5_01,V$P50_Q6,V$RELBP52_01 | 10 |
| *NFKB2* | ENSG00000077150 | V$NFKB_Q6_01,V$RELBP52_01 | 10 |
| *REL* | ENSG00000162924 | V$CREL_01,V$CREL_Q6 | 10 |
| *RELA* | ENSG00000173039 | V$NFKAPPAB65_01,V$NFKAPPAB_01,V$NFKB_Q6_01,V$P50RELAP65_Q5_01,V$RELA_Q6,V$RELBP52_01 | 10 |
| *RELB* | ENSG00000104856 | V$RELBP52_01 | 10 |
| *RUNX1* | ENSG00000159216 | V$AML1_Q4,V$AML1_Q6,V$AML_Q6,V$PEBP_Q6 | 10 |
| *RUNX2* | ENSG00000124813 | V$AML3_Q6,V$AML_Q6,V$OSF2_Q6,V$PEBP_Q6 | 10 |
| *RUNX3* | ENSG00000020633 | V$AML2_01,V$AML_Q6,V$PEBP_Q6 | 10 |
| *SPI1* | ENSG00000066336 | V$PU1_Q4,V$PU1_Q6,V$SPI1_01 | 10 |
| *SPIB* | ENSG00000269404 | V$SPIB_01 | 10 |

Table S7: NF-κB co-operating transcription factors identified by MEALR

1. Koschmann J, Bhar A, Stegmaier P, et al. "Upstream Analysis": An Integrated Promoter-Pathway Analysis Approach to Causal Interpretation of Microarray Data. *Microarrays (Basel)* 2015;4(2):270-86. doi: 10.3390/microarrays4020270 [published Online First: 2015/01/01]

2. Stegmaier P, Kel A, Wingender E, et al. A discriminative approach for unsupervised clustering of DNA sequence motifs. *PLoS computational biology* 2013;9(3):e1002958. doi: 10.1371/journal.pcbi.1002958 [published Online First: 2013/04/05]
